# Most published human disease RNA-seq cannot be uniformly reanalyzed: a population-scale audit of reprocessability in the Gene Expression Omnibus

**DOI:** 10.64898/2026.08.05.742891

**Authors:** Abdullah Bin Murad

## Abstract

**Background:** Reuse of archived transcriptomic data underpins a large and growing share of published genomics. Because differences in upstream processing confound cross-study comparison, uniform reprocessing compendia — recount3, ARCHS4, DEE2, refine.bio, Expression Atlas — are widely treated as the remedy, and their availability is routinely assumed at the point of study design. Whether that remedy is actually obtainable for the population of published disease RNA-seq has not been measured. Prior audits have characterised metadata completeness and deposition rates, but none has quantified, across the published population, what fraction of studies can be uniformly reprocessed or where in the path from publication to comparable counts that capability is lost.

**Results:** We enumerated 1,124 MeSH disease descriptors exhaustively, retrieved 16,820 human RNA-seq series from the Gene Expression Omnibus, and audited the 3,631 bulk, Illumina-platform series of at least 25 samples under three independently pre-registered, tool-enforced analysis plans. Raw reads were publicly available for 94.1% of series under a dual-route evidence standard, but only 46.7% appeared in any uniform reprocessing compendium (bounds 46.7–57.2%) and only 27.3% were usably covered at a 90% run threshold (bounds 27.3–38.8%). Of the 3,418 series whose reads are public, 991 were usably covered, leaving **71.0% of read-public series reprocessed by nothing usable**. Presence overstated usability: DEE2 was present for 30.2% of series but usable for 3.4%. Design attrition was independent and severe — 24.7% met bulk primary-tissue case–control criteria, 7.4% additionally reached a minimum replication threshold counted on sample accessions, and 4.1% did so counted on distinct donors. Among the 199 series where donor identity resolves, 4.1% pass the replication criterion on donors against 15.1% on accessions, a 3.73-fold difference; across the census frame, accessions exceeded distinct donors by 2.65-fold (Manski bounds 1.09–10.77, Imbens–Manski 95% CI 1.07–11.88). Independently, 36.4% of series-to-disease attributions produced by a conventional keyword query were refuted by the curated MeSH headings of the series’ own linked publication.

**Conclusions:** Uniform reanalysis of published human disease RNA-seq is unavailable for most studies in the population audited, and the binding constraint is usable coverage rather than deposition of raw reads. The loss occurs at several independent layers with different remedies, and the coverage layer — unlike the others — is one that resource maintainers can act on. Automated retrieval further over-counts eligible studies, both by admitting designs outside scope and by assigning studies to diseases their publications do not support.

## 1. Background

Public transcriptomic archives have become primary research infrastructure. The Gene Expression Omnibus (GEO) holds well over 200,000 studies, and a substantial secondary literature now rests on reanalysing them [1]. Cross-study comparison, however, is confounded by upstream processing: two studies of the same disease quantified with different aligners, annotation releases and filtering conventions are not directly comparable, and differences attributable to pipeline choice can be mistaken for biology. Recent work confirms that preprocessing heterogeneity degrades cross-study inference, and that differential-expression results are themselves fragile at the cohort sizes typical of archived disease studies [14,15].

The standard remedy is uniform reprocessing. Compendia including recount3 [2], ARCHS4 [3], DEE2 [4], refine.bio and Expression Atlas [23] re-quantify archived raw reads through a single pipeline, so that counts from different studies become comparable. A cross-cohort analysis that plans to use uniformly reprocessed counts presupposes that such counts exist for the cohorts of interest. That presupposition has not been tested at population scale.

Existing audits of public transcriptomic data address adjacent questions. Metadata completeness in GEO has been characterised in detail, with over a quarter of critical metadata fields omitted across 253 studies and more than 164,000 samples [5]. Sample-level attribute reporting is sparse — sex was reported for 7.3% of nearly 50,000 human SRA RNA-seq samples [6] — and deposition rates during the microarray-to-sequencing transition have been estimated [7]. Curation projects have annotated subsets of GEO to enable reuse [8,9]. A standards-oriented perspective on genomic reusability has been articulated qualitatively [30]. None of this work measures whether the uniform reprocessing that reuse-based studies depend on is obtainable, nor where that capability is lost. To our knowledge no published work quantifies coverage or overlap among uniform reprocessing compendia; the only existing comparison is a descriptive sample-count table in the recount3 paper [2].

Here we report a population-scale audit of reprocessability. We enumerate MeSH disease descriptors exhaustively, retrieve the corresponding GEO series, and measure a layered cascade from retrieval to usable uniform coverage, distinguishing at each step between what is absent, what is present but unusable, and what cannot be assessed. We additionally verify whether each series is genuinely about the disease under which it was retrieved, and quantify two systematic biases in automated screening that inflate apparent study availability. Retrieval is reported against PRISMA 2020 flow conventions [13] with search reporting following the PRISMA-S extension [31], adapted from a literature search to a dataset search.

## 2. Results

### 2.1 Retrieval, screening and the audited population

Exhaustive enumeration over the 1,124 frozen MeSH disease descriptors returned 16,820 unique GEO series. Four screening criteria, applied before any coverage was measured, defined the audited population of 3,631 series (Table 1, Fig. 1a). The dominant criterion was a minimum of 25 samples per series, which removed 9,967 series — more than all other criteria combined. This threshold is a scope decision rather than a finding: a series with fewer than 25 samples cannot in principle satisfy the replication criterion applied at layer L1b, and retaining such series would have made the design layer uninformative. It is recorded in a configuration file that is explicitly declared post hoc and carries no freeze, and every rate reported below is conditional on it. We measure that conditioning directly rather than merely disclosing it (Fig. 5), and the result does not favour our headline; it is reported in full below.

**Table 1.** Retrieval and screening attrition. Criteria were applied in the order shown; each count is the number removed among series surviving the preceding criteria.

| Step | Removed | Remaining | Basis |
| --- | --- | --- | --- |
| Unique GEO series returned by the enumerated disease queries | — | **16,820** | Exhaustive over 1,124 frozen MeSH descriptors |
| Single-cell or single-nucleus protocol | 2,798 | 14,022 | Platform and study-text screen |
| Microarray platform | 174 | 13,848 | Platform screen |
| Non-Illumina sequencing platform | 250 | 13,598 | Platform screen |
| **Fewer than 25 samples** | **9,967** | 3,631 | Post-hoc declared scope threshold; see text |
| **Audited population (L0)** | — | **3,631** | Carried into all subsequent layers |

*Reconciles exactly with the PRISMA 2020 flow (Additional file 6). The screening rule is implemented in a committed script that asserts it reproduces this population exactly.*

### 2.2 A layered cascade from retrieval to usable coverage

Layers were measured against layer-specific denominators, because they have different data dependencies: design eligibility requires the sample table, count-matrix deposition requires the supplementary file listing, and compendium coverage requires querying three external resources. Reporting a single pooled denominator would present three distinct evidentiary bases as one. Table 2 gives each layer with its denominator; Fig. 1b shows the cascade.

**Table 2.** The reprocessability cascade, with layer-specific denominators.

| Layer | Definition | Rate | Denominator | Basis |
| --- | --- | --- | --- | --- |
| L0 | Series in the audited population (Table 1) | 3,631 | — | Exhaustive over 1,124 frozen descriptors |
| L1a | Qualifying bulk primary-tissue case–control design | 24.7% | weighted | Census stratum 850 + sampled stratum 598 |
| L1b | L1a and $\geq 15$ case, $\geq 10$ control samples | 7.4% | weighted | Counted on sample accessions |
| L1c | L1a and $\geq 15$ case, $\geq 10$ control donors | 4.1% | n = 199 | Donor-resolvable stratum only; conditional |
| L2 | Gene-level raw count matrix deposited | 40.1% | 1,084 | GEO-verified supplementary listing |
| L2' | Normalised values only, no raw counts | 8.8% | 1,084 | GEO-verified supplementary listing |
| L3 | Raw reads publicly available | 94.1% | 3,631 | Dual-route adjudicated; route A alone 93.5% |
| L4 | Present in $\geq 1$ uniform reprocessing compendium | 46.7% | 3,631 | Bounds 46.7–57.2% |
| L5 | Usably covered ($\geq 90\%$ of runs, QC-passing) | 27.3% | 3,631 | Bounds 27.3–38.8% |

*Denominators differ by layer because the layers have different data dependencies. L4 and L5 are reported as bounded intervals: 421 series have at least one compendium lookup that could not be performed, and an unassessable lookup is counted as a third outcome rather than as an absence. The lower bound is the confirmed count; the upper bound additionally counts every unassessable series as covered. Of the 3,418 series reaching L3, 991 (29.0%) reach L5.*

### 2.3 Reads are deposited far more often than they are reprocessed

Raw reads were publicly available for 94.1% of the audited population under a standard requiring agreement between an independent programmatic route and a direct reading of the primary GEO record. Deposition is therefore not the binding constraint. Coverage is: 46.7% of series appeared in at least one uniform compendium and 27.3% were usably covered. Restricting attention to the 3,418 series whose reads are public, 991 were usably covered — so **71.0% of series whose raw data are available to anyone are reprocessed by nothing usable** (Fig. 1b).

Completing the second resolution route materially changed this layer and is reported as a finding rather than a correction. Of the 237 series the programmatic route could not resolve, **24 (10.1%) are contradicted by the primary record** — their reads are public and the programmatic route missed them. Two of these, GSE79210 and GSE84023, are usably covered by DEE2, which requires that their reads were processed. Read availability measured through a single programmatic route is therefore an underestimate, and the single-route figure of 93.5% is retained alongside the adjudicated 94.1% rather than discarded.

The remaining series are reported in categories that are never pooled, because they describe different behaviours with different remedies (Fig. 3b): 28 series explicitly withhold raw data, 50 direct the reader to a controlled-access repository, and 96 show no evidence of deposition and no withholding statement. A further 39 GEO records carry no raw-data status line at all, which matters because the pre-registered evidence standard for this layer rests on that line.

**Fig. 1.**
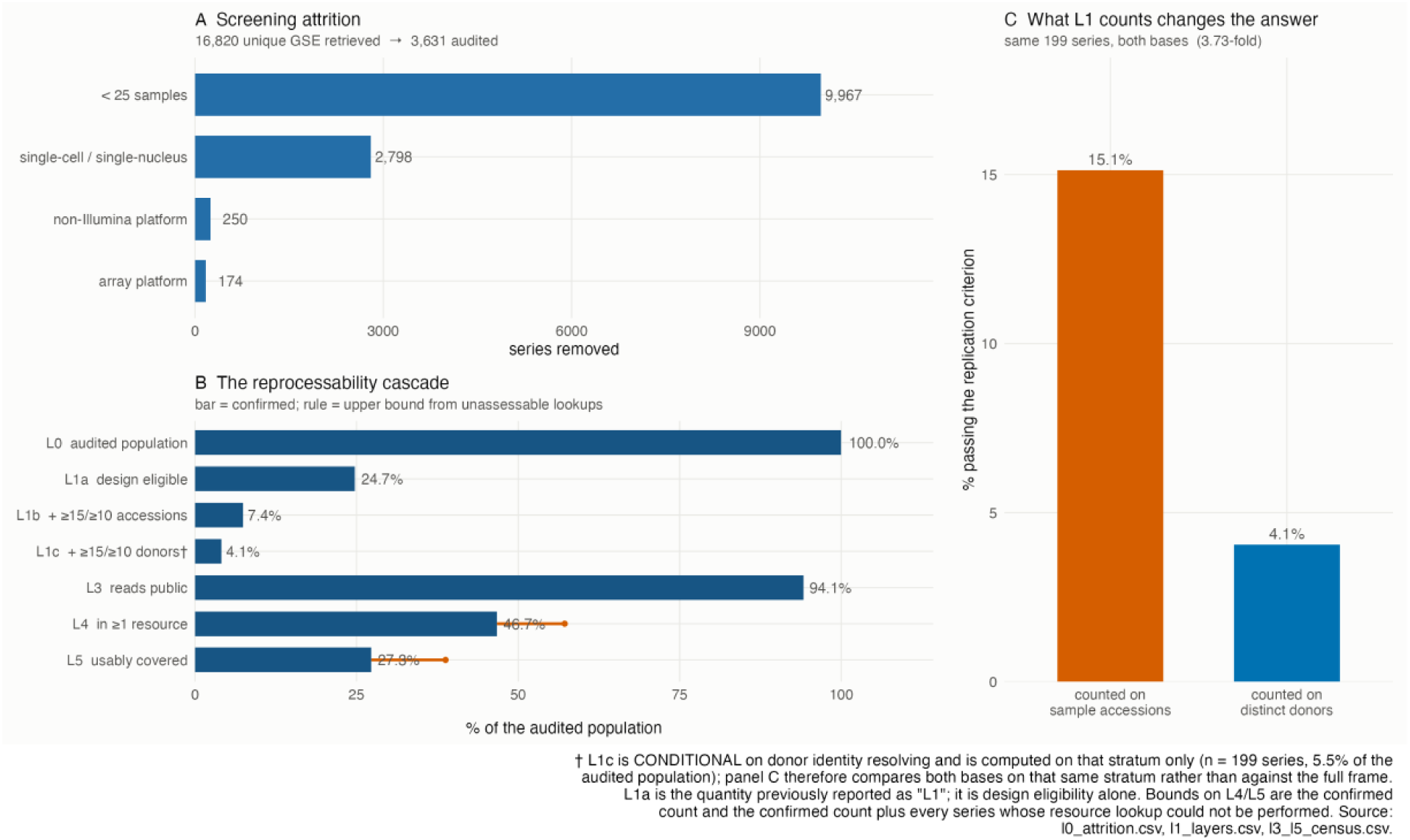
The audited population and the reprocessability cascade. (a) Screening attrition from 16,820 retrieved series to the 3,631 audited; the ≥25-sample threshold removes more series than all other criteria combined. (b) Each layer as a percentage of its own denominator; bars are confirmed counts and rules extend to the upper bound implied by unassessable lookups. (c) The replication criterion evaluated two ways on the same 199 series in which donor identity resolves: 15.1% pass when case and control counts are taken from sample accessions against 4.1% when taken from distinct donors, a 3.73-fold difference. L1a is the quantity reported in earlier drafts of this work as “L1”; it is design eligibility alone.

### 2.4 Presence in a compendium does not imply usable coverage

Coverage differed markedly between resources, and the gap between presence and usability differed more (Table 3, Fig. 2a). ARCHS4 was the most complete resource at 33.8% present and 23.4% usable. recount3 reached 11.8% present and 8.7% usable, consistent with its human SRA collection reflecting a processing snapshot that predates much of the retrieved literature. DEE2 was present for 30.2% of series but usable for only 3.4% once its own per-run quality flags were honoured — a nine-fold collapse, and the clearest demonstration that a presence/absence coverage statistic materially overstates what a reanalyst can use.

Coverage is computed as the fraction of a series’ own sequencing runs present in the resource, which is bounded in [0,1] by construction (Fig. 2b). Unassessable lookups — chiefly where the series-to-project mapping could not be resolved — are recorded as a third outcome and never as absences, which is why L4 and L5 are intervals rather than points. 421 series carry at least one such lookup.

**Table 3.** Coverage by uniform reprocessing compendium, as a percentage of the 3,631 audited series.

| Resource | Present | Usable | Unassessable | Notes |
| --- | --- | --- | --- | --- |
| ARCHS4 v2.4 | 33.8% | 23.4% | 2 | Series-keyed; pseudoalignment estimated counts. Most complete resource. |
| recount3 | 11.8% | 8.7% | 421 | Project-keyed; snapshot predates much of the retrieved literature. |
| DEE2 | 30.2% | 3.4% | 381 | Nine-fold collapse once per-run QC flags are honoured. |
| **Any resource** | **46.7%** | **27.3%** | **421** | Union. Bounds 46.7–57.2% and 27.3–38.8%. |

Coverage is snapshot-dependent and will change as resources ingest new studies. Access dates are given in Methods. Usable coverage requires $\geq 90\%$ of the series' runs to be present and, where the resource publishes per-run quality flags, quality-passing.

**Fig. 2.**
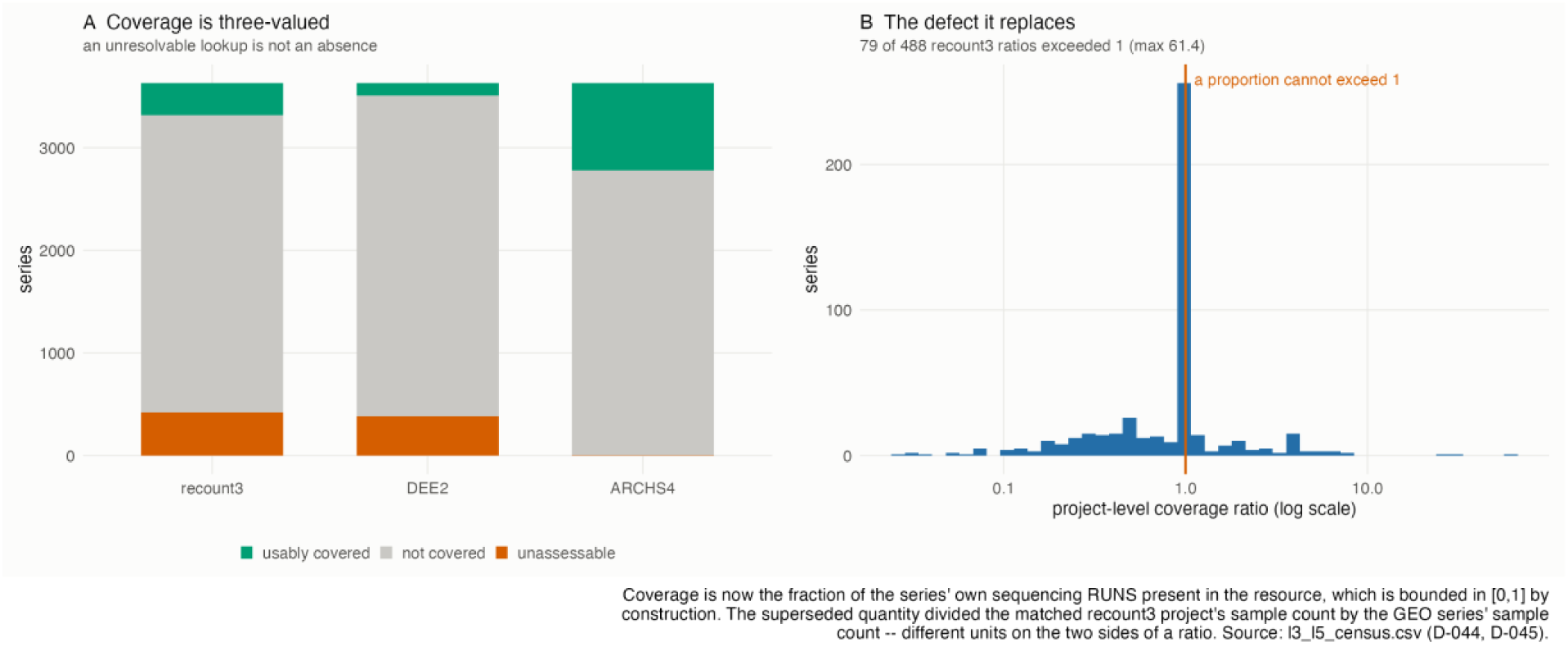
Coverage is three-valued and bounded. (a) Per-resource outcomes across the audited population: usably covered, not covered, and unassessable. An unresolvable lookup is not an absence, and treating it as one would understate coverage. (b) Distribution of the superseded project-level coverage ratio, which divided a matched project’s sample count by the series’ sample count — different units on the two sides of a ratio — and exceeded 1 for 79 of 488 series, to a maximum of 61.4. The run-level intersection used here cannot exceed 1.

### 2.5 Design attrition, and what the replication criterion is counted on

Of series with a verifiable sample table, 24.7% satisfied the design criteria — human bulk RNA-seq of primary tissue, excluding cultured or isolated-cell material, experimental perturbation arms and model-organism material (L1a). Adding a minimum of 15 case and 10 control samples for the descriptor under which the series was retrieved reduced this to 7.4% (L1b). Of series with a verifiable supplementary file listing, 40.1% deposited a gene-level raw count matrix and a further 8.8% deposited normalised values only, the latter unusable for count-based differential expression.

We report the replication criterion twice, because what it is counted on changes the answer. Sample accessions are widely treated as independent replicates; they are not. Restricting to the 199 series in which donor identity resolves from sample metadata, **4.1% pass the replication criterion when case and control counts are taken from distinct donors against 15.1% when taken from sample accessions** — a 3.73-fold difference on the same series (Fig. 1c). Counted on accessions across the full weighted frame, 7.4% qualify; the donor-counted figure of 4.1% is conditional on the resolvable stratum and is reported as such, with its denominator stated, rather than extrapolated.

This is the audit’s own layer measuring the audit’s own thesis, and we report it in that form deliberately. A design that reads sample counts off an accession listing will overstate its power, and the overstatement is large enough to move a study across a replication threshold.

### 2.6 Accession-to-donor inflation across the census frame

Across the census frame, sample accessions exceeded distinct donors by 2.65-fold in the 247 series where donor identity could be resolved. Because that stratum is not randomly selected, we report the conditional estimate together with worst-case and best-case population bounds under stated assignment assumptions [12], an Imbens–Manski 95% confidence interval, and an inverse-probability-weighted sensitivity estimate (Table 4, Fig. 4). All four are computed on the same frame; an earlier version of this analysis paired a point estimate from one population with bounds from another, and the bounds did not bracket the estimate they accompanied.

Resolvability is not random with respect to inflation. Donor identity is most often resolvable in exactly those designs — paired sampling, treatment arms, multiple regions, longitudinal series — that inflate the ratio most, while a study with one sample per donor and no donor field appears unresolvable yet has no inflation at all. The direction supports this: median inflation is 3.67 among series carrying a structural repeat-design marker against 2.73 among those without. The test, however, does not reach significance (Wilcoxon rank-sum, W = 8,321.5, p = 0.086, 101 versus 146 series), and we report it as a direction with its p-value rather than as a demonstrated mechanism.

We had hypothesised that poorly documented studies would also be poorly deposited, so that metadata and data unavailability would compound. On the census frame the association runs the other way: reads are public for 91.1% of donor-resolvable series against 97.1% of non-resolvable series (Fisher exact p = 3.2 × 10⁻⁵, odds ratio 0.309). Series that document their donors are somewhat *less* likely to have public reads, not more. We report this as found and do not have a mechanism for it.

**Table 4.** Accession-to-donor inflation, all statistics computed on the census frame.

| Quantity | Value | Basis |
| --- | --- | --- |
| Conditional estimate | 2.65× | Resolvable stratum, $n = 247$ series |
| Population lower bound | 1.09× | Every unresolved series assigned inflation 1 |
| Population upper bound | 10.77× | Every unresolved series assigned the maximum observed (20.0×) |
| Imbens–Manski 95% CI | 1.07–11.88× | Bootstrap B = 2,000; covers the parameter, not the identified set |
| IPW sensitivity estimate | 2.67× | Missing-at-random given log sample count, characteristics-key count, repeat-design marker, title-stem repetition |
| Enrichment test | W = 8,321.5, p = 0.086 | Wilcoxon rank-sum; median 3.67 with repeat-design marker versus 2.73 without |
| Resolvability × read availability | p = $3.2 \times 10^{-5}$, OR = 0.309 | Fisher exact; 91.1% versus 97.1% reads public |

The upper bound is a function of the largest observed per-series value, which is capped by a declared post-hoc guard against grouping variables being read as donor identifiers. Bounds are assumption-based, not sharp identification.

**Fig. 3.**
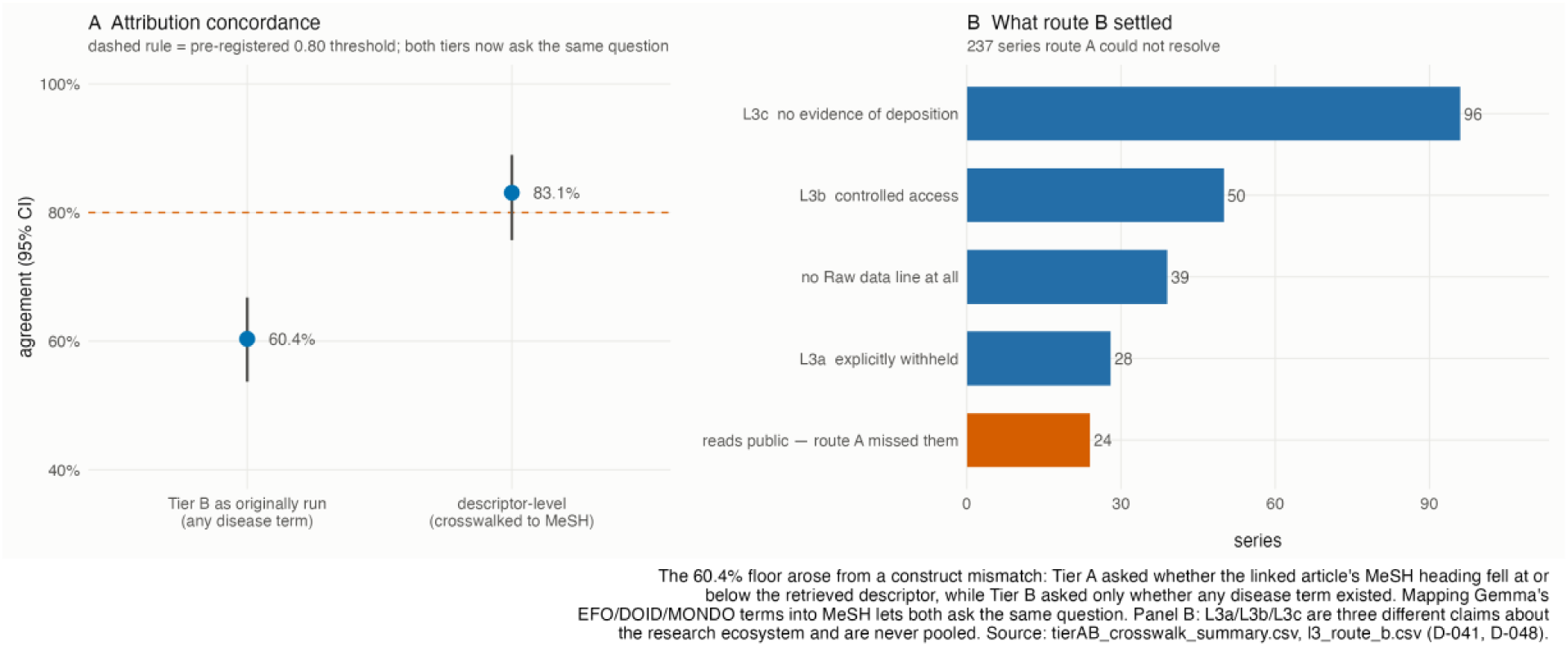
Attribution validation and the resolution of read availability. (a) Concordance between tree-based attribution and an independent curated source, before and after crosswalking the curated resource’s terms into MeSH so that both tests ask the same question. The dashed rule is the pre-registered 0.80 threshold. (b) Outcomes for the 237 series the programmatic route could not resolve. L3a, L3b and L3c are three different claims about the research ecosystem and are never pooled; 24 series are contradicted by the primary record and have public reads.

**Fig. 4.**
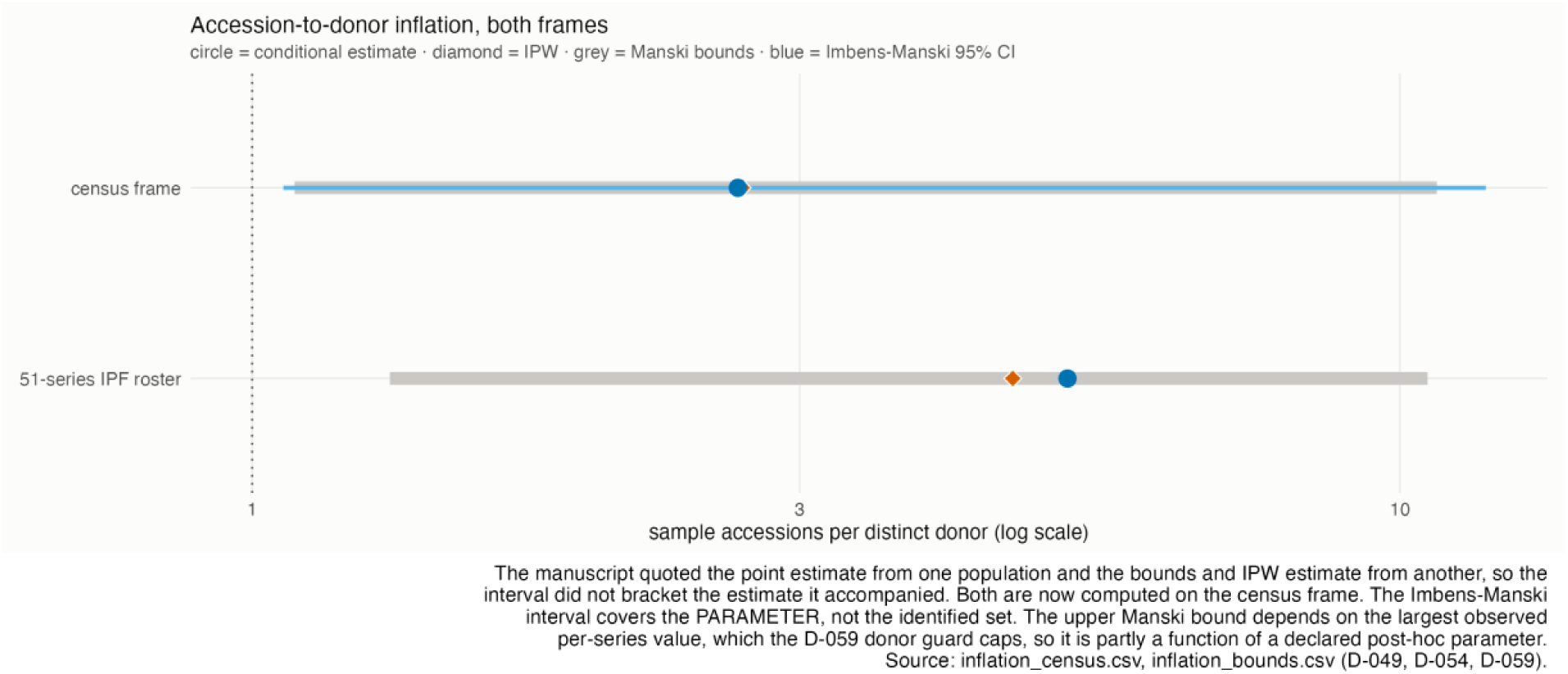
Accession-to-donor inflation, computed on two frames. Circles are conditional estimates, diamonds inverse-probability-weighted estimates, grey bars Manski bounds and blue bars Imbens–Manski 95% intervals. All statistics for a given frame are now computed on that frame; the interval brackets the estimate it accompanies. The 51-series roster is retained as a smaller, more intensively curated comparison and not as the source of population bounds.

### 2.7 A third of keyword attributions are not supported by the source publication

Retrieval assigns each series to the disease term under which it was found. We verified those assignments independently by following each series to its linked publication and adjudicating that publication’s curated MeSH headings against the MeSH tree: an assignment was confirmed when a heading fell at or below the retrieved descriptor, and refuted when none did. Because adjudication is structural rather than lexical, sibling descriptors cannot cross-attribute — a study of one disorder cannot be credited to its sibling because the two share a word.

Of 4,536 series-descriptor pairs, 45.9% were confirmed, 36.4% refuted, and 17.7% unresolved for want of a linked, indexed publication (Table 5). A refutation rate above one third means a conventional keyword query assigns a substantial minority of studies to diseases their own publications do not support.

Stratifying by indexing era at a 1 July 2022 boundary — set shortly after the National Library of Medicine completed its transition to automated MeSH assignment with human quality review on selected subsets in April 2022 [26] — refutation was 41.6% for human-indexed and 45.4% for machine-assisted records. We report the 3.8-point difference and do not over-interpret it: era is confounded with publication recency, journal mix and topic. Independent evaluation of automated indexing has reported systematic differences from human indexing in check-tag assignment [27], consistent with a real but small effect.

**Table 5.** Attribution verification outcomes and era stratification.

| Outcome | Pairs | Share | Definition |
| --- | --- | --- | --- |
| Confirmed | 2,084 | 45.9% | A curated MeSH heading of the linked publication falls at or below the retrieved descriptor |
| **Refuted** | **1,650** | **36.4%** | No heading falls at or below the retrieved descriptor |
| Unresolved | 802 | 17.7% | No linked, MeSH-indexed publication; never recorded as refuted |
| Pre-2022, human-indexed | 1,169 | 41.6% refuted | Of pairs with a resolved outcome |
| Post-2022, machine-assisted | 2,565 | 45.4% refuted | Of pairs with a resolved outcome |

*Era is confounded with publication recency, journal mix and topic. The difference is reported and not interpreted causally.*

### 2.8 Attribution accuracy, validated against an independent curated source

Comparing our tree-based attributions against an independent curated resource requires the two to be asking the same question. Our test asks whether a specific descriptor or its descendants apply; a naive comparison against a curated annotation asks only whether any disease annotation exists. Under that construct mismatch the apparent concordance was 60.4% — a floor rather than a validation, and one that left the pre-registered 0.80 decision threshold undetermined.

Mapping the curated resource’s EFO, DOID and Mondo terms into MeSH through published ontology mapping files [10,11,25] lets both tests ask the same question. At descriptor level, concordance is **83.1% (113 of 136 comparable series, 95% CI 75.7–89.0%)**, above the pre-registered threshold (Fig. 3a). Restricting to exact mappings and admitting broad mappings give the identical result; adding database cross-references gives 82.9% (150 of 181). Terms that could not be mapped are recorded as unresolved and never as disagreement. The pre-registered threshold is therefore met, and the limitation recorded during the original run is closed rather than merely bounded.

### 2.9 Retained diseases and their breadth

Applying the pre-registered retention rule of at least five qualifying series per disease gave 5 diseases and 34 qualifying series among directly verified records, and a weighted estimate of 15 diseases and 147 qualifying series across the full frame. Weighting is required rather than optional: the retention threshold was written for a census and systematically undercounts when applied to a partially sampled frame, in which the sampled stratum was drawn at 21.5%. Retained diseases spanned ten MeSH categories, indicating the cascade is not an artefact of one disease area (Table 6).

**Table 6.** Retained diseases under the pre-registered rule of at least five qualifying series per descriptor.

| Quantity | Value | Note |
| --- | --- | --- |
| Diseases retained, directly verified | 5 | Breast neoplasms, arthritis, pancreatic neoplasms, lung neoplasms, dermatitis |
| Qualifying series, directly verified | 34 |  |
| Diseases retained, weighted estimate | 15 | Retention threshold written for a census; undercounts on a sampled frame |
| Qualifying series, weighted estimate | 147 | Pre-registered $\geq 300$ target retired; see Discussion |
| **MeSH categories spanned** | **10** | C01, C04, C05, C06, C08, C10, C14, C15, C17, C19 |
| Highest-yield diseases (weighted) |  | Breast neoplasms 32.8; diabetes mellitus 13.9; pancreatic neoplasms 12.3; lung neoplasms 11.6; thyroid neoplasms 10.3 |

Generated from the retention table rather than transcribed. The full per-descriptor cascade, including descriptors yielding no qualifying series, is Additional file 1.

### 2.10 The resources that would verify attribution are themselves sparse

Independent verification of disease attribution requires a curated annotation source. At census scale across all 3,631 candidates, Gemma held records for 263 series (7.2%) and MetaSRA for 341 (9.4%, bounds 9.4–17.0%). The curated resources a reader would use to check the assignments in this or any comparable audit are therefore about as sparse as the reads and the donor metadata. We report this inside the cascade rather than as a methodological footnote, because it is a further layer of the same finding.

### 2.11 Automated screening filters systematically over-count eligible studies

Two classes of bias inflate apparent study availability, and both were characterised by running the pipeline against primary records. We report them because both will recur in any comparable automated screen (Table 7).

First, naive text filters over-admit. A filter specified for bulk primary tissue admits a study of cultured primary fibroblasts under drug and cytokine treatment, because “primary cultured fibroblasts” satisfies a naive match on “primary tissue”. Four distinct failure modes were characterised and each is pinned by a regression test: exact-term matching defeated by a misspelling present in the repository’s own metadata; study-level summary text describing validation experiments that were never sequenced, which caused genuine bulk-tissue series to be wrongly excluded; short-acronym collisions, where a match intended for a culture method fired on an unrelated clinical abbreviation; and metadata key-name collisions, where a term intended to match an experimental variable fired on a quality-control field.

Second, retrieval itself is imprecise in a way that is documented but easily missed. Quoted multi-word phrases submitted to the GEO DataSets database do not enforce phrase matching; when a phrase is absent from that database’s phrase index the terms are silently combined, so multi-word disease names lose their qualifiers and sibling diseases become indistinguishable. The effect is direct and measurable: under the unfielded template, queries for two distinct sibling descriptors returned identical hit counts, because the shared leading token determined the result. Before correction, retrieval produced 1.63 disease attributions per series, one series was attributed to 43 diseases, and eight to more than ten. Field-targeted phrase queries against title and description fields, with comma-inverted MeSH labels rotated into natural-language order, substantially reduce this; without the rotation the corrected template under-retrieves catastrophically, returning zero hits for inverted labels such as “Arthritis, Rheumatoid”. We report the measured imprecision rather than only the correction, since the behaviour is a property of the retrieval system that other automated screens will inherit [28,29].

**Table 7.** Characterised failure modes of automated screening, with the correction applied in each case.

| Failure mode | Mechanism | Correction |
| --- | --- | --- |
| Misspelling in repository metadata | Exact-term matching defeated by a misspelling present in the source record | Stemmed matching; regression test per mode |
| Study-level text describing unsequenced work | Summaries describe validation experiments never sequenced, wrongly excluding genuine series | Sample-level fields decisive; study-level text flags for review only |
| Short-acronym collision | A term intended for a culture method matched an unrelated clinical abbreviation | Spelled-out forms only for ambiguous acronyms |
| Metadata key-name collision | A term intended for an experimental variable matched a quality-control field name | Key and value scope discipline |
| Silent phrase tokenisation in retrieval | Quoted multi-word phrases are combined term-wise when absent from the database phrase index, so sibling diseases become indistinguishable | Field-targeted phrase queries with label de-inversion; measured imprecision reported |
| Sample-accession counting | Sample accessions counted as biological replicates, inflating apparent sample size | Distinct-donor resolution with three-valued outcomes, reported as a separate layer |
| Grouping variable read as donor identifier | A substring match on donor-like field names admitted grouping columns, collapsing thousands of samples onto a handful of apparent donors | Field-name rule plus a declared plausibility cap |

*Each correction is pinned by a regression test in the archived repository.*

### 2.12 What the screening threshold conditions away

Every rate above is conditional on a bulk, Illumina, at-least-25-sample population. We measured that conditioning by drawing a stratified random sample of 1,262 of the 13,189 excluded series and running the coverage layers on them (Table 8, Fig. 5).

The result runs against the audit’s interest and is reported at equal prominence. Usable coverage is *higher* in the excluded population — 35.7% against 27.3% in the audited population (p = 1.8 × 10⁻⁸) — driven by the sub-25-sample stratum at 43.0%. The audited population is therefore the pessimistic case for usable coverage among sequencing-based series, and the headline rate does not generalise upward to what the screen removed. Two features qualify this. The finding is confounded with size by construction: usable coverage requires at least 90% of a series’ runs, and one missing run costs 12.5% of an eight-run series against 2% of a forty-nine-run series, so small series are mechanically advantaged. And the array stratum returns exactly 0%, which is the expected result — array studies have no reads to reprocess — and serves as an internal check that the measurement behaves.

**Table 8.** Coverage of the population removed by screening, measured on a stratified random sample.

| Population | n sampled | L3 | L4 | L5 | Median samples |
| --- | --- | --- | --- | --- | --- |
| AUDITED ($\geq 25$ samples, bulk, Illumina) | 3,631 | 94.1% | 46.7% | 27.3% | 49 |
| EXCLUDED (all reasons) | 1,262 | 96.0% | 48.5% | 35.7% | 9 |
| fewer than 25 samples | 907 | 97.8% | 54.7% | 43.0% | 8 |
| non-Illumina platform | 50 | 98.0% | 50.0% | 36.0% | 11 |
| single-cell / single-nucleus | 255 | 90.6% | 34.5% | 16.9% | 10 |
| array platform | 50 | 90.0% | 6.0% | 0.0% | 42 |

Excluded population sampled from 13,189 series. L5 difference against the audited population: $p = 1.8 \times 10^{-8}$ overall, $5.4 \times 10^{-20}$ for the sub-25-sample stratum. The array stratum at 0% is an internal validity check.

**Fig. 5.**
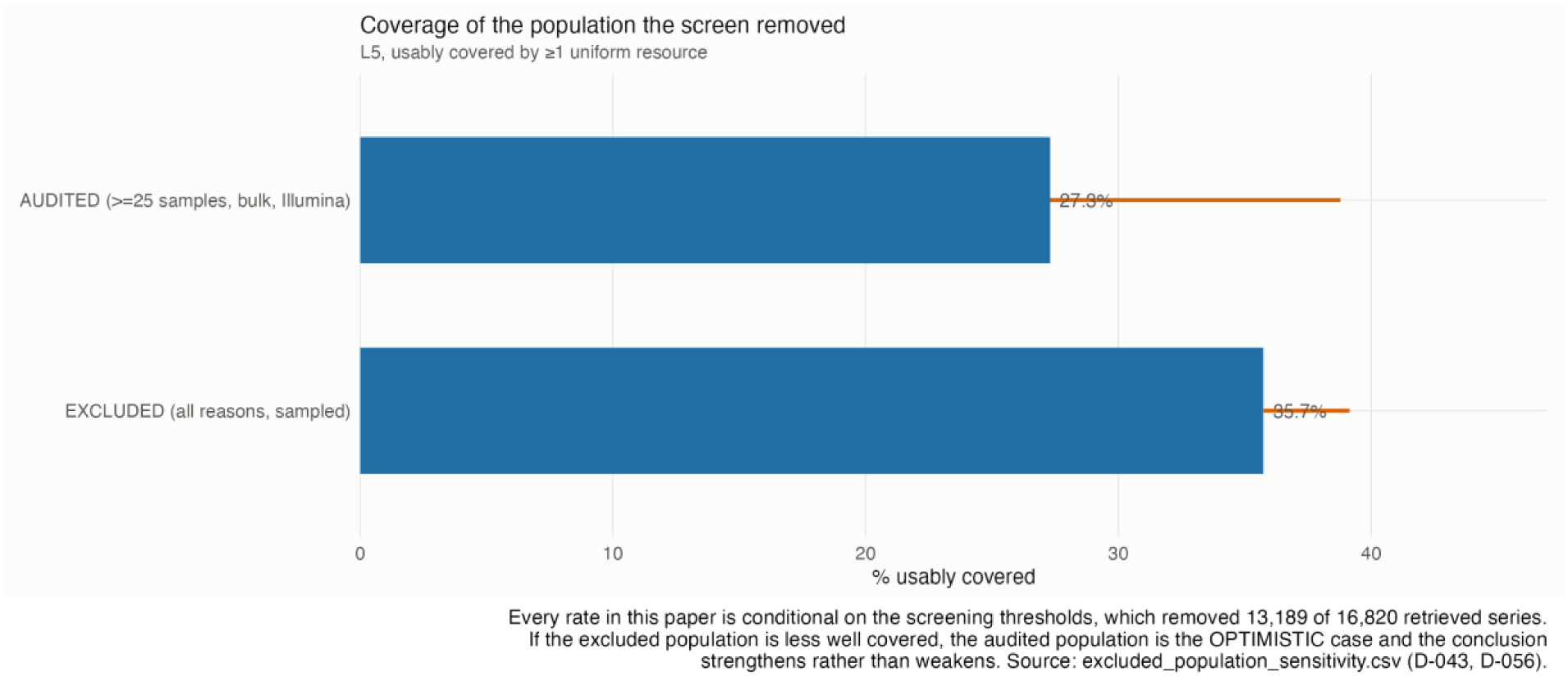
Usable coverage in the audited population against the population the screen removed. Bars are confirmed counts; rules extend to the upper bound implied by unassessable lookups. Coverage is higher in the excluded population, which is the opposite of the direction that would have favoured the audit’s headline. The comparison is confounded with series size by construction, because usable coverage is a proportion of a series’ own runs and small series are mechanically advantaged; this is stated with the result rather than used to set it aside.

## 3. Discussion

The practical implication is narrow and firm: for the population of published human disease RNA-seq we audited, uniform reanalysis is not broadly available, and the constraint is coverage rather than deposition. Reads are archived for the large majority of studies, yet fewer than a third are usably reprocessed by any compendium. A study design that assumes uniformly reprocessed counts will be obtainable for its cohorts of interest is likely to be wrong, and the failure will surface only after cohort selection.

The loss is distributed across independent layers, which matters because the layers have different remedies. Design ineligibility is a property of the literature and is not addressable by infrastructure. Withheld reads are frequently a considered decision about patient privacy in human tissue studies, and we do not read the deposition gap as negligence: a submitter who withholds reads from identifiable patient material and deposits counts instead has made a defensible choice, and the resulting unreprocessability is a cost of that choice rather than a compliance failure. That is why the three absence categories are reported separately and never pooled. Coverage gaps, by contrast, are directly addressable — they reflect the age of a compendium’s snapshot, the breadth of its ingestion, and the strictness of its quality filtering, all decisions maintainers can revisit.

The attribution result has a separate implication for how reuse studies are assembled. If a keyword query assigns more than a third of studies to diseases their own publications do not support, then any automated study set built by keyword retrieval and not independently verified will contain a substantial fraction of off-topic studies. This affects meta-analyses, benchmark corpora and training sets alike, and the correction is available: curated MeSH headings from the linked publication, adjudicated on the ontology tree rather than by string overlap. Validated against an independent curated source through an ontology crosswalk, that adjudication agrees at 83.1%. The magnitude of the underlying retrieval problem is consistent with earlier work on the recall and bias of identifier-based dataset retrieval [18] and with the recall-precision trade-offs documented for automated study identification in systematic reviews [16,17].

The replication-inflation result extends to bulk RNA-seq a problem characterised for single-cell data, where treating cells as independent replicates inflates false-positive rates [32]. Our estimates are conditional by necessity, but the direction is unambiguous and the magnitude is large enough to matter for any power calculation that reads sample counts off an accession listing. The clearest form of the result is the L1 comparison itself: on the same 199 series, the replication criterion is met by 15.1% counted on accessions and 4.1% counted on donors.

Three features of the design are worth separating from the findings. First, every yield threshold set before measurement — a minimum retained-disease count, a minimum qualifying-series count — proved optimistic, because each estimated a yield for a population this study was built to characterise. The minimum qualifying-series target was retired rather than missed: it was never a validity threshold, and at the achieved n a proportion near 0.25 carries a 95% confidence interval of roughly ±7 points against ±5 at the original target. Population claims about GEO from comparable or smaller frames are established practice [5]. Second, the analysis plans were frozen, their integrity is verifiable from the archived repository, and their existence is anchored to an external timestamp rather than asserted from commit metadata. Third, several results in this paper ran against the direction that would have suited it — the excluded population is better covered, not worse; MetaSRA exceeds Gemma rather than matching it; donor-resolvable series are less likely to have public reads; and the enrichment argument is directionally supported but not significant. We report these at the same prominence as the convenient ones, because selective reporting is among the practices this paper exists to measure.

### 3.1 Limitations

**Donor-level replication is conditional.** L1c is computed on the 199 series in which donor identity resolves, roughly 5.5% of the audited population. It is reported as a conditional statistic with its denominator stated, never extrapolated. That a donor-level replication criterion is not measurable at population scale from GEO metadata is itself an obstacle to reuse and is reported as a finding.

**Case membership is assigned by exclusion.** Because the frozen enumeration selects descriptors at intermediate MeSH depth, sample-level text frequently names a specific entity that does not match the broad descriptor label; requiring a literal match reduced qualifying pairs by roughly 97% and retained no disease. Sample-level classification therefore treats a sample as a case unless it matches a control lexicon, a third-group lexicon, or lacks usable text. This is a liberal definition and inflates case counts; the retention result depends on it. The sensitivity analysis under literal matching is Additional file 5.

**Coverage figures are bounded, not exact.** 421 series carry at least one lookup that could not be performed. Coverage is also snapshot-dependent and will change as resources ingest new studies.

**Population conditioning does not resolve in our favour.** All rates are conditional on bulk, Illumina series of at least 25 samples, and the excluded population is better covered (Table 8). The audited population is the pessimistic case for L5 among sequencing-based series, and the headline rate should not be generalised to smaller studies.

**Grain.** The enumeration sits at intermediate MeSH depth — broad disease categories — whereas research practice targets specific entities. A depth-stratified sub-analysis was the designed mitigation; at the achieved yield only one descendant descriptor independently cleared the retention threshold, so the sub-analysis is too thin to serve.

**Stratum confounding.** One sampling stratum was defined using a criterion identical to a coverage layer, so that layer is complete within it by construction. Coverage statistics are quoted from the full census or the second stratum alone, never from the first or an unweighted pool.

**Discoverability, not existence.** Read availability reflects what is discoverable through the routes queried. The dual-route standard raised the figure from 93.5% to 94.1%, and 39 GEO records carry no raw-data status line at all, so the layer’s evidence base is itself incomplete.

**Search peer review was not performed.** PRISMA-S item 14 asks whether the search strategy was peer reviewed. It was not, and the item is reported as not done rather than left blank.

## 4. Conclusions

Uniform reanalysis of published human disease RNA-seq is unavailable for most studies in the population audited, and the binding constraint is usable coverage by reprocessing compendia rather than deposition of raw reads. Automated retrieval further over-counts eligible studies, both by admitting designs outside scope and by assigning studies to diseases their publications do not support, and sample accessions overstate biological replication severely enough to move studies across a replication threshold. Reuse-based study designs should verify the availability of uniform counts for their specific cohorts before committing to a design that requires them, verify disease attribution against curated headings rather than trusting a keyword query, and resolve replication to donors rather than accessions. Three concrete steps follow: compendium maintainers should publish machine-readable per-series usable-coverage indices rather than presence lists; submitters should deposit a donor identifier field even when reads are withheld; and journals should require that reuse-based analyses state the compendium and coverage threshold on which their cohort selection depends.

## 5. Methods

### 5.1 Pre-registration and verification

Three analysis plans were frozen before the work each governs, each in its own file so that amending one never requires lifting another’s protection: the retrieval and disease-selection plan, committed before any coverage query was issued; the attribution-verification plan; and the verification sampling plan, committed before the sample was drawn. A fourth configuration file records the screening thresholds and is **explicitly declared post hoc**; it carries no freeze, and the ≥25-sample threshold it records removed 9,967 series. Every rate in this paper is conditional on it.

Freeze integrity is asserted from the repository history by an automated test rather than claimed: no pre-registered parameter changed after its freeze, verified by comparing parsed configuration values across every subsequent commit. One commit after the retrieval freeze modified that file to record its own freeze hash in a header comment, changing no parameter; the test reports the parsed-value identity and the byte difference separately. The four freeze commits carry OpenTimestamps proofs anchored in the Bitcoin blockchain at blocks 961233 and 961237, mined on 6 August 2026 at 01:48:11 and 02:29:29 UTC. Each proof therefore establishes that its freeze existed no later than that time, independently of the repository’s own commit metadata, which is self-asserted and rewritable; the anchor is an upper bound on the time of freezing rather than a record of it. Verification requires either a Bitcoin node or the public OpenTimestamps verifier, and the command-line client reports an error where no node is available; that error reflects the absence of a node and not a failed proof. All 40 deviations from the pre-registered plans in the original run, including five defects found in the analysis code, are disclosed in the archived repository, together with a further 19 recorded during a subsequent remediation pass; counts are generated by test rather than transcribed.

### 5.2 Disease enumeration and retrieval

Disease terms were enumerated from MeSH descriptors at tree depths 2 and 3 within the declared human-disease categories, retrieved from the National Library of Medicine’s MeSH RDF SPARQL endpoint with paginated queries. English labels only were retained. The retention rule — a disease enters the audit if at least five series survive design eligibility — was fixed before any coverage was measured, and no disease was added or removed afterwards; diseases yielding no qualifying series are reported alongside those that did. Series were retrieved from the GEO DataSets database via Entrez. The revised retrieval template targets the title and description fields with quoted phrases and includes each descriptor’s MeSH entry terms and comma-de-inverted label forms as alternates; the original template used an unfielded quoted phrase, whose silent tokenisation is characterised in Results. Both templates were run and the difference is reported as a result. Full query strings, field tags, limits and retrieval dates are reported against PRISMA-S [31] in Additional file 6.

### 5.3 Screening and the audited population

Retrieved series were screened on four criteria before any coverage was measured: single-cell or single-nucleus protocol, microarray platform, non-Illumina sequencing platform, and fewer than 25 samples. Attrition at each step is in Table 1. The screening rule is implemented in a committed script that asserts it reproduces the audited population of 3,631 exactly.

### 5.4 Design eligibility and replication counting

Design eligibility (L1a) required a human bulk RNA-seq case–control design in primary tissue. Cultured-cell, isolated-cell and perturbation designs were excluded on stemmed matches against sample-level fields

— source name and characteristics keys and values — with study-level title and summary text used only to flag records for review, because summaries describe experiments that were often not sequenced. Stemmed rather than exact matching was necessary: the repository’s own metadata contains misspellings that defeat literal terms. Short acronyms were excluded from automatic matching after one collided with an unrelated clinical abbreviation, and key-name scope was enforced after a term intended for an experimental variable matched a quality-control field.

The replication criterion required at least 15 case and 10 control units for the descriptor under which the series was retrieved, evaluated per series-descriptor pair because case membership is defined relative to the descriptor. Sample-level classification assigned a sample as control on a control lexicon, as a third group on a competing-disease lexicon, as ambiguous where sample text was absent or under eight characters, and as a case otherwise; the consequences of this by-exclusion definition are addressed in §3.1 and Additional file 5. The criterion was evaluated twice: on sample accessions (L1b) across the full frame, and on distinct donors (L1c) on the stratum where donor identity resolves. Distinct-donor resolution is three-valued — resolved, ambiguous or unresolved — and a constant identifier column is not treated as an identifier. Donor-like field names are matched by an explicit name rule rather than a substring match, and a declared plausibility cap on samples per unit guards against grouping variables such as treatment group being read as subject identifiers; without that guard a single series contributed an apparent 4,574-fold inflation from two values of a grouping field across 9,148 samples. Every pair records which basis its counts were taken on.

### 5.5 Read availability

Read availability was determined by linking each series to its project record and querying the sequence archives, with ENA queried independently of SRA, since presence in one and absence from the other is a statement about archive synchronisation rather than about the study. A series is recorded as reads-absent only where an independent programmatic route and a direct reading of the primary GEO record agree; disagreements were adjudicated against the verbatim record text. Both the single-route and adjudicated figures are reported. Controlled-access repositories were checked before any series was considered undeposited. Explicitly withheld, controlled-access and not-deposited outcomes are recorded as three separate categories and never pooled.

### 5.6 Compendium coverage

Coverage was determined for recount3, ARCHS4 and DEE2 by querying each resource’s own index. Coverage is the fraction of the series’ own sequencing runs present in the resource, computed as a run-level intersection and therefore bounded in [0,1]; a series is usably covered when that fraction is at least 0.90 and, where a resource publishes per-run quality flags, only quality-passing runs are counted. Where a resource could not be queried for a series — chiefly where the series-to-project mapping was unresolved the outcome is recorded as unassessable and is neither counted as covered nor as absent, which is why L4 and L5 are reported as intervals with the unassessable count stated. Presence and usable coverage are reported separately throughout. ARCHS4 gene-level values are pseudoalignment estimated counts and are not identical to alignment-based counts from other resources; author-deposited matrices distributed alongside compendia were not counted as uniform reprocessing. Resources were accessed on 4–5 August 2026; versions are given in Table 3.

### 5.7 Attribution verification and ontology crosswalk

Each series was linked to its publication and that publication’s curated MeSH headings retrieved. An attribution was confirmed when a heading fell at or below the retrieved descriptor in the MeSH tree, determined by tree-number descent, and refuted when none did; ancestors were obtained by tree-number truncation, and a series contributing multiple descriptors under one ancestor was counted once for that ancestor. Sibling descriptors cannot cross-attribute under this scheme because their tree numbers diverge above their shared ancestor. Records lacking a linked, indexed publication were recorded as unresolved and never as refuted. Linked-publication coverage was measured before the design was fixed: 79.2% of candidates carried a linked PubMed identifier against 79.7% population-wide, so publication linkage introduces no detectable selection. Outcomes were stratified at a 1 July 2022 boundary, set after the April 2022 completion of the National Library of Medicine’s transition to automated indexing [26].

For validation against an independent curated resource, the resource’s EFO, DOID and Mondo terms were mapped to MeSH descriptor identifiers using published SSSOM mapping sets and database cross-references [10,11,25], preserving the mapping predicate. Concordance was then evaluated at descriptor level by the same tree-descent rule, so that both instruments ask the same question. Exact-only, exact-plus-broad and exact-plus-broad-plus-cross-reference policies are reported separately; unmappable terms are unresolved, never disagreement.

### 5.8 Sampling, weighting and bounded reporting

Verification was performed as a census in one stratum and a stratified sample in another, drawn at 21.5% with per-category weights; estimates across the full frame are weighted accordingly, and the sampling plan was frozen before verification began. For quantities measurable only in a non-randomly selected stratum notably accession-to-donor inflation — we report the stratum estimate as an explicitly conditional statistic, worst-case and best-case bounds for the population under stated assignment assumptions following a partial-identification approach [12], an Imbens–Manski 95% confidence interval computed by bootstrap with a fixed seed, and an inverse-probability-weighted estimate as a missing-at-random sensitivity analysis with propensity modelled on log sample count, characteristics-key count, a structural repeat-design marker and title-stem repetition. The Imbens–Manski interval covers the parameter, not the identified set. All statistics for a given frame are computed on that frame. A selection-correction model requiring an exclusion restriction was considered and rejected as indefensible for a descriptive statistic.

### 5.9 Evidence standards

Every accession-level claim carries the verbatim status string read from the primary record, stored as a distinct field rather than as a derived indicator. Resolution outcomes are three-valued — resolved, not-found, or unresolved — and an unresolved outcome is never collapsed to absent at any layer; where a layer must be summarised as a single quantity it is reported as a bounded interval with the unassessable count stated. Records that cannot be classified are reported as an explicit category in the denominator rather than dropped, following flow-reporting practice for unassessable records [13,31].

### 5.10 Software and reproducibility

Analysis was performed in R 4.6.1 (2026-06-24) with Bioconductor 3.23 on aarch64-apple-darwin23, using GEOquery 2.80.0, rentrez 1.2.4 and data.table 1.18.4. Every table and figure regenerates from the archived result tables with a single command, which completes in 2.3 minutes across 30 registered stages without contacting any external resource; offline operation is enforced at the transport layer, so an outbound call fails loudly rather than silently fetching. The regression suite comprises 10 test files covering the screening rule, coverage bounds, three-valued handling, replication-counting basis, freeze integrity and deviation counts.

### 5.11 Use of AI tools

The analysis pipeline was written and executed by an AI coding agent (Claude Code, Anthropic; Claude Opus 5) under the direct supervision of the author. The author specified the analysis design and all pre-registered thresholds, reviewed the generated code, and independently verified every numerical result reported here against the archived result tables. No AI system selected the research question, made any design decision, or authored any scientific claim. All reported statistics are produced by deterministic code operating on frozen inputs; no language model was used at inference time to generate any reported value. The author takes full responsibility for the integrity and accuracy of the work, including all AI-generated code and output. A contemporaneous disclosure record, including errors the agent made and corrected, is included in the archived deposit.

## Declarations

### Ethics approval and consent to participate

Not applicable. This study analyses metadata from publicly archived studies; no individual-level or controlled-access data were accessed.

### Consent for publication

Not applicable.

### Availability of data and materials

The complete analysis pipeline, the four configuration files, all result tables, the deviation log, the regression test suite and the reporting checklists are archived on Zenodo at https://doi.org/10.5281/zenodo.21817531. The exact code state used for all reported results is recorded in the deposit, together with OpenTimestamps proofs of the pre-registration freeze commits, anchored in Bitcoin blocks 961233 and 961237. All primary data analysed are publicly available from the Gene Expression Omnibus, the Sequence Read Archive, the European Nucleotide Archive, and the recount3, ARCHS4 and DEE2 compendia; accessions and download scripts are provided in the deposit. Third-party datasets are not re-hosted. No new sequence data were generated.

### Authors’ contributions

ABM conceived and designed the study, specified and froze the analysis plans, supervised and verified all computational work, verified all numerical results against the archived result tables, and wrote the manuscript. The author read and approved the final manuscript.

## Acknowledgements

The author thanks the maintainers of the Gene Expression Omnibus, the Sequence Read Archive, the European Nucleotide Archive, MeSH, recount3, ARCHS4, DEE2, Gemma and MetaSRA, whose public infrastructure made this audit possible.

