## Supplementary files 1-7 for "Most published human disease RNA-seq cannot be uniformly reanalyzed: a population-scale audit of reprocessability in the Gene Expression Omnibus": Additional_file_3_deviation_log.docx

### DEVIATIONS.md

Every departure from docs/SPEC_v1.0.md or docs/SPEC_AMENDMENT_v1.1.md, dated, with the reason. This is a reviewer-facing audit trail and feeds the Methods section. **An honest deviation log strengthens the submission — do not tidy it or omit awkward entries** (CLAUDE.md).

Status values: OPEN (needs a decision), RESOLVED, ACCEPTED (a permanent, declared departure).

**Numbering.** D-026 was never issued; the identifier was skipped between D-025 and D-027. The original run therefore ends at D-041 but contains **40 entries**, not 41 — of which **five**, not four, are marked own defect. MANIFEST.md and docs/CP0b_REPORT.md §8 previously said 41 and four, and the manuscript inherited both errors; all are corrected.

Counts are generated, not remembered: tests/test_deviations.R parses this file and writes results/tables/deviation_counts.csv, which is what the manuscript cites.

The 2026-08-05 remediation reserves **D-042 – D-060** by subject, so each defect keeps a stable identifier from the moment it is described. Entries appear as their fix lands; the reserved identifiers not yet written are listed as remediation_pending in deviation_counts.csv.

#### D-001 — msigdbdf could not be installed; not required by the installed msigdbr

**Date:** 2026-08-04 · **Stage:** 0 · **Source:** amendment §6.1 · **Status:** RESOLVED

Amendment §6.1 requires installing the msigdbdf data package from https://igordot.r-universe.dev, on the basis that msigdbr ≥ 10.0.0 errors without it. The r-universe install failed (package 'msigdbdf' is not available) — see the tail of setup_install.log.

The requirement turned out not to apply to the installed release. msigdbr **26.1.0** (MSigDB **2026.1.Hs**) ships its data and returns gene sets without msigdbdf: Gate 4 verification retrieves 7,331 Hallmark rows and 113,653 C2:CP:REACTOME rows. The collection= / subcollection= API required by amendment §6.1 is the one in use, and collection identifiers are enumerated at runtime with msigdbr_collections() (26 rows) rather than hard-coded.

**Effect on results:** none. **Action:** record MSigDB version 2026.1.Hs in config/config.yaml and the Methods, as amendment §6.1 requires.

#### D-002 — Stage scripts 02–10 are created at their own stage, not at Stage 0

**Date:** 2026-08-04 · **Stage:** 0 · **Source:** spec §7 · **Status:** ACCEPTED

Spec §7 lists src/01_…R through src/10_…R as part of the repository structure. Stage 0 creates the directory tree, src/00_seed.R, src/utils/, and the Stage 1 scripts. Empty placeholder files for stages not yet run would be committed artifacts that do nothing, and would break snakemake -c1 all by advertising rules whose scripts are stubs. Each script is created when its stage runs; the Snakefile is extended in step with them.

**Effect on results:** none — presentational only.

#### D-003 — ENTREZ_KEY not configured; Stage 1 GEO screening blocked

**Date:** 2026-08-04 · **Stage:** 1 · **Source:** kickoff prompt Stage 0(3), amendment §6.2 · **Status:** OPEN

The NCBI API key is not present in the environment (Sys.getenv("ENTREZ_KEY") is empty; no ~/.Renviron). Amendment §6.2 requires one because spec §5.3 makes many getGEO calls and will trip the unauthenticated 3 req/s limit; the kickoff prompt instructs a hard stop before any bulk GEO call if it is unset.

Stage 1 is therefore split. The portion that does not touch NCBI — recount3 / ARCHS4 / DEE2 uniform-pipeline coverage — was executed. The Entrez gds screen (amendment §3.1) and per-candidate GEO sample-table verification (spec §5.1) are held.

**Effect on results:** none yet; Stage 1 is incomplete and CP-0 cannot be presented. **Action required from the human operator:** register a key at https://account.ncbi.nlm.nih.gov/settings/ and export it (see README.md), then re-run Rscript src/01_screen_cohorts.R.

#### D-004 — ARCHS4 membership cannot be determined programmatically

**Date:** 2026-08-04 · **Stage:** 1A · **Source:** amendment §2.1, §2.2 step 2 · **Status:** OPEN

Amendment §2.2 requires querying membership in recount3, ARCHS4 and DEE2 for every candidate. recount3 and DEE2 both expose a public, machine-readable catalogue (recount3::available_projects(); https://dee2.io/metadata/hsapiens_metadata.tsv.cut, 92 MB). **ARCHS4 exposes neither.** Verified 2026-08-04:

- Sample-level metadata lives only inside the human gene HDF5 (meta/samples/series_id). Current build is v2.latest, 1,098,771 samples, **62.3 GB** (/api/versionfile).
- The download endpoint /api/file/download/{id} returns **HTTP 401** — an ARCHS4 account is required.
- The documented API (/api/static/swagger.json) has 23 paths, all file/collection/user management. There is no accession or metadata search endpoint. Legacy Maayan-lab endpoints (maayanlab.cloud/archs4/search/..., maayanlab.cloud/matrixapi/...) return

archs4_has() therefore returns unresolved with this reason, never absent — absence of a queryable index is not absence from the resource. Setting options(ipf.archs4_h5 = "<path>") to a manually downloaded copy enables the check (rhdf5 would need installing; it currently is not).

**Effect on results:** the ARCHS4 column of metadata/uniform_source_coverage.csv is unresolved throughout, so ARCHS4 cannot currently be selected as uniform_source, and the amendment §4 multi-pipeline upgrade to Axis 2 would be a 3-level factor (recount3 / DEE2 / author-deposited) rather than 4-level.

**Decision needed at CP-0:** accept the 3-level Axis 2, or authorise the one-off 62 GB ARCHS4 download (disk is not the constraint — 560 GB free). This is itself a reportable finding about the state of public uniform reprocessing, which amendment §2.2 step 5 anticipates.

*2026-08-04 — operator deferred this decision to CP-0, once it is known whether recount3 or DEE2 covers ≥3 qualifying cohorts. ARCHS4 stays unresolved until then.*

#### D-005 — DEE2 presence is not DEE2 usability

**Date:** 2026-08-04 · **Stage:** 1A · **Source:** amendment §2.2 · **Status:** OPEN

DEE2 assigns each run a QC_summary of PASS, WARN(...) or FAIL(...). Two candidates are present in DEE2 with **zero** QC-PASS runs (GSE99621: 0/23; GSE213001: 0/13), and GSE92592 has 27/39. A presence/absence coverage matrix would score all of these as “covered” and materially overstate DEE2’s usable coverage.

dee2_has() therefore returns the QC-PASS count alongside presence, and metadata/uniform_source_coverage.csv carries it. **The uniform_source selection rule in amendment §2.2 step 3 counts “qualifying cohorts” — at CP-0, decide whether a cohort whose DEE2 runs all fail DEE2’s own QC counts as covered by DEE2.** The plain reading is that it does not.

**Effect on results:** none yet — affects the CP-0 pipeline choice.

#### D-006 — GSE→SRP resolution needed the BioProject route; first Gate 1 result was wrong

**Date:** 2026-08-04 · **Stage:** 1B · **Source:** spec §5.3 Step 2 · **Status:** RESOLVED

The first Stage 1B run resolved GSE→SRP from GEO relation fields only, as in spec §5.3 Step 2’s example code. That works for **12 of 51** screened series: modern GEO series expose a *BioProject* relation (**50 of 51**) and no SRA relation. Because recount3 is keyed by SRP, its coverage column came back unresolved for 8 of 11 qualifying cohorts, and the selection rule — which counts pipelines by cohorts covered — handed uniform_source to DEE2 on that gap. **That run reported Gate 1 PASS with n_tier_a = 5. It was an artifact and has been discarded.**

resolve_srp() in src/utils/entrez.R now tries, in order: SRP from a GEO relation → BioProject relation resolved against SRA via Entrez → for series with no relations at all, a direct SRA search by GSE accession. Spec §5.3 Step 2 permits the elink route explicitly. Outcomes are three-valued (resolved / no_sra_records / unresolved) so that “the reads were never deposited” is recorded as the finding it is rather than as a lookup failure.

Assertion checks pass: GSE92592 → SRP095361 (amendment §1.2 verified value) and GSE213001 → SRP396424 (matches the value independently obtained from DEE2’s GEO_series index at Stage 1A).

**Effect on results:** corrects the Tier A roster and the uniform_source choice. The corrected run is the one reported at CP-0.

#### D-007 — “present in a pipeline” replaced by a sample-coverage threshold

**Date:** 2026-08-04 · **Stage:** 1B · **Source:** amendment §2.2 step 3 · **Status:** OPEN (needs CP-0 approval)

Amendment §2.2 step 3 selects uniform_source as “the pipeline covering the largest number of qualifying cohorts” without defining *covering*. Read as presence of ≥1 run, DEE2 “covers” GSE188464 with **1 of its 48 samples** and GSE169500 with **4 of 30**, and both were seated in Tier A by the first run. Combined with D-005 (GSE213001 present with **0 of 139** runs passing DEE2’s own QC), presence/absence materially overstates usable coverage.

A pipeline now covers a cohort only if it holds usable counts for at least cohorts.pipeline_coverage_min (**0.9**) of that cohort’s GEO samples; for DEE2, “usable” means QC-PASS by DEE2’s own flag. Declared in config/config.yaml as a Stage-1 *screening* rule, confirmed per sample at acquisition.

**The threshold is not load-bearing.** Sensitivity over the qualifying cohorts:

| threshold | recount3 | DEE2 |
| --- | --- | --- |
| ≥50% | 1 | 2 |
| ≥70% | 1 | 1 |
| ≥90% | 1 | 1 |
| =100% | 1 | 0 |

Gate 1 (≥3) fails at every threshold from 50% to 100%, so the conclusion does not depend on where the line is drawn.

**Action:** approve or amend pipeline_coverage_min at CP-0, before the CP-2 freeze.

#### D-008 — Four to five qualifying cohorts have no SRA records at all

**Date:** 2026-08-04 · **Stage:** 1B · **Source:** amendment §2 · **Status:** OPEN — this is the CP-0 decision

Amendment §2 attributes GSE150910’s expected absence from recount3 to the ~2019-10-01 snapshot, and prescribes making the uniform source configurable (recount3 → ARCHS4 → DEE2). Verification shows the diagnosis is incomplete and the prescribed fix does not reach the affected cohorts.

**GSE150910 and GSE134692 have no SRA records whatsoever.** Their BioProjects exist with correct titles (PRJNA634074 — “RNA-Sequencing of Chronic hypersensitivity pneumonitis compared with Idiopathic Pulmonary…”; PRJNA556144 — “…Transplant Stage Idiopathic Pulmonary Fibrosis Lung…”), and all three of elink(bioproject → sra), esearch sra "PRJNA634074[BioProject]" and a free-text SRA search return **zero**. GSE173355, GSE205525 and GSE199949 are in the same position. These studies deposited count matrices to GEO and no reads to SRA.

recount3, ARCHS4 and DEE2 all reprocess SRA. A study absent from SRA cannot appear in any of them, now or later. **These cohorts are Tier B by construction, for any choice of uniform_source** — including the two largest IPF bulk RNA-seq cohorts in existence (103 IPF / 103 control and 46 / 26).

This also confirms amendment §1.2’s warning: PRJNA635732 is not GSE150910’s BioProject. PRJNA634074 is, resolved programmatically at runtime.

Where reads *are* in SRA, SRA holds the complete set — 27/27, 48/48, 60/60, 139/139, 30/30 and 39/39 runs for the six SRP-resolved qualifying cohorts. **The shortfall is in the reprocessing pipelines, not in the archive**, which is a sharper and more useful finding for a benchmarking paper than “recount3 stops in 2019”. It is exactly the reportable result amendment §2.2 step 5 anticipates.

**Consequence for D-004:** the ARCHS4 question is no longer optional.

#### D-009 — Resolver defect: SRA-search-by-GSE produced false read-absent calls

**Date:** 2026-08-05 · **Stage:** 1C · **Source:** CP-0 resolution Task 1 · **Status:** RESOLVED — **my code was at fault**

Amendment v2.0 §5.1 flagged GSE199949 as disputed: Stage 1 recorded no SRA records, the GEO record says reads are in SRA. **The GEO record is correct and my code was wrong.** PRJNA819837 holds **42 SRA runs**.

**The defect.** resolve_srp() tried, as a last resort for series with no relation field, esearch(db="sra", term=<GSE accession>). **SRA does not index GEO series accessions**, so that query returns 0 for every series ever passed to it. GSE199949’s cached SOFT file carries no relation field at all, so it fell through to that route and was recorded no_sra_records — a false negative asserted with full confidence. This is the worst failure mode available to this project: a wrong answer that looks like a finding.

**The fix.** The GSE-accession route is removed and must never return. The chain is now relation → elink(gds → bioproject) → esearch(sra, "PRJ…[BioProject]"). The middle step is new and is what recovers GSE199949: the SOFT file has no relation, but elink(gds → bioproject) returns 819837. elink(gds → sra) is also deliberately **not** used — it returns 0 links for GSE199949 despite the BioProject holding 42 runs, so it is a second source of false negatives.

**Scope of the error.** Every L3 negative has been re-derived from scratch by src/01c_rederive_l3.R — not only the ones previously recorded absent — with dual-route agreement required (CLAUDE.md rule 4), independent ENA lookup, and a dbGaP/EGA check before any L3c. Nothing produced by the defective route is carried forward.

**Two further bugs found while building the direct-GEO route**, both of which would have inverted availability calls:

1. Detagging the GEO HTML removes the element boundary, so “Raw data not provided for this record” runs directly into “Processed data are available on Series record”. The capture now truncates at the next field, otherwise an availability test reads the *processed*-data phrase.
2. "not available" contains "available". The availability test now checks negation **first**; the opposite order marks every absent record present.

**Effect on results:** GSE199949 moves from reads-absent to reads-present. All other L3 calls re-derived; see results/tables/l3_rederivation.csv.

#### D-010 — Shipped config_v2.yaml was not valid YAML

**Date:** 2026-08-05 · **Stage:** CP-0 · **Source:** operator bundle · **Status:** RESOLVED

benchmark.shared_source_risk had handling: at the same indentation as the - sequence items below it. YAML does not permit a mapping key and sequence entries at one level, so the **entire config failed to parse** and every script sourcing src/00_seed.R aborted.

Fixed by nesting the three cohort entries under shared_source_risk.cohorts: so handling: can sit beside them. **No content changed** — same three cohorts, same handling text. Access path is now benchmark$shared_source_risk$cohorts.

Separately, pipeline_coverage_min moved from cohorts: (v1) to audit: (v2); src/01_screen_cohorts.R updated to read the new path and to fail loudly rather than silently defaulting if it is absent from both.

**Effect on results:** none. Mechanical repair.

#### D-011 — esearch sra "PRJ…[BioProject]" is unreliable; no single Entrez route may report absence

**Date:** 2026-08-05 · **Stage:** 1C · **Source:** CP-0 resolution Task 3 · **Status:** RESOLVED

The dual-route check from D-009 immediately caught a **third** false-negative route — the one amendment v2.0 §2.4 and the CP-0 prompt both name explicitly. esearch(db="sra", term="PRJ…[BioProject]") returns the correct count for some BioProjects and **silently returns 0 for others whose records plainly exist**:

| BioProject | [BioProject] field | free-text | elink(bioproject→sra) |
| --- | --- | --- | --- |
| PRJNA326204 | **0** | 223 | 223 |
| PRJNA549450 | **0** | 32 | 32 |
| PRJNA601416 | **0** | 25 | 25 |
| PRJNA1235680 | **0** | 27 | 27 |
| PRJNA878759 | 139 | 139 | 139 |
| PRJNA358081 | 39 | 39 | 39 |

Ten candidates were wrongly recorded reads-absent by this route. They were caught only because the ENA cross-check disagreed and the dual-route rule refused to classify on a disagreement — the mechanism CLAUDE.md rule 4 mandates, doing exactly its job.

**The general lesson, now demonstrated three times (D-009, this entry twice over): a zero from any single NCBI route is not evidence of absence.** sra_experiment_count() therefore queries three independent routes, takes the maximum, and records whether they agreed. A zero is believed only when field query, free-text, elink, **and** ENA all return zero.

GSE150910 and GSE134692 are unaffected — all four routes return zero for both, so the L3a classification stands on the strongest available evidence.

**Units correction.** db="sra" UIDs are **experiments** (SRX); ENA’s read_run report counts **runs**. GSE213001 is 139 experiments / 695 runs, GSE92592 is 39 / 39. The earlier table compared the two directly, which made normal multi-run experiments look like archive desynchronisation. The column is now n_sra_experiments and the two are reported separately.

**Effect on results:** ten L3 classifications corrected. None were in the qualifying IPF roster, but all ten would have entered Paper 1’s cascade as false L3c.

#### D-012 — Condition counts must be parsed structurally, at patient level, not by regex

**Date:** 2026-08-05 · **Stage:** 1D · **Source:** CP-0 resolution Task 2 · **Status:** OPEN — roster re-verification in progress

Amendment v2.0 §5 disputed the Stage 1 counts for GSE213001 and GSE199949. Both disputes are upheld, and the cause is a defect in the Stage 1 screen affecting **every** cohort.

**Defect 1 — regex over concatenated free text.** CTRL_RX contains normal, and GSE213001’s characteristics include the *key name* diseasenormal: on every sample. Every sample therefore matched “control”. The screen reported 56 cases / 51 controls; the structured diseasegroup field gives 62 IPF / 41 NDC / 26 ILD / 10 CLAD samples.

**Defect 2 — samples counted as if they were patients.** Neither cohort is one-sample-per- patient:

|  | samples | patients | by patient |
| --- | --- | --- | --- |
| GSE213001 | 139 | **46** (1–5 regions each) | IPF 20 · NDC 14 · ILD 9 · CLAD 3 |
| GSE199949 | 42 | **21** (2 sites each) | IPF 13 · control 8 |

The secondary figures amendment §5 called irreconcilable are now explained: **“20 IPF / 9 non-IPF ILD / 14 control” is correct — it counts patients.** “98 IPF / 41 control” counts samples and lumps ILD + CLAD in with IPF. Mine counted samples through a broken regex.

**Consequence for the roster:**

- **GSE213001 qualifies** at patient level (20 IPF, 14 control ≥ 15/10) — but only with one region per patient or donor as a blocking factor.
- **GSE199949 FAILS the ≥15-case gate at patient level** (13 IPF). Amendment §5 suspected “26/16 counts biopsies”; it does, and the patient-level count falls below the threshold. Recommend dropping from the primary roster, or retaining as a declared underpowered held-out set with subject as a blocking factor.

Every remaining cohort is being re-counted the same way: structured key: value parsing, a patient/donor identifier where one exists, and counts reported per patient and per sample.

**Effect on results:** changes the qualifying roster and therefore amended Gate 1 (≥8 cohorts). Reported at CP-0b.

#### D-013 — Amended Gate 1 threshold was mis-specified; corrected from ≥8 to ≥7

**Date:** 2026-08-05 · **Stage:** CP-0b · **Source:** operator decision 3 · **Status:** RESOLVED

**Stated plainly: the ≥8 threshold in amendment v2.0 §3.4 was mis-specified.** It was set to the roster size the amendment expected after its own §5 corrections — “on current evidence this gate passes at roughly 8 cohorts” — not to any analytical requirement. The roster landed at 7 because §5 did not know GSE188464 was cultured fibroblasts under drug perturbation (D-016).

**Amended Gate 1:** ≥7 qualifying cohorts, each with ≥15 IPF cases and ≥10 non-fibrotic controls **counted at the patient level**, author-deposited raw counts, unambiguous labels, independent patient series; plus ≥1 uniform-subset cohort for the Axis 2 bound or a documented reason for dropping Axis 2.

**Rationale (recorded before the roster was re-run):** 7 cohorts give 21 cohort pairs as blocking units for the Axis 3 attribution model and ≥6 studies per LOCO iteration.

The threshold was repaired by amending it with a stated rationale, **not** by admitting a cohort to reach it. Both cohorts short of the gate were resolved on their own merits — GSE188464 rejected on eligibility, GSE199949 held out — and neither decision was taken to change the count.

#### D-014 — k_primary_fraction: 0.75 withdrawn; effect-size meta-analysis promoted to primary

**Date:** 2026-08-05 · **Stage:** CP-0b · **Source:** operator Task A2 · **Status:** RESOLVED

ceiling(0.75 × n) does not hold stringency constant as n moves: n=6 → 83%, n=7 → 86%, n=8 → 75%, n=9 → 78%. A rule that reads as pre-specified therefore silently changes strictness whenever the roster changes — and inside LOCO, where n drops by one every iteration, it changes *within a single analysis*.

Replaced with **k_primary: 5**, fixed a priori for n=7, full sweep k=2..7 reported.

Separately, **vote-counting is demoted from primary to corroboration.** Its power depends on per-study power: when per-study power is low, adding studies can *decrease* the probability of detecting a true effect. The effect-size meta-analysis (already specified, metafor::rma.uni) becomes the primary core-signature definition.

#### D-015 — Shrunken log2FC removed from the concordance input

**Date:** 2026-08-05 · **Stage:** CP-0b · **Source:** operator Task A1 · **Status:** RESOLVED

de.primary_shrinkage: apeglm applied to the concordance input was a hazard specific to this study. apeglm and ashr shrink each gene in inverse proportion to its information, so a small cohort’s log2FCs are compressed toward zero more aggressively than a large cohort’s. Across a panel spanning 27 to 288 samples, that scale mismatch deflates cross-cohort Spearman **by itself** — and does so in proportion to exactly the sample-size differences the study is trying to reason about. The headline number would have been partly an artifact of the shrinkage estimator.

- Concordance consumes **unshrunken MLE log2FC**, uniformly across all cohorts. Spearman is rank-based, so the effect is mostly on near-zero ties. Degen & Medo used unshrunken log2FC for the same reason: “to preserve intrinsic variability”.
- Shrunken (apeglm) values are retained for reporting, forest plots and visualisation, and the concordance analysis is repeated with them as a **declared sensitivity analysis**.
- **Never mix shrunken and unshrunken log2FC across cohorts within one metric** — recorded in config as never_mix_shrunken_and_unshrunken: true.

#### D-016 — GSE188464 excluded on ELIGIBILITY (tissue and design), not sample size

**Date:** 2026-08-05 · **Stage:** CP-0b · **Source:** operator decision 1 · **Status:** RESOLVED

Verified against the live GEO record, not the secondary description. Verbatim overall design:

“Primary **cultered** human fibroblasts from 16 individuals, 8 with idiopathic pulmonary fibrosis (IPF) and 8 age-matched controls”

(GEO’s own misspelling of “cultured”, preserved — and see D-017, it defeats a literal-term filter.) Summary: fibroblasts “treated with 0, 3 or 10 µM ACT001 in the presence/absence of pro-fibrotic cytokine transforming growth factor” β1. Title: “Disease-specific transcriptomic changes induced by NFκB inhibition in idiopathic pulmonary fibrosis.” Illumina NovaSeq 6000 (GPL24676), 48 samples, submitter Nicholas C Wong, Monash University.

So: **cultured cells, not lung tissue, under drug + cytokine perturbation, not an untreated case–control contrast.** 48 samples = 16 donors × dose/treatment arms. Fails spec §5.1 on tissue *and* design.

This also resolves the 24/24 vs 8/8 discrepancy: 24/24 counted sample files grouped by donor disease status, 8/8 counted distinct donors. Both describe the series correctly; the real biological replication is 8 IPF vs 8 control donors.

**Not admissible in any form**, including as a held-out set — a cell-type- and treatment-confounded signature has no place in a tissue-signature benchmark. Recorded under eligibility, not sample size.

#### D-017 — GSE199949 admitted as a declared held-out replication cohort only

**Date:** 2026-08-05 · **Stage:** CP-0b · **Source:** operator decision 2 · **Status:** OPEN — two pre-declarations due before FREEZE-B

Genuine bulk-tissue IPF cohort: Illumina HiSeq 4000 (GPL20301), 42 samples, raw counts deposited. Source publication Huang Y, Guzy R, Ma S-F, Noth I, et al., *BMJ Open Respir Res* 2023;10(1):e001391 (PMID 36725082): “paired central and peripheral lung explant biopsies from 13 IPF and 8 non-IPF donors”. The Stage 1D patient-level count of 13/8 is confirmed.

**Held out rather than primary** because 13/8 patients sits below the replicate count needed for reliable fold-change estimation, which is what the concordance metric consumes (Schurch et al., *RNA* 2016;22(6):839 — ≥6 a floor, ≥12 to identify DE genes at all fold changes, >20 to recover >85%; Degen & Medo 2025 — recall < 0.5 for N < 7). Noisy log2FC mechanically attenuates Spearman, Jaccard, CAT and π₁ through measurement-error attenuation, for reasons unrelated to IPF biology. Admitting it would confound the headline concordance with measurement error and invite the critique that low concordance is an artifact of tiny cohorts.

**Still to pre-declare before FREEZE-B, both marked TBD-FREEZE-B in config:**

1. **Region contrast** — peripheral_only or pooled_regions. Not a formality: this study’s scientific point is that the central, macroscopically non-fibrotic region is *already* transcriptionally abnormal, so “IPF vs control” is ambiguous until the region is fixed, and the choice materially changes the log2FC vector entering the comparison.
2. **Paired handling** — patient-level mean-collapsing (simplest, most comparable to the unpaired cohorts) or variancePartition::dream with block = patient (most powerful; Hoffman & Roussos, *Bioinformatics* 2021;37(2):192).

**Forbidden:** fitting 42 samples with an unpaired model. DEFF = 1 + (m−1)ρ, so effective n is ≈21 patients, not 42. Patient as a *fixed* effect in DESeq2 is also forbidden — DESeq2 has no random-effects support.

A null replication result is **pre-declared power-limited** and will not be read as refutation. Six cohorts have complete public SRA runs; recount3 holds one of them and DEE2 usably holds one. ARCHS4 is continuously updated (build 2026-07-04, 1,098,771 samples) and is the only remaining candidate that could put Gate 1 back in reach. See the CP-0 options.

#### D-018 — BioSample cannot supply donor identity; the absence is a finding

**Date:** 2026-08-05 · **Stage:** CP-0b r2 · **Source:** operator correction 1 · **Status:** RESOLVED

An earlier plan to recover donor identity from NCBI BioSample was wrong and is withdrawn. NCBI’s **Human 1.0** BioSample package has exactly seven mandatory attributes — isolate, age, biomaterial_provider, collection_date, geo_loc_name, sex, tissue — and **none is a subject identifier**. isolate is unconstrained free text. host_subject_id exists but belongs to the MIxS/host-associated and Pathogen packages, not Human, and is optional there. Real subject→sample mappings live in dbGaP behind controlled access, which CLAUDE.md rule 9 forbids accessing.

**Confirmed empirically, not just accepted:** across 42 SRA-backed series, the tiered resolver (D-021) resolved **zero** series from an SRA/BioSample attribute column (tier T2). MetaSRA’s raw_SRA_metadata blob yielded **zero** isolate values across the three series it covers.

**Recorded as a substantive result:** no public resource systematically exposes donor identity for human RNA-seq. That is a finding about the research ecosystem, not a limitation of this pipeline, and it belongs in Paper 1 alongside the metadata-incompleteness literature.

#### D-019 — MetaSRA validation: host had moved; 100% agreement but only 3/42 coverage

**Date:** 2026-08-05 · **Stage:** 1g · **Source:** CP-0b Task B · **Status:** RESOLVED, with a coverage caveat that materially weakens the claim

**Host correction.** The host named in the CP-0b instruction, metasra.biostat.wisc.edu, is dead: DNS resolves to 144.92.73.116 but ports 80 and 443 both refuse connections, verified while NCBI returned 200 from the same machine at the same moment. The service has **moved to metasra.platformx.wisc.edu**, per the MetaSRA-pipeline README. Using the instructed host would have produced a false “resource unavailable” conclusion and silently lost the whole validation.

**Working API form:** /api/v01/samples.csv?study=<SRP>. study_id and sample_id both return empty; limit is ignored on study queries.

**Result: agreement 3/3 = 100%, zero disagreements.**

| Series | SRP | MetaSRA call | conf | filter call |
| --- | --- | --- | --- | --- |
| GSE92592 | SRP095361 | tissue (39/39) | 0.92 | tissue ✔ |
| GSE83505 | SRP076780 | cell line (219/219) | 0.62 | culture ✔ |
| GSE133298 | SRP212077 | cell line (54/54) | 0.96 | culture ✔ |

**This is a spot check, not a validation, and must be reported as such.** MetaSRA covers **3 of 42** SRP-resolved series (7%). 38 are absent, 1 lookup failed (SRP106785 returns a persistent HTTP 500). Three concordant series cannot establish a filter’s accuracy; the honest claim is “no disagreement was found in the three series where independent classification was available.”

**Refresh date: UNVERIFIED.** No version or api-docs endpoint exists — both 404 — so the data refresh cannot be read from the API. Bounded empirically instead, and the bound shows coverage is **patchy rather than a clean date cutoff**: SRP212077 (2019-09-30) is present while SRP108496 (2017) is absent. Latest present is SRP212077.

**Coverage gap is itself reportable** for Paper 1: the only public ontology-based sample-type classifier for SRA covers 7% of a current disease-series set.

MetaSRA carries no donor identifier and no same-individual grouping, so it contributed nothing to Task A, as anticipated.

#### D-020 — The 5.13× inflation figure is conditional; population bounded, selection confirmed

**Date:** 2026-08-05 · **Stage:** 1h · **Source:** CP-0b Decision 2 · **Status:** RESOLVED

| quantity | value | basis |
| --- | --- | --- |
| **Conditional** | **5.13×** | the 15 series where donor identity was resolvable (1,309 GSMs → 255 units); per-series median 4.50, range 2.00–19.33 |
| Population lower bound | 1.32× | every unresolved/ambiguous series assigned inflation 1 |
| Population upper bound | 10.57× | every unresolved/ambiguous series assigned the max observed (19.33×) |
| IPW sensitivity | 4.60× | MAR given size, metadata richness, repeat markers, title repetition |

Bounds are **plausible-range under stated assumptions, not sharp identification bounds**. never_report_as_population_parameter: true is recorded in config.

**A first attempt at the selection model was invalid and is withdrawn.** It derived the repeated-measures marker from unit_field and unit_note — both *outputs* of donor resolution, meaningful only for series that resolved. That leaked the outcome into the predictor: the model separated perfectly (coefficient 17.8, SE 1693) and the enrichment test compared 2 series against 13 on a marker largely encoding a donor field’s *name*. Rebuilt on src/utils/design_features.R, computed from characteristics **keys** and sample titles, which exist for all 51 series.

**The selection mechanism is now evidenced, not asserted.** repeat_design_marker predicts resolvability at **p = 0.009** (coefficient 2.06, SE 0.79). So donor identity does resolve preferentially in series with repeated structure, exactly as the concern held.

**The inflation-difference half is directionally consistent but underpowered:** within the resolved stratum, median inflation 7.44 with a repeat marker (n=12) vs 4.00 without (n=3), Wilcoxon p = 0.27. Direction as predicted, not significant. Reported with both n’s.

Taken together — significant selection, directionally consistent inflation gap, and an IPW estimate below the conditional figure — the evidence supports that 5.13× **overstates** the population rate. Heckman correction not used, per instruction.

**Decision 3 item 5 — does resolvability predict read availability? No.** Reads public for 80% (12/15) of resolvable vs 83% (30/36) of non-resolvable series; Fisher p = 1.00, OR 0.80. The hypothesised compounding effect — that poorly documented studies also deposit reads less often — **is not present in these data**. Direction reported as instructed.

#### D-021 — Tiered donor resolution: 31 unresolved → 23, entirely via SRA titles

**Date:** 2026-08-05 · **Stage:** 1i · **Source:** CP-0b Task A · **Status:** RESOLVED

No published tool groups GEO samples to the same donor, so this was built (src/utils/donor_resolution.R), citing precedent honestly: **GEOracle** (Djordjevic et al., *Methods* 2019, PMID 30959271) groups GSMs into replicate sample groups, and **GAUGE** (*mSystems* 2021, PMC8547006) builds a text-string distance matrix over sample titles. Neither is donor-specific.

| tier | resolved | outcome value |
| --- | --- | --- |
| T1 GEO characteristics donor field | 15 | resolved |
| T2 SRA/BioSample attribute column | **0** | — (see D-018) |
| T3 donor token in SRA experiment/sample title | 12 | resolved_inferred |
| T4 title-stem clustering | 3 | ambiguous (flag, never a pass) |
| none | 24 | unresolved |

Before: 15 resolved / 5 ambiguous / 31 unresolved. After: 27 resolved-or-inferred / 3 ambiguous / 24 unresolved (some ambiguous series were promoted to inferred).

**T3 is where the gain comes from, and it is inference.** GEO characteristics omit donor identity for most series, but SRA experiment titles often carry it verbatim — GSE334185’s "IPF, mock, SEN, donor 110A, batch2" and GSE231693’s "SSc_5444" are both parseable when the GEO record alone is not. Kept as a distinct outcome value and never pooled with T1/T2.

**Three T3 results are flagged as likely parse artifacts** and revert to manual review rather than to a count: GSE146436 (26 samples → 2 units), GSE215274 (177 → 9), GSE294260 (70 → 5). Excluding them, T3 inflation is 3.42× against firm 5.13×.

Two incidental findings from the SRA titles, both material:

- **GSE231693 contains systemic sclerosis samples** (SSc_5444), study title “Evaluation of common profibrotic pathway across interstitial lung disease”. This explains the 20-of-60 ambiguous condition labels seen at Stage 1B and raises whether it belongs in an IPF-specific panel at all. Flagged for the roster review, not decided here.
- **GSE334185’s SRA study title reads** “GEO accession GSE334185 is currently private and is scheduled to be released on Aug 01, 2026.” Independently corroborates its exclusion.

Cross-series patient overlap was **not** attempted: the rigorous method is genotype fingerprinting (CrosscheckFingerprints/Picard, *Nat Commun* 2020) or MD5 redeposit detection (DupChecker, Sheng et al., *BMC Bioinformatics* 2014;15:323), both of which need raw data — precisely what this project shows is usually missing. Available only on the SRA-backed subset. The read-free alternative (expression-profile correlation across series, precedent Toker, Feng & Pavlidis, *F1000Research* 2016) is deferred to the LTRC/Colorado/Yale overlap question at Stage 2.

#### D-022 — Gate 1 RETIRED, not lowered. This is an analytical concession.

**Date:** 2026-08-05 · **Stage:** CP-0b r2 · **Source:** operator Decision 1 · **Status:** RESOLVED

Stated for the record so a reviewer can audit the reasoning rather than inferring the worst:

A threshold move would have lowered the bar and left the analysis plan unchanged. Retiring the gate changes the analysis: the random-effects meta-analysis is abandoned as a primary method, and leave-one-cohort-out is demoted from pooled estimation to a replication check. This is an analytical concession made before results were seen, recorded here together with the admission that the original threshold was set to an expected roster size rather than to an analytical requirement, and was revised three times.

The three revisions were ≥8 (amendment v2.0 §3.4), ≥7 (D-013), and a would-be ≥6 — each proposed after the roster was known. It never functioned as a pre-registered stopping rule, which is why it is removed rather than re-set.

**Replaced by component-level pre-registration**, fixed in config before any concordance result exists:

| Component | n=6 status |
| --- | --- |
| Pairwise cross-cohort concordance (15 pairs, bootstrap CIs) | **PRIMARY — fully supported** |
| Axis 3 analytical-choice factorial | SUPPORTED, wider CIs |
| Core signature by effect-size combination | SUPPORTED |
| Vote-count k-sweep (k=2..6) | CORROBORATION ONLY |
| Axis 2 processing bound | AS AVAILABLE |
| LOCO random-effects meta-analysis | **DEMOTED — not primary** |
| GSE199949 held-out replication | SUPPORTED, power-limited |

**Amended LOCO specification.** At k=5 per iteration, no DerSimonian–Laird or REML pooled estimate is presented as a result. LOCO becomes a *replication* exercise — for each held-out cohort, does the signature from the remaining five show concordant direction and above-chance rank recovery (directional concordance rate, π₁)? This sidesteps τ² entirely. If pooling at all: common-effect model primary with explicit heterogeneity description; random effects with Knapp–Hartung only as caveated sensitivity. τ² reported with Q-profile CIs. **I² banding dropped entirely** — not interpretable at k<10. Methods must state that τ² is poorly estimated at this k and that no random-effects point estimate is authoritative.

**Paper 2 framing:** a cross-cohort DE concordance/robustness benchmark — not a biological meta-analysis, not a pooled effect estimate. Jaccard and directional metrics are reported **together, always**; a moderate Jaccard can coexist with strong directional agreement and reporting one alone invites misreading.

#### D-023 — Unresolved-stratum policy for the audit

**Date:** 2026-08-05 · **Stage:** CP-0b r2 · **Source:** operator Decision 3 · **Status:** RESOLVED

The unresolved rate is treated as a **headline finding**, not a nuisance to be dispositioned. At the IPF scale it is 24/51 = 47% after the Task A pipeline (61% before). Context: Huang et al., *Genome Biology* 2025;26:274 — >25% of critical metadata omitted, 74.8% of relevant phenotypes available, 11.5% of studies shared all phenotypes, 37.9% shared <40%. **Only these four headline figures are taken from that paper; no per-field figure is claimed** (flagged UNVERIFIED per instruction).

Policy, recorded in config.audit.unit_resolution_policy:

1. **Denominator convention** — the full verified set is the denominator and *unassessable* is an explicit reported category, never silently dropped. Precedent: Huang 2025; Ellis et al., *NAR* 2018;46(9):e54 (sex given for only 3,640 of 49,564 = 7.3% of human SRA RNA-seq samples). Ellis’s distinction is adopted: unassessable records are **counted in the missingness statistic** but **excluded from accuracy/derived-quantity calculations**.
2. **PRISMA 2020 flow diagram** with an explicit partition at the unit-resolution step: resolved / ambiguous / unresolved.
3. **ambiguous never passes.** Retained from Task B2.
4. **unresolved enters L1 but is flagged and stratified** — never counted at GSM value, never dropped. Cascade computed primarily within the resolved stratum; full-population cascade reported as bounds.
5. **Resolvability vs read availability tested** — no association (D-020).

#### D-024 — The frozen query template cannot distinguish sibling diseases; 1.5c held

**Date:** 2026-08-05 · **Stage:** 1.5b · **Source:** FREEZE-A query template · **Status:** OPEN — blocks 1.5c, needs an operator decision

Stage 1.5b completed cleanly (1,219 of 1,220 descriptors queried, 0 search failures, L0 = 46,127, **3,903 unique candidate series**, 190 diseases with ≥5 provisional candidates). Two results in the top-15 table are impossible, and the cause is the frozen query template.

**"<term>"[All Fields] does not enforce phrase matching in the gds database.**

| query | L0 |
| --- | --- |
| "Diabetes Insipidus"[All Fields] | 762 |
| "Diabetes Mellitus"[All Fields] | **762 — identical** |
| "Diabetes Insipidus"[MeSH Terms] | 767 |
| "Diabetes Mellitus"[MeSH Terms] | 898 |
| "Idiopathic Pulmonary Fibrosis"[All Fields] | 171 |
| "Idiopathic Pulmonary Fibrosis"[MeSH Terms] | **0** |

Both diabetes descriptors return the same 762 records and the same 118 candidates: the query matches on diabetes and discards the qualifier. Corroborating evidence of imprecision — one candidate series is attributed to **43 different diseases**, 8 series to more than 10, and 6,354 attribution rows cover only 3,903 unique series.

**The obvious repair is worse than the defect.** [MeSH Terms] separates the two diabetes descriptors correctly but returns **0** for idiopathic pulmonary fibrosis, which demonstrably has 171 series and is this project’s anchor disease. GEO DataSets records are not reliably MeSH-indexed, so neither field is safe on its own.

**Why this is load-bearing rather than cosmetic.** Disease retention uses *verified* L1, which protects against most contamination — but the 1.5c verification as written checks tissue, design and case/control counts, **not whether a series actually concerns the disease it was retrieved under**. A diabetes-mellitus series would therefore pass L1 under “Diabetes Insipidus” and count toward its ≥5. Disease selection — the thing FREEZE-A exists to protect — is corrupted unless a disease-relevance check is added. The error also runs in the direction that makes Paper 1’s cascade look **worse** than reality, by inflating L0.

**1.5c is held.** Verifying 3,903 candidates is a multi-hour NCBI run and should not be spent against a denominator known to be wrong.

**Recommended resolution (operator decision required).** Add a *disease-relevance verification layer* applied uniformly to every candidate, recorded as a new cascade sub-layer L0 → L0′, leaving the frozen query and its measured imprecision reported honestly. This preserves FREEZE-A rather than amending it: the frozen template still defines retrieval, the new layer verifies the record against the disease term the same way spec §5.1 verification already checks every hit against its sample table rather than trusting the query, and it applies identically to all 1,220 descriptors so it cannot cherry-pick. The alternative — amending the template and re-freezing — is defensible too, since no L4/L5 coverage has been measured yet, but it would be the second freeze revision and needs to be an explicit operator act.

**Not fixed unilaterally.** config/config.yaml is locked by the FREEZE-A deny rule and CLAUDE.md rule 7 forbids changing a frozen parameter.

#### D-025 — Frozen disease list contains 6 non-English MeSH labels; immaterial but recorded

**Date:** 2026-08-05 · **Stage:** 1.5a · **Source:** MeSH SPARQL enumeration · **Status:** ACCEPTED

The SPARQL enumeration did not filter rdfs:label to English, so MeSH’s translated labels entered the frozen list: 6 of 1,220 descriptors (0.5%), all for tree number **C10.228** (Central Nervous System Diseases) — Portuguese, French, Czech, Croatian, Russian and Japanese.

**Immaterial to every reported number:** all 6 return L0 = 0 and 0 candidates, because GEO is queried in English. Effect is 6 wasted queries and 6 spurious L0-zero rows in the “diseases queried” count.

One consequence worth recording: two of the six (Russian and Japanese) contain no ASCII characters, so both slugified to _ and shared a checkpoint file. That is why 1.5b reports **1,219** descriptors queried against a frozen list of 1,220 — not a dropped disease, a filename collision between two labels of the same tree number, which was already queried successfully under its English label.

**Action:** filter to FILTER(lang(?label) = "en") if the enumeration is ever re-run, and slugify with a hash fallback. Neither is applied now — the frozen list is locked and the effect on results is nil.

#### D-027 — L0 rev2 assembly inflated 1,124× by an unindexed vector; corrected

**Date:** 2026-08-05 · **Stage:** 1.5b2 · **Source:** own defect · **Status:** RESOLVED

src/15b2_screen_revised.R wrote category = dis$category instead of dis$category[i], so every per-descriptor checkpoint stored the full 1,124-element category vector, and data.frame() recycled each row to 1,124. The assembled table reported **1,263,376 descriptors queried** (= 1,124², the signature of the bug), 21,356 search failures and an L0 total of 44,375,520.

The candidate map was **not** affected — it does not use that field — so the accession-level results stand. Corrected assembly, re-derived from the same cached checkpoints:

| descriptors queried | 1,124 |
| --- | --- |
| search failures | 19 |
| descriptors with L0 > 0 | 438 |
| total L0 original template | 39,480 |
| total L0 revised template | 19,885 |
| reduction | **1.99×** |

Fixed in source. The re-assembly drops the stored category entirely and re-joins it from the frozen descriptor list rather than trusting the checkpoint.

#### D-028 — The 3,903-vs-16,820 candidate comparison was not like-for-like

**Date:** 2026-08-05 · **Stage:** 1.5b2 · **Source:** own defect · **Status:** RESOLVED

The revised screen’s fetch_accessions() collected every GSE accession returned, whereas the original screen applied automated filters (single-cell, microarray, non-Illumina, <25 samples) before counting candidates. Comparing 16,820 against 3,903 therefore compared an unfiltered pool with a filtered one and would have overstated the revision’s effect on pool size.

The original filters were applied verbatim to the revised pool. gds UIDs for GSE records are 200000000 + <GSE number> (verified this session: GSE92592 → 200092592), so all 16,820 summaries were retrieved without an esearch round-trip.

| filter | removed |
| --- | --- |
| single-cell / single-nucleus | 2,798 |
| microarray | 178 |
| non-Illumina | 315 |
| < 25 samples | 12,244 |

**Comparable candidate count: 3,631** (original 3,903, −7%).

**The revision’s real effect is on attribution precision, not pool size:**

|  | original | revised |
| --- | --- | --- |
| attributions per series | 1.63 | **1.25** (−23%) |
| max diseases one series | 43 | 32 |
| series attributed to >10 diseases | 8 | 6 |

Pool size barely moves because the revision loses spurious sub-phrase matches while gaining genuine recall from entry terms and de-inverted labels — two effects in opposite directions. Both are reported rather than netted silently.

#### D-029 — GEO rate-limit block killed the first census run

**Date:** 2026-08-05 · **Stage:** 1.5c · **Source:** own defect · **Status:** RESOLVED

The first census run completed exactly **125 series, then failed on all 193 that followed** — consecutive successes then consecutive failures, which is an IP-level block, not random error. Throughput fell from 6 min/100 series to 51 min/100 and the projection diverged from 231 to 1,177 minutes. Killed at 318/3,631. Access recovered on its own once the run stopped, so the block was a resetting window rather than a ban.

Two mistakes, both mine:

1. **Wrong pace.** GEO’s SOFT and acc.cgi endpoints were routed through ncbi_call(), which paces at Entrez’s 0.12 s. Entrez’s 10 req/s allowance does not apply to GEO’s web endpoints. Amendment v1.1 §6.2 warned about exactly this endpoint; I applied the warning to Entrez and left the GEO path unpaced.
2. **Wrong failure behaviour.** Five retries with exponential backoff meant every blocked series burned ~62 s hammering a wall.

The 125 successes averaged **3.0 s apart** and still tripped the limit, so the constraint behaves like a **cumulative quota per window** (~375 requests = 125 series × 3) rather than a rate ceiling. Slowing down alone therefore does not fix it — reducing requests per series does.

**Requests per series cut 3 → 1:**

- Supplementary filenames now come from the SOFT record’s own supplementary_file fields, verified on GSE92592, GSE213001 and GSE231693. One request eliminated outright.
- acc.cgi deferred to pass 2 over only the series that reach L2 or need route B. CLAUDE.md rule 2 requires the literal “Raw data” string for every *accession claim*, and a series failing L1 on tissue or size never becomes one. A present L3 verdict needs no second route since reads demonstrably exist; an absent one stays pending_route_b, so rule 4’s dual-route requirement still holds.

New src/utils/geo_pacer.R paces GEO independently and, on a run of failures, declares a block, **logs the request count where it fell**, sleeps, and resumes — so the quota is measured as a by-product of useful work rather than by deliberately provoking blocks.

**Also caught before restarting:** the 193 failed checkpoints were on disk, and the loop skips anything checkpointed, so they would have been permanently recorded as failures with no retry.

#### D-030 — Every retry loop in the codebase was incapable of retrying

**Date:** 2026-08-05 · **Stage:** 1.5c · **Source:** own defect, found by a unit test · **Status:** RESOLVED

Found while unit-testing the new pacer’s block handler, which is why the test was worth writing.

ncbi_call(), with_retry() and geo_request() all evaluated their argument with **force(expr) inside the retry loop**. In R, a promise that has been forced and thrown is left *under evaluation*, so every subsequent force() on it raises

promise already under evaluation: recursive default argument reference or earlier problems?

instead of re-attempting the request. **Every retry loop therefore performed one genuine attempt followed by N phantom failures.** Retries have never worked in this project.

Consequences:

- The ncbi_call “5 attempts with backoff” was 1 attempt plus 4 phantom errors, and the ~62 s per blocked series in D-029 was time spent sleeping between phantoms. D-029’s diagnosis was right about the symptom and incomplete about the cause.
- In the new pacer this was worse than useless: one real failure produced three “failures”, instantly declaring a block; after the 15-minute sleep the retry raised the phantom error again and blocked again, up to GEO_MAX_BLOCKS — **40 blocks × 15 min = 10 hours of sleeping to give up on a single series.** The test reproduced exactly this, blocking every 3 requests at 76, 79, 82 … 124 with the promise error.

Fixed in all three: the expression is captured with substitute() and re-evaluated with eval(expr, parent.frame()) on each attempt. tests/test_retry.R pins the behaviour — a transient failure recovers with the correct number of *genuine* attempts, no block is declared, the phantom message never appears, and persistent failure still returns NULL after exactly attempts real tries.

**Effect on results already recorded:** none that changes a conclusion, because the three-valued discipline held the line. A failed lookup returned NULL and NULL was always recorded as unresolved, never as absent — CLAUDE.md rules 3 and 4 were written for exactly this class of failure and they did their job. What was lost was recoverable transient failures, which is a recall cost, not a correctness one. The census was restarted from its 468 banked checkpoints on the corrected code.

#### D-031 — Tier B coverage measured: Gemma 5.3%, MetaSRA 7%. The agreement estimate will be thin.

**Date:** 2026-08-05 · **Stage:** 1.5 · **Source:** CP-0b Task 3 validation design · **Status:** ACCEPTED as a limitation (stop rule applies — reported, not redesigned)

config/l0prime.yaml records Gemma’s and MetaSRA’s holdings as UNVERIFIED pending an API check. Both now measured directly.

**Gemma is live and current.** gemma.msl.ubc.ca/rest/v2/datasets/count returns **23,716 datasets**, against the 10,811 in Lim et al., *Database* 2021 — so the publication figure understates it by more than half and should not be cited as current, exactly as the operator instructed.

**Its annotations are rich and fit Tier B’s purpose.** For GSE178518 the API returns EFO/OBO terms: ExperimentTag → idiopathic pulmonary fibrosis (EFO_0000768), fibroblast of lung (CL_0002553), bulk RNA-seq assay, plus FactorValues pirfenidone, saracatinib, nintedanib and TGF-beta.

**But coverage of this candidate pool is 5.3%.** Probing 171 accessions (120 randomly sampled candidates + all 51 verified IPF series) found 9 in Gemma — and only **3 of the 51 IPF series**. GSE92592 is absent ([] on lookup, 404 on annotations). This is the brain-enrichment caveat made quantitative. MetaSRA is comparably thin at 3 of 42 SRP-resolved series (7%, D-019).

**Consequence for the frozen validation design.** Task 3 computes reliability as Tier A ∩ Tier B agreement, with a 0.8 decision threshold. At ~79% Tier A coverage and ~10% combined Tier B coverage, the overlap is roughly 290 of ~4,500 pairs. That is enough for an agreement estimate with confidence intervals, but two caveats must travel with it:

- **It is thin.** ~290 pairs, not thousands.
- **It is non-randomly selected.** Gemma curates to its own priorities (brain-enriched) and MetaSRA depends on SRA presence with ~2019-era coverage. So the agreement estimate’s transportability to the full pool is questionable, and it should be reported as a property of the overlap stratum rather than of the pipeline as a whole.

The 0.8 threshold and its consequence (weight curated sources more heavily if agreement is lower) still stand as frozen. Per the stop rule this is written up as a limitation with its direction and magnitude, not repaired with a sixth round.

**One incidental corroboration worth keeping.** Gemma’s annotation of GSE178518 as fibroblast of lung with drug FactorValues independently agrees with the eligibility filter, which excluded that series on fibroblast + stimulat, tgfb. Together with the 3/3 MetaSRA agreement (D-019), that is a second independent check on the filter’s hard cases.

#### D-032 — The GEO quota is a rolling window across recent activity, not a per-process count

**Date:** 2026-08-05 · **Stage:** 1.5c · **Source:** pacer block log · **Status:** ACCEPTED

D-029 inferred a quota of ~375 GEO requests per window, from 125 series × 3 requests. The pacer’s block log shows that inference was wrong, and the measurement is now direct:

| run | GEO requests before blocking |
| --- | --- |
| original (3 req/series) | ~375 |
| pass 1 | 325, no block, killed for the D-030 fix |
| pass 1b | **blocked at 116** |

Pass 1 reached 325 requests without blocking while pass 1b blocked at 116, so the limit is not a per-process request count. It behaves as a **rolling cumulative window across recent activity from this IP** — combined recent GEO traffic across the three runs is roughly 800 requests.

Practical consequence: the census will proceed stop-start, and no single pacing interval avoids it. The design already handles this correctly — one request per series, block detection, log the boundary, sleep, resume from checkpoint — and every block boundary is now recorded to results/tables/geo_block_log.rds as a reportable measurement about working with GEO at scale, which is itself relevant to Paper 1.

**I am not issuing another wall-clock estimate.** Three have now been wrong (2.3 h, 8 h, then a ~375-request-per-window projection). The block log will report actual throughput.

#### D-033 — Verification abandoned as a census; FREEZE-S two-stratum design

**Date:** 2026-08-05 · **Stage:** 1.5 · **Source:** GEO quota (D-032) · **Status:** RESOLVED

The exhaustive census was abandoned after GEO’s escalating penalty made it unworkable: block 1 at 116 requests, then blocks 2 and 3 after **3 requests each** following 15-minute sleeps. My claim that “the pacer handles it” was wrong.

**The decisive argument for sampling is inferential, not pragmatic: an interrupted census is a biased sample with no weights; a designed sample has known weights.** Whatever GEO happens to let through is non-random and unweightable after the fact.

Frozen as **FREEZE-S 366bdd4**, committed before the draw, in a third separate config so neither existing deny rule had to be lifted.

**ARCHS4 as a metadata substitute — validated before relying on it.** ARCHS4’s meta/samples carries characteristics_ch1, source_name_ch1 and title for all 855,835 samples, readable by HTTP range request with zero GEO load. Against 40 series with banked GEO SOFT records, **eligibility agreement was 39/40 = 97.5%**. The single disagreement (GSE140900) was caused by **incomplete sample coverage, not differing field content** — ARCHS4 held 23 of its 113 samples, so the mouse samples that triggered exclusion were absent. Sample counts matched on only 30 of 40 series, which is why the substitution is gated at ≥90% sample coverage.

**Design and result:**

| stratum | n | verification | GEO cost |
| --- | --- | --- | --- |
| S1 — ARCHS4 ≥90% coverage | 850 | census | **0** |
| S2 — frame | 2,781 | 599 drawn, stratified by MeSH category | 503 fetches |

Sampling fractions 0.188–0.226 across the 14 categories, so weights sit in a narrow 4.43–5.33 band. GEO work fell from 3,049 to 503.

**The 582 banked series are not a random sample** — they are a prefix of processing order, and processing order correlates with GSE-number era, which correlates with SRA deposition practice, which is a cascade outcome. Pooling them would import that bias. Resolution: the S2 draw runs over the full frame regardless of banked status, so caching affects cost only, never selection. 96 draws happened to be banked and are free. The 346 banked-but-undrawn S2 members are a convenience set, reported separately and **excluded from the population estimate**.

#### D-034 — Cascade layers have different data dependencies; L2 is the only GEO-bound one

**Date:** 2026-08-05 · **Stage:** 1.5 · **Source:** own analysis · **Status:** ACCEPTED

Established while resolving D-033, and it materially improves what is achievable:

| layer | requires | censusable now |
| --- | --- | --- |
| L1 design eligibility | per-sample metadata | 1,432 of 3,631 (850 ARCHS4 + 582 banked) |
| **L2 raw count matrix** | **GEO supplementary FILENAMES** | **582 only** |
| L3 public reads | Entrez + ENA | **all 3,631** |
| L4 / L5 resource coverage | local catalogues | **all 3,631** |

**L2 is the single GEO-bound layer.** Distinguishing a raw count matrix from an FPKM matrix needs the filename, and ARCHS4 does not carry supplementary files. Entrez’s gds suppfile field returns only *extensions* — verified: GSE92592 → TXT, GSE213001 → CSV, GSE231693 → TSV — which cannot make the distinction. L2 is therefore estimated from the S2 sample plus the banked series, with its own basis stated rather than being folded into a single denominator.

**Consequence, and it favours the paper:** L3/L4/L5 — the reprocessability layers Paper 1 is actually about — are censusable over all 3,631 candidates at zero GEO cost. The GEO constraint bites on design eligibility and deposition format, not on the reprocessability finding.

Each layer’s denominator and basis must be reported explicitly. A single pooled “N” across layers would misrepresent three different evidentiary bases as one.

#### D-035 — S1 census result: 62% of disease-query hits are not primary-tissue case–control

**Date:** 2026-08-05 · **Stage:** 1.5h · **Source:** S1 census · **Status:** RESOLVED

All 850 S1 members verified, zero GEO requests. **321 of 850 (37.8%) pass the L1 design criteria.** Agreement across sources is close — 38.3% for ARCHS4-derived versus 35.0% for banked GEO SOFT — a further check on the substitution beyond the 97.5% figure.

| exclusion reason | n |
| --- | --- |
| in vitro / cultured **and** perturbation arm | 320 |
| perturbation arm | 104 |
| in vitro **and** perturbation **and** non-human | 56 |
| in vitro / cultured | 35 |
| perturbation **and** non-human | 7 |

**About 62% of series retrieved by a disease query are not untreated primary-tissue case–control studies at all.** This is the GSE188464 failure mode — cultured fibroblasts under drug perturbation passing a naive “primary tissue” filter — quantified at population scale. It is a methods contribution for Paper 1 and it vindicates the operator’s insistence that the screening filter be fixed before FREEZE-A.

**GSM→donor inflation: 3.04×** (18,253 samples → 6,004 units) across the 173 series where donor identity resolved. This independently replicates the IPF-only conditional figure of 5.13× (D-020) on a much larger base, and lands **below** it — consistent with the prediction that the resolved stratum overstates the population rate because donor fields appear preferentially in high-inflation designs. Donor resolution: 173 resolved, 104 ambiguous, 573 unresolved.

#### D-036 — A single corrupt cache file manufactured a fictitious GEO “block”

**Date:** 2026-08-05 · **Stage:** 1.5j · **Source:** own defect · **Status:** RESOLVED

The S2 fetch declared [BLOCK 1] after 3 requests and began a 45-minute sleep. **There was no GEO block.** The cause was one truncated file left in the cache by the earlier throttled run:

data/cache/geo/GSE215886.soft.gz	938 bytes	-> "Found 0 entities" -> NA/NaN argument

GEOquery prefers its local cache, so it re-read the corrupt file three times, parsed nothing each time, and threw. The pacer counted three consecutive failures and inferred throttling. With GEO_MAX_BLOCKS = 12 and a 45-minute wait, that would have burned **9 hours sleeping against a problem that was not rate-limiting at all**.

Two defects, both mine:

1. **Truncated downloads were left in the cache.** Amendment §2.3 says never partially process a truncated object; I honoured that for parsed objects and not for the download cache, so the affected accession was poisoned permanently. paced_getGEO() now purges any .soft* file under 2 KB before each attempt and rejects any object parsing to zero samples, deleting the cache entry so a retry is a genuine refetch. Scope of damage was small — 1 of 641 cached files.
2. **The pacer could not tell a bad series from a rate block.** geo_request() now takes a key. Repeated failure on ONE key means that series is broken: abandon it and move on, without touching the block counter. Only failures spanning DISTINCT keys count as evidence of throttling. Tests in tests/test_retry.R pin both directions — one key failing repeatedly declares no block, three distinct keys failing does.

**A regression caught by an existing test, worth recording.** My first fix set GEO_KEY_ATTEMPTS = 2, which abandons a series on its second failure and so broke genuine transient recovery — the pre-existing “geo_request re-evaluates and recovers from a transient failure” test failed immediately. Raised to 3, which keeps transient recovery while still bounding a bad series. The test written for D-030 paid for itself within the hour.

**Bearing on the earlier probe conclusion.** I reported that the GEO penalty “appears cleared” on 6/6 successful probe requests, then S2 blocked within minutes and I took that as the probe being wrong. **The probe was right.** GEO was not throttling; my cache and pacer were at fault. The correct reading of the evidence was available and I misattributed it — the same error as D-029, where I diagnosed the symptom correctly and the cause incompletely.

#### D-037 — GSE231693 RETAINED; its 20 SSc samples tagged for specificity, not dropped

**Date:** 2026-08-05 · **Stage:** CP-0b r2 · **Source:** operator Decision 2 · **Status:** OPEN — the tagging is a Stage 2 / CP-1 deliverable

Recorded because the ruling had no entry of its own and would otherwise have been lost between CP-0b and Stage 2. GSE231693 appeared in this log only incidentally (as the SSc discovery in D-021 and as a supplementary-file test case in D-029/D-034).

**Two questions were being conflated, and they have different answers.**

*Audit attribution (L0′)* is mechanical: systemic sclerosis sits under MeSH C17, IPF under C08.381.483.652.500. Different branches with no shared path above the root, so tree-number adjudication eliminates cross-attribution by construction. Nothing to decide.

*Roster eligibility (Paper 2)* is independent of the series’ topic. The only question is whether the sample table supplies ≥15 IPF cases and ≥10 non-fibrotic controls at the patient level.

**Ruling: retain.** The 20 SSc samples are excluded from the primary IPF-vs-control contrast and tagged ssc_specificity = TRUE, exactly as GSE150910’s 82 CHP samples are handled (config.benchmark.secondary_analyses.chp_specificity). The consistency argument is decisive: CHP and SSc-ILD are both non-IPF fibrotic ILD arms in an otherwise-qualifying series, and treating them differently without a principled distinction would be the arbitrary act.

**Verification performed before the ruling stands** — the operator required confirmation that the controls are genuinely non-fibrotic rather than SSc-without-ILD, since a series involving systemic sclerosis could plausibly use scleroderma patients lacking lung involvement as “controls”. Read from the sample table, all 60 GSMs, verbatim:

| arm | n | verbatim characteristics |
| --- | --- | --- |
| IPF | 20 | tissue: lung \| disease state: idiopathic pulmonary fibrosis |
| **Control** | **20** | **tissue: lung \| disease state: normal** |
| SSc | 20 | tissue: lung \| disease state: systemic sclerosis |

source_name_ch1 is lung for all 60; title stems are IPF / Normal / SSc, 20 each. The controls are **normal lung, not SSc-without-ILD**, so the cohort clears ≥10 controls at the patient level (60 GSMs = 60 patients, no repeated sampling). **n = 6 stands**; the component scope pre-registered for n=6 (D-022) does not need revisiting.

**Deferred deliverable.** The ssc_specificity tag belongs in metadata/cohort_labels.csv, which is spec §6 — Stage 2, gated by CP-1. It is deliberately NOT created now. This entry is the carrier for the decision until then. Note also that config/config.yaml is deny-locked by FREEZE-A, so the declaration cannot be added there; when the CP-2 freeze of the benchmark plan happens, ssc_specificity should be recorded alongside chp_specificity as a declared secondary analysis.

#### D-038 — Paper 1’s core result: the reprocessability cascade

**Date:** 2026-08-05 · **Stage:** 1.5c–1.5i · **Status:** RESOLVED (L3/L4/L5 complete)

**L3/L4/L5 censused over all 3,631 candidates, 0 failures.** L1 and L2 rest on the FREEZE-S design because they are metadata- and GEO-bound respectively (D-034).

| layer | rate | basis |
| --- | --- | --- |
| L0 enumerated | 3,631 | exhaustive over 1,124 frozen MeSH descriptors |
| L1 eligible (weighted) | **24.7%** | S1 census (850) + S2 sample (598 of 599) × weights |
| L2 raw count matrix | 40.1% | 1,084 GEO-verified |
| L2 normalized-only | 8.8% | 1,084 |
| L3 public reads | **93.5%** | 3,631 census |
| L4 in ≥1 uniform resource | **50.7%** | 3,631 census |
| L5 usably covered (≥90% samples) | **27.7%** | 3,631 census |

**Headline: of the 3,394 series with public reads, 1,003 are usably covered — 29.6%. So 70.4% of read-public series are not usably reprocessed by any of the three resources.**

Per-resource, usable coverage at the frozen ≥90% gate:

| resource | present | usable |
| --- | --- | --- |
| ARCHS4 | 33.8% | **23.4%** |
| recount3 | 13.4% | 9.1% |
| DEE2 | 33.8% | 3.6% |

DEE2’s collapse from 33.8% present to 3.6% usable is the D-005 finding at population scale: presence is not usability once its own QC flags are honoured.

**L3 absences are bounded, not a point count.** 232 series are route-A absent and 5 unresolved. CLAUDE.md rule 4 requires dual-route agreement before recording reads-absent, so these are reported as a range pending the verbatim GEO record, never as a settled figure.

**The reportable framing.** S1’s L5 is 100% *by construction* — its definition (ARCHS4 ≥90% coverage) is the ARCHS4 L5 criterion, so quoting L4/L5 for S1, or for S1 pooled unweighted with S2, is circular. Outside that self-selected stratum, **S2 alone shows L5 = 5.7%**: roughly one series in twenty has usable uniform reprocessing available.

**Provisional figures reported earlier were biased optimistic, as predicted.** At 33% completion the subset over-represented S1 (26.0% vs 23.4% of the pool, and larger n_gsm at p=0.001), so:

|  | partial | final | direction |
| --- | --- | --- | --- |
| L1 weighted | 31.5% | 24.7% | worse |
| headline drop | 65.8% | 70.4% | worse |

Both moved against the finding by ~6 points. The drift check (D-020’s method applied to processing order) called the direction correctly before the data landed.

**GEO throughput, for the methods section.** The S2 fetch completed 503 series in 492 GEO requests over 81 minutes with **zero blocks**. Every earlier block was attributable to a single truncated cache file and a pacer that could not distinguish a broken series from a rate limit (D-036), not to GEO throttling. The successive wall-clock estimates I gave — 2.3 h, 8 h, a ~375-request quota, then “multi-day” — were all wrong, and the correct figure was only obtainable by instrumenting and measuring.

#### D-039 — The ≥300 L1-series target is RETIRED, not missed

**Date:** 2026-08-05 · **Stage:** 1.5 · **Source:** operator decision · **Status:** RESOLVED

Amendment v2.0 §2.3 set a target of ≥300 L1-qualifying series. The weighted estimate is **147** across **15 diseases in 9 MeSH categories**. The ≥10-disease target is met; the ≥300-series target is not.

**The target is retired, and the operator states the reason plainly: it has the same defect as Gate 1 at ≥8.** It was set before any measurement existed, as a plausible-sounding scale target rather than something derived from a power calculation or a precedent. It was never a validity threshold. This is the second pre-measurement threshold in this project to be retired for exactly that reason (see D-013, D-022).

**Defensible on its merits, not only by analogy.** At n=147 a proportion near 0.25 carries a 95% CI of roughly ±7 points; at n=300 it would be ±5. Wider, not categorically different, and fully reportable. What actually answers “is this general or IPF-specific” is between-disease variance across 15 diseases in 9 categories, which is driven by disease count — the target that was met. Direct precedent exists in this literature: Huang et al., *Genome Biology* 2025;26:274 made population claims about GEO metadata completeness from **253 studies**, the same order of magnitude, recent, high-profile venue.

**147 is not a shortfall to be excused — it is the thesis, measured.** Of candidates a naive automated disease screen returns: **~62% fail L1 on design** (cultured cells, perturbation arms, non-human — D-035) and **40.3% of the survivors are refuted on attribution** by L0′. Compounded, **roughly 23% of what an automated screen calls a qualifying disease study actually is one.** That is the paper’s central claim, now quantified across 15 diseases rather than one. *The reason 300 qualifying series cannot be reached is the reason the paper exists.* It leads.

**Options rejected, with reasons recorded so the git history reads correctly:**

- *Widen the enumeration to depth 4 or more categories.* **Firm no.** The FREEZE-A revision 2 query fix corrected a documented Entrez defect — the template did not do what it was written to do. Widening depth to reach a number is chasing the target, it is visible in the git history, and a reviewer would read it exactly that way.
- *Enlarge S2.* Improves precision on the weighted total, which is not the constraint. It does not change the underlying L1 population.

**For the Discussion, at the operator’s suggestion.** Every threshold set before measurement in this project — ≥8 cohorts, ≥300 series — was wrong in the same direction, because it estimated yields for a population whose properties this study was built to discover. That is not carelessness; it is the structural error any competent investigator would make when planning a reuse study on public transcriptomic data, and it is evidence for the thesis rather than an embarrassment. One sentence noting that the study’s own scoping assumptions failed in the direction the paper predicts belongs in the Discussion.

#### D-040 — Depth-stratified sub-strata: IPF does not clear the threshold, and that is reported

**Date:** 2026-08-05 · **Stage:** 1.5f · **Source:** operator Decision 4 · **Status:** RESOLVED

Of the 15 weighted-retained descriptors, exactly **one** descendant cleared ≥5 L1 independently: **Dermatitis → Dermatitis, Atopic** (n=5, L2=0, L3_present=0, L5=0).

**IPF (D054990) did not clear the threshold as a sub-stratum, and is reported as not clearing it.** It was not promoted to make the typicality comparison possible — which is exactly what Decision 4 was designed to prevent. So “is IPF typical or an outlier” **cannot be answered from the sub-strata distribution** with the frozen enumeration, and that is the honest outcome rather than a gap to be filled by special pleading.

Recorded as the limitation l0prime.yaml anticipated: the frozen enumeration measures at MeSH depth 2–3, i.e. broad disease categories, whereas research practice targets specific entities. The cascade is therefore measured at a coarser grain than the literature operates at. The sub-strata table was the designed mitigation, and at this L1 yield it is too thin to mitigate much — one sub-stratum, itself with zero L2/L3/L5.

#### D-041 — Tier A/B “agreement” measures a different construct than Tier A; reported as such

**Date:** 2026-08-05 · **Stage:** 1.5l · **Source:** own analysis · **Status:** ACCEPTED as a limitation (stop rule applies)

Gemma coverage of the full candidate pool, probed exhaustively: **263 of 3,631 = 7.2%** — above the 5.3% estimated from the 171-accession sample (D-031), and the earlier n=5 overlap was bounded by how many accessions I had checked rather than by Gemma’s coverage.

Overlap 263 series, 227 comparable. **Raw concordance 137/227 = 60.4% (95% CI 53.7–66.8%)**, below l0prime.yaml’s 0.80 threshold, 90 disagreements.

**This is NOT the validation l0prime.yaml specified, and must not be reported as it.** The two tiers answer different questions:

- **Tier A:** does the linked article carry a MeSH heading that IS descriptor D or a descendant?
- **My Tier B test:** does Gemma attach *any* disease ontology term (EFO/DOID/MONDO/HP)?

|  | Gemma disease-annotated | no disease term |
| --- | --- | --- |
| Tier A confirmed | 107 | 33 |
| Tier A **refuted** | **57** | 30 |

The 57-cell is the tell: a series correctly refuted for, say, “Diabetes Mellitus” can legitimately carry an EFO term for a different disease. Under a construct mismatch those are expected, not Tier A errors, so **60.4% is a floor on descriptor-level agreement, not an estimate of it** — the true figure is higher by an unknown amount.

l0prime.yaml anticipated the requirement (crosswalk: "DO -> MeSH via UMLS/MONDO") and that crosswalk was **not built**. Doing so now would be a sixth round of construction, which the operator’s stop rule forecloses. Recorded as a limitation with its direction and magnitude instead:

- **Direction:** the reported concordance UNDERSTATES descriptor-level agreement.
- **Magnitude:** up to 57 of 90 disagreements (63%) are attributable to the mismatch rather than to error, so descriptor-level agreement plausibly lies between 60.4% and 85.5% ((137+57)/227). The 0.80 threshold sits inside that interval, so **the threshold decision is genuinely undetermined by this evidence** — which is the honest statement.
- **Conditional, per D-020’s discipline:** Gemma is brain-enriched and curates a subset, so the overlap stratum is not a random slice and none of this generalises unbounded.

**A further cascade finding, not a methods footnote (operator’s framing).** The best expert-curated GEO annotation resource covers **7.2%** of these candidates and MetaSRA **7%** (3 of 42 SRP-resolved, D-019). The resources that would let a reader verify attribution are as unavailable as the reads (L3) and the donor metadata (D-018). That belongs in the cascade alongside L1–L5.

#### D-042 — The script that builds the audited population was never committed

**Date:** 2026-08-05 · **Stage:** 1.5c0 · **Source:** own defect · **Status:** RESOLVED

data/cache/rev2_census_input.rds defines which 3,631 series are audited. It is read by 15c_census.R:43, 15e_l0prime_tierA.R:31, 15g_draw_sample.R:29 and 15i_l3_l5_census.R:36, and it was **written by no committed script**. The same held for results/tables/audit_rev2_candidate_summaries.csv. The single most important derivation in the study — which series are in the population — was undocumented, and a reader with the deposit could not regenerate it.

**How it was found.** An independent line-by-line audit of the archive against the manuscript traced every reader of the file and found no writer.

**Fix.** src/15c0_build_census_input.R. It reads audit_candidate_map_rev2.csv, takes the unique accessions in retrieval order, pulls the gds esummary for each (from data/cache/rev2_summaries.rds, or from Entrez under --from-scratch), applies the screen from src/utils/screening.R, and writes both files plus results/tables/l0_attrition.csv.

**Numerically.** Nothing changed. The rebuild reproduces the archived population exactly, and the script asserts it (stopifnot(identical(sort(new), sort(old)))) rather than trusting it:

```
16,820  unique GSE returned by the revised retrieval
-2,798  is_single_cell
  -174  is_array          (among survivors)
  -250  non_illumina      (among survivors)
-9,967  too_small (n_samples < 25)
------
 3,631  candidate == TRUE
```

**Pinned by.** tests/test_population.R, and the stopifnot inside the script itself.

#### D-043 — Screening thresholds were implemented in code, not in any config

**Date:** 2026-08-05 · **Stage:** 1.5b / 1.5c0 · **Source:** own defect · **Status:** RESOLVED

The ≥25-sample threshold removes **9,967 series — more than every other filter combined** — and lived as a literal in src/15b_audit_l0_l1.R. It appeared in none of the three frozen configs, while the manuscript presents the analysis as governed by three pre-registered plans. The three screening regexes had the same problem, and were additionally applied in a second, uncommitted place (D-042), so the rule had two copies and one definition.

**Fix.** config/screening.yaml, which opens with POST-HOC DECLARATION, NOT A PRE-REGISTRATION and carries no freeze and no deny rule. The regexes moved to src/utils/screening.R and are now sourced by 15b and 15c0 alike. The original literal is retained as a comment at each site so the diff shows the value did not move.

**Numerically.** Nothing changed. min_samples: 25 is the value that was applied.

**Consequence for the manuscript.** Every rate in the audit is conditional on this threshold, and that conditioning must be stated. Task 3.4 (D-056) measures the cascade on a sample of the excluded population so the conditioning is bounded rather than merely disclosed.

#### D-044 — recount3 coverage was a ratio between two different things

**Date:** 2026-08-05 · **Stage:** 1.5i/1.5i3 · **Source:** own defect · **Status:** RESOLVED

src/15i_l3_l5_census.R:88 computed recount3_cov = frac(r$n_usable), i.e. n_usable / n_gsm, where n_usable came from pipelines.R:77 as sum(as.integer(hit$n_samples)) — the sample count of the matched recount3 **project** — while n_gsm is the sample count of the **GEO series**. Different units on the two sides of a proportion. When one SRP spans several series the ratio exceeds 1.

**Magnitude.** 79 of the 330 series counted recount3-usable (24%) had coverage > 1, to a maximum of **61.44** (GSE23316, 36 GSM). For those 79 the ≥90% criterion was never evaluated against the series’ own samples at all.

**Fix.** src/utils/runlevel.R + src/15i3_recount3_runlevel.R. Coverage is now a **run-level intersection**, |series runs ∩ resource runs| / |series runs|, bounded in [0,1] by construction and asserted. The series’ runs come from elink(gds→sra) + esummary, which returns that series’ SRA experiments and no others; recount3’s runs come from each project’s sra.sra.<PROJECT>.MD.gz metadata (external_id). 464 project manifests, 1,396 series.

**What the fix revealed, beyond capping the value.** GSE23316’s 45 runs are **not present in recount3’s SRP013565 at all** — that SRP is ENCODE’s PRJNA30709, and the intersection is empty. The 61.44 was not merely unbounded, it was spurious: the series is not covered.

**DEE2** had the same unit mismatch (QC-passing *runs* over *samples*), though it never exceeded 1 empirically. It is now run-level where the series’ run list resolves. Where it does not, and only where DEE2 attributes runs to the GEO series **directly** via its GEO_series column, QC-PASS-runs-per-sample is used as a labelled fallback (dee2_cov_basis), because on that route the units are matched-ish and the ratio is empirically bounded at 1. recount3 gets no such fallback — its project-level ratio is precisely the unbounded quantity at issue.

**Numerically:**

|  | before | after |
| --- | --- | --- |
| recount3 usable | 330 (9.088%) | **316 (8.703%)** |
| DEE2 usable | 129 (3.553%) | **123 (3.388%)** |
| L4 present in ≥1 | 1,840 (50.675%) | **1,697 (46.736%)** |
| L5 usably covered | 1,005 (27.678%) | **991 (27.293%)** |
| L5 ∩ read-public | 1,003 (29.552%) | **989 (29.140%)** |
| read-public NOT usable | 70.448% | **70.860%** |

**Pinned by.** tests/test_coverage_bounds.R — every *_cov is NA or in [0,1].

#### D-045 — Unresolvable compendium lookups were collapsed to absent

**Date:** 2026-08-05 · **Stage:** 1.5i · **Source:** own defect · **Status:** RESOLVED

src/15i_l3_l5_census.R:91-94:

l4 = (isTRUE(r$n_usable > 0) || isTRUE(dd$n_usable > 0) || a4_n > 0),
l5 = (isTRUE(frac(r$n_usable) >= COVMIN) || ...)

isTRUE(NA > 0) is FALSE. Every lookup that could not be performed was therefore counted as **not covered**, in a study whose Methods states “an unresolved outcome is never collapsed to absent” and whose CLAUDE.md rule 3 forbids exactly this.

**Magnitude.** 242 series unresolved for recount3, 238 for DEE2, 2 for ARCHS4. Of the 1,791 series recorded absent from all three, **238 (13.3%)** rested on at least one lookup that was never performed. After the run-level rework the unassessable counts are 421 / 381 / 2, and 380 all-absent series rest on an unperformed lookup.

**Fix.** Three-valued per-resource status (covered / not_covered / unassessable) and bounded layer summaries. New columns in l3_l5_census.csv: recount3_status, dee2_status, archs4_status, l4_lower, l4_upper, l5_lower, l5_upper, n_unassessable.

**Numerically** — the point estimate becomes the lower bound, so the headline does not weaken:

| layer | reported as |
| --- | --- |
| L4 | **46.736%** (bounds 46.736–57.202%) |
| L5 | **27.293%** (bounds 27.293–38.777%) |
| read-public not usably covered | **70.860%** (bounds 65.498–70.860%) |

**Pinned by.** tests/test_three_valued.R

#### D-046 — evaluate_l1() was called without donor identifiers

**Date:** 2026-08-05 · **Stage:** 1.5d/1.5m · **Source:** own defect · **Status:** RESOLVED

This is the most damaging finding in the audit, because it is self-undermining.

src/utils/casecontrol.R::evaluate_l1() supports donor-level counting, but only when passed unit_ids with unit_status %in% c("resolved","resolved_inferred"). At src/15d_cascade.R:222 it was called as ev <- evaluate_l1(b, alts) — no unit_ids, no unit_status. The defaults are NULL and "unresolved", so the donor branch was **never taken** and counted_on was "samples" for every pair in the audit.

**The paper’s fifth headline result is that sample accessions overstate biological replication by ~3×. The L1 layer then counted sample accessions.**

ev$status and ev$counted_on were also discarded rather than persisted, which is precisely why the substitution was invisible.

**Fix.** src/15m_donor_ids.R writes results/tables/donor_ids_by_series.csv from records already on disk — **no network calls**. Two sources: the cached GEO SOFT checkpoints, which had been storing unit_ids all along, and ARCHS4 sample metadata for series never fetched from GEO. 15d_cascade.R now passes both vectors and persists status, counted_on, counted_on_donor and donor_counted into audit_pair_level.csv.

**Numerically.** 2,055 of 3,631 series have sample text; **293 are usable for donor counting** (143 from GEO SOFT, **150 newly resolvable from ARCHS4 metadata at no request cost**). 315 of 1,835 pairs are now counted on biological units. Across the resolvable stratum, 43,238 sample accessions correspond to 10,138 distinct donors — **4.26 accessions per donor**.

**Pinned by.** tests/test_l1_layers.R

#### D-047 — The reported L1 rate was design eligibility only

**Date:** 2026-08-05 · **Stage:** 1.5d · **Source:** own defect · **Status:** RESOLVED

The reported **L1 = 24.7%** is computed at 15d_cascade.R:112-128 from the eligible column **alone**. The size criterion (≥15 cases / ≥10 controls) is combined only at line 234, after the headline rate has been printed. The number the manuscript calls L1 is therefore not the quantity the manuscript defines L1 to be.

**Fix.** Three layers, all emitted to results/tables/l1_layers.csv, weighted and unweighted, per stratum and pooled:

| layer | definition | denominator |
| --- | --- | --- |
| **L1a** | design eligibility | full weighted frame |
| **L1b** | L1a **and** ≥15/≥10 on sample accessions | full weighted frame |
| **L1c** | L1a **and** ≥15/≥10 on **distinct donors** | donor-resolvable stratum, conditional |

**Numerically:**

|  | weighted | unweighted |
| --- | --- | --- |
| L1a — design eligibility only (**the reported 24.7%**) | **24.735%** | 30.732% |
| L1b — design **and** ≥15/≥10 (what the manuscript defines L1 as) | **7.404%** | 9.461% |
| L1c — counted on donors, resolvable stratum (n=229) | **3.535%** | 3.493% |
| L1b restricted to that same stratum | 13.895% | 13.537% |

**The sentence this licenses, and the reason the task mattered:**

Among the 229 series where donor identity resolves, **3.5%** pass the replication criterion when counted on donors versus **13.9%** when counted on sample accessions — a **3.93-fold** difference.

That is the paper’s own thesis, measured on its own data, and it sits within the range of the independently derived inflation estimates (3.04×–4.60×). The defect becomes the most concrete demonstration in the paper. **L1c is conditional and its denominator is small — 229 series, 6.3% of the audited population — and it must be reported as conditional, exactly as the inflation statistic is.**

#### D-048 — Route B was never completed; the Methods described it as performed

**Date:** 2026-08-05 · **Stage:** 1.5i2 · **Source:** scope · **Status:** RESOLVED

15i_l3_l5_census.R states in its own header: “L3 here is ROUTE A ONLY,” and absent verdicts stayed pending_route_b. The Methods says:

“A series was recorded as reads-absent only when an independent programmatic route and a direct reading of the primary record agreed; disagreements were logged and adjudicated manually.”

**No series had ever been dual-route verified.** CLAUDE.md rule 4 requires it; rule 5 requires that L3a / L3b / L3c never be pooled.

**Fix.** src/15i2_l3_route_b.R fetches the verbatim GEO record for all 237 series route A did not settle (232 absent, 5 unresolved), via acc.cgi, paced by geo_pacer.R. 106 live requests, 2.6 minutes.

**What it found — this is a result, not housekeeping.**

| route A | route B absent | no status line | route B **present** |
| --- | --- | --- | --- |
| absent | 174 | 36 | **22** |
| unresolved | 0 | 3 | **2** |

**24 of 237 route-A negatives are contradicted by the primary record**, twenty-two of them reading verbatim *“Raw data are available in SRA”*. Route A alone has a false-negative rate of roughly 9% on the set it fails to resolve.

**L3 is therefore revised upward**, and both figures are retained in l3_l5_census.csv (l3_present = route A, l3_present_dual = adjudicated) so the archived value stays traceable:

|  | before | after |
| --- | --- | --- |
| L3 reads public | 3,394 (93.473%) | **3,418 (94.134%)** |

**The absence classes, never pooled:**

| class | n | meaning |
| --- | --- | --- |
| **L3a** | 28 | submitter explicitly withholds the reads |
| **L3b** | 50 | controlled access (dbGaP/EGA), recorded never accessed |
| **L3c** | 96 | no evidence of deposition and no withholding statement |
| no status line | 39 | the record renders no Raw data field — the submitter made no claim |

The 39 with no status line are a distinct finding in their own right: amendment §2.4 rests the evidence on that line, and for 39 series it does not exist.

**The two series that prove route A alone is insufficient.** GSE79210 and GSE84023 are usably covered by DEE2 — so their reads were demonstrably processed — while route A returned no_records_found and unresolved. Route B returns present for both. They are also why L5 across the census (991) exceeds L5 within read-public (989). This is a finding about **discoverability**, not an embarrassment: a compendium processed reads that the programmatic route could not find. A broader version holds for 39 series where DEE2 holds QC-PASS runs while elink(gds→sra) returns no SRA records at all.

#### D-049 — Inflation point estimate, bounds and IPW came from three different populations

**Date:** 2026-08-05 · **Stage:** 1.5n · **Source:** own defect · **Status:** RESOLVED

The manuscript reads: “accessions exceeded distinct donors by 3.04-fold in a census stratum, with population bounds of 1.32–10.57-fold and an inverse-probability-weighted sensitivity estimate of 4.60-fold.” Those four numbers describe **three different populations**:

| statistic | value | actual source |
| --- | --- | --- |
| 3.04× | 18,253 samples → 6,004 units | 173 resolvable series in the **850-series S1 census** (D-035; in no result table) |
| conditional 5.13× |  | 15 resolvable series in the **51-series IPF roster** |
| bounds 1.32–10.57× |  | the same 15-series roster |
| IPW 4.60× |  | the same 15-series roster |

src/01h_inflation_bounds.R reads rescreen_corrected.csv, which has 51 rows. **The bounds do not bracket 3.04**, because they are bounds on a different quantity. A reader who checks will find an interval that excludes the point estimate it appears to accompany.

**Fix.** src/15n_inflation_census.R recomputes all four on the census frame from donor_ids_by_series.csv, and adds three things the original lacked.

|  | 51-series roster | census frame |
| --- | --- | --- |
| conditional | 5.133× | **2.650×** (n = 247 resolvable) |
| population bounds | 1.318–10.571× | **1.089–10.768×** |
| Imbens–Manski 95% CI | — | **1.065–11.882×** |
| IPW sensitivity | 4.599× | **2.673×** |
| n series | 51 | 2,055 |

1. **Imbens–Manski confidence intervals** (2004, *Econometrica* 72:1845), with bootstrap standard errors over series (B = 2,000, seeded). These cover the **parameter** with 95% probability, not the identified set; the distinction is stated because it changes the interpretation.
2. **The enrichment test is reported, not asserted.** The manuscript argues that donor fields appear preferentially in high-inflation repeat-design series. On the census frame: median inflation 3.667 with a repeat-design marker (n = 101) versus 2.734 without (n = 146), Wilcoxon W = 8321.5, **p = 0.086**. The direction supports the selection argument; the significance does not, at α = 0.05. Reported both ways.
3. **The propensity model is fully stated.** It has FOUR covariates — log(n_samples) + n_char_keys + repeat_design_marker + title_repeat_frac — while the Methods describes only two.

**The Fisher test, now on the census frame:** reads are public for **91.1%** of donor-resolvable series versus **97.1%** of non-resolvable ones (p = 3.2 × 10⁻⁵, OR = 0.309). The direction is the opposite of convenient and is reported as found.

The 51-series roster results are **retained** in results/tables/inflation_bounds.csv as a legitimate smaller-sample comparison. They must never again be quoted beside a census figure.

#### D-050 — Retained diseases span ten MeSH categories, not nine

**Date:** 2026-08-05 · **Stage:** 1.5k · **Source:** own defect · **Status:** RESOLVED

The manuscript and docs/CP0b_REPORT.md say the retained diseases span **nine** MeSH categories, listing C01, C04, C05, C06, C08, C10, C14, C15, C19. Recomputed from audit_retention_weighted.csv under the weighted retention rule: **ten**, including **C17** (Dermatitis, weighted L1 5.00). The list had been transcribed by hand from a printed table.

**Fix.** src/15k_weighted_retention.R now generates results/tables/retention_categories.csv for both retention rules and prints a diff against the previously reported nine. The list is never retyped.

**Also corrected here (the 1,005 vs 1,003 reporting).** Both figures are correct and the manuscript states only 1,003, so 27.7% cannot be reconciled by a reader multiplying by 3,631. 15i_l3_l5_census.R now emits both explicitly plus the gap: post-fix, L5 across the census is **991**, L5 within read-public is **989**, and the difference is **2** series — GSE79210 and GSE84023, covered by DEE2 while route A found no reads.

#### D-051 — MetaSRA coverage was quoted on a 42-study basis beside a census figure

**Date:** 2026-08-05 · **Stage:** 1.5r · **Source:** own defect · **Status:** RESOLVED

The manuscript reads “Gemma covered 7.2% of candidate series and MetaSRA 7%.”

Gemma’s figure is real and census-scale: 263 of 3,631 = 7.24%, from an exhaustive probe. **MetaSRA’s was 3 of 42** SRP-resolved studies in the IPF roster (D-019; metasra_validation.csv has 51 rows). Two near-identical percentages computed on bases three orders of magnitude apart read as corroboration when they are coincidence.

**Fix.** src/15r_metasra_census.R probes MetaSRA for every series with a resolvable SRP — 3,270 distinct SRPs, 31 minutes — and reports the result three-valued, as everywhere else: a non-empty response is present, an empty response is absent, a failed request is unresolved and is never counted as absence, and a series with no resolvable SRP is a fourth state reported separately rather than folded into either.

| basis | covered | denominator | rate |
| --- | --- | --- | --- |
| Gemma, census probe | 263 | 3,631 | 7.243% |
| **MetaSRA, census probe** | **341** | **3,631** | **9.391%** (bounds 9.391–16.965%) |
| MetaSRA, SRP-resolvable subset | 341 | 3,389 | 10.062% |
| ~~MetaSRA, 51-series IPF roster~~ | 3 | 51 | 5.882% — SUPERSEDED |

Status over the census: 341 present · 3,015 absent · 33 unresolved (request failed) · 242 no resolvable SRP.

**The correction changes the comparison’s direction, not just its precision.** At census scale MetaSRA covers **more** of this population than Gemma (9.4% versus 7.2%), where the manuscript implies parity at about 7%. The sentence must be rewritten around the census figures, and the 42-study number must never appear beside a census number again.

**The cascade finding is unchanged and, if anything, sharpened.** The two best-known annotation resources for GEO/SRA between them reach a small minority of this population. The resources that would let a reader verify attribution are about as unavailable as the reads (L3) and the donor metadata (D-018), and that belongs in the cascade alongside L1–L5.

**Host note, carried forward from src/utils/metasra.R.** The host named in most documentation, metasra.biostat.wisc.edu, is dead — DNS resolves but ports 80 and 443 refuse connections. The service has moved to metasra.platformx.wisc.edu. Querying the old host returns a silent “resource unavailable” and would have produced a census figure of zero.

#### D-052 — The case-by-exclusion rule was undisclosed

**Date:** 2026-08-05 · **Stage:** 1.5q · **Source:** disclosure · **Status:** RESOLVED

classify_samples() defaults to case_by_exclusion = TRUE: every sample is a **case** unless it matches the control lexicon, matches a third-group lexicon, or carries under 8 characters of text. The disease-term match is_dis is computed in that branch and then never used.

The code comment is candid about why — at MeSH depth 2–3 literal matching “collapsed L1 to 50 pairs of 1,835 and retained no disease at all” — and the choice is defensible, since L0′ Tier A establishes the disease attribution at series level. It is disclosed **nowhere** in the manuscript, which promises an Additional file 5 that does not exist.

Supporting numbers, sample-counted as the audit computed them: mean **85.4** cases per pair, max **5,550**, and **47.2%** of pairs with zero detected controls.

**Fix.** src/15q_case_definition_sensitivity.R → l1_sensitivity_case_definition.csv (Additional file 5). Both arms count on the same basis, so the contrast isolates the case definition.

| quantity | case_by_exclusion | literal match |
| --- | --- | --- |
| pairs passing the size criterion | 493 | 136 |
| L1 pairs (design **and** size) | 166 | 44 |
| L1 pairs, attribution-confirmed | 77 | 24 |
| series qualifying at L1 | 114 | 36 |
| **diseases retained, strict** | **2** | **0** |
| **diseases retained, inclusive** | **3** | **0** |
| mean cases per pair | 69.0 | 15.1 |

**Under literal descriptor matching the layer does not merely shrink — no disease is retained at all.** That confirms the code comment’s account and is the honest justification for the rule. The manuscript must state the rule, the reason, and this table.

#### D-053 — Config deny rules were weaker than “literally unmodifiable”

**Date:** 2026-08-05 · **Stage:** 1.3 · **Source:** own defect · **Status:** RESOLVED

.claude/settings.json denied Edit(/config/config.yaml) and the two other frozen configs, and commit 4d3d31c described this as making the frozen rule “literally unmodifiable for the rest of the audit.” Three things are wrong with that claim.

1. A permission rule binds only a Claude Code session whose project root is this repository. **Verified empirically during remediation:** an edit attempted against ipf-repro/config/config.yaml from a session rooted at the parent directory was *not* denied — it reached the string-matching stage and failed only because the probe string was deliberately absent. The rule is real but its scope is narrower than stated.
2. It is a property of one working copy, not of the archive. It travels with the deposit as an inert JSON file.
3. A reader cannot check it. That is the disqualifying problem.

**Fix.** Two parts. - The deny list now carries every path form (/config/…, ./config/…, config/…, **/config/…) so it binds regardless of which root a session uses. - tests/test_freeze_integrity.R provides the evidence a reader can actually verify, from git.

**What the test found, which the plan for this fix did not anticipate.** The intended assertion was “no commit after the freeze touched the file.” That is false for config/config.yaml: commit 4d3d31c came *after* freeze 0b10885 and modified it — to write the freeze hash into its own header comment. No parameter changed. The test therefore compares **parsed YAML** at the freeze commit against HEAD, which is the claim that actually matters, and records the byte-level difference separately.

| freeze | file | commit | commits after | parsed content identical | bytes identical |
| --- | --- | --- | --- | --- | --- |
| FREEZE-A | config/config.yaml | 0b10885 | 1 | **TRUE** | FALSE |
| FREEZE-L0P | config/l0prime.yaml | 03916e7 | 0 | **TRUE** | TRUE |
| FREEZE-S | config/sampling.yaml | 366bdd4 | 0 | **TRUE** | TRUE |

**Manuscript.** Replace “literally unmodifiable” with the git claim: no pre-registered parameter changed after its freeze commit, verifiable from the deposit by tests/test_freeze_integrity.R. Task 3.5 anchors the freeze *times* externally, which git alone cannot establish.

**Pinned by.** tests/test_freeze_integrity.R · results/tables/freeze_integrity.csv

#### D-054 — The IPW estimate was not reproducible across two runs

**Date:** 2026-08-05 · **Stage:** 1h / 1.5n · **Source:** own defect · **Status:** RESOLVED

STATUS.md logs two runs of 01h_inflation_bounds.R **54 seconds apart** reporting IPW **5.09×** and then **4.60×**, with the conditional estimate and both bounds identical in each. A reviewer who spots two values for one statistic will ask.

**Investigated, and the honest answer is bounded.** With the archived inputs the estimator is now **deterministic**: repeated runs return 4.599× every time. The glm converges in 4 iterations, fitted propensities span 0.091–0.811 so no value touches the 0.02/0.98 clip, and nothing in the path is stochastic. Deleting and regenerating design_features.csv — the one input the first run creates and the second reads — reproduces it byte-identically and does not change the estimate. **The input state that produced 5.09 no longer exists in the archive and could not be reconstructed**, so the cause is recorded as unidentified rather than guessed at. The most likely candidate is a mid-update unit_status in rescreen_corrected.csv, since Task A donor resolution was being iterated in the same minutes.

**Fix.** src/15n_inflation_census.R pins SEED <- 20260805L, draws every bootstrap resample from it, and records the seed in inflation_census.csv. tests/test_determinism.R runs the census estimator twice in one session and requires bit-identical output.

**Numerically:** the census-frame IPW estimate is 2.673×, stable across runs.

#### D-055 — Environment record disagreed with itself

**Date:** 2026-08-05 · **Stage:** 1.5 · **Source:** own defect · **Status:** RESOLVED

docs/AI_USE_STATEMENT.md said the work ran on “macOS 15 (Darwin 25.6.0)”. results/session_info.txt, the Gate 4 environment record written by sessioninfo::session_info(), says **macOS Tahoe 26.6**, **aarch64, darwin23**. Two different OS identifications for one run.

results/session_info.txt is authoritative: it is machine-generated at Gate 4 and archived. The AI use statement is corrected to match, and now also carries the toolchain values the manuscript must use: **R 4.6.1 (2026-06-24)**, **Bioconductor 3.23**, **aarch64-apple-darwin23**, GEOquery 2.80.0, rentrez 1.2.4, data.table 1.18.4.

**Numerically.** No result depends on this. It is a provenance defect, and a reviewer who spots two OS strings for one analysis will discount the rest of the provenance record.

#### D-056 — The excluded population is BETTER covered, not worse

**Date:** 2026-08-05 · **Stage:** 1.5p · **Source:** own analysis · **Status:** ACCEPTED as a reported limitation

The screen removed **13,189 of 16,820** retrieved series, 9,967 of them on the ≥25-sample threshold alone (D-043). Every rate in the paper is conditional on that, and “we don’t know how the excluded population behaves” is a weak answer when it can be measured.

**Method.** 1,262 series drawn from the excluded population, stratified by exclusion reason, seeded from config/screening.yaml (seed 20260805), with every stratum guaranteed at least 50 so the small ones are estimable. L3/L4/L5 measured with the same machinery as the audited population, including run-level coverage and the three-valued statuses.

**Result — and it is the opposite of what was expected.**

| population | n | L3 | L4 | **L5** | median samples | p vs audited |
| --- | --- | --- | --- | --- | --- | --- |
| **AUDITED** (≥25, bulk, Illumina) | 3,631 | 94.13% | 46.74% | **27.29%** | 49 | — |
| **EXCLUDED** (all reasons, sampled) | 1,262 | 96.04% | 48.49% | **35.74%** | 9 | 1.8 × 10⁻⁸ |
| excluded: too_small | 907 | 97.79% | 54.69% | **43.00%** | 8 | 5.4 × 10⁻²⁰ |
| excluded: non_illumina | 50 | 98.00% | 50.00% | 36.00% | 11 | 0.23 |
| excluded: single_cell | 255 | 90.59% | 34.51% | 16.86% | 10 | 3.6 × 10⁻⁴ |
| excluded: array | 50 | 90.00% | 6.00% | **0.00%** | 42 | 3.2 × 10⁻⁵ |

The remediation plan predicted the excluded population would be **less** covered — “compendia ingest what SRA indexes and small studies are less likely to be linked” — which would have made the audited population the optimistic case and **strengthened** the paper’s conclusion. That is not what happened. Small studies are covered *better*, by a wide and highly significant margin.

**Mechanism, stated so the finding is neither over-read nor explained away.** L5 requires **≥90% of a series’ runs** to be present in one resource, and that threshold is far easier to clear with eight runs than with forty-nine: one missing run costs 12.5% of an 8-run series and 2% of a 49-run series. The excluded sample’s median size is 9 against the audited population’s 49, so the comparison is **confounded with size by construction**. The array stratum behaves exactly as it must — 0% L5, because array studies have no reads to reprocess — which is a useful internal check that the measurement is doing what it claims.

**What the manuscript must say.** The audited population is the **pessimistic** case for L5 among sequencing-based series, not the optimistic one. The headline rate does **not** generalise upward to the series the screen removed, and the honest statement is that every reported rate is conditional on the ≥25-sample threshold in a way that **does not resolve in the paper’s favour**.

Reported at equal prominence with the convenient results, per CLAUDE.md’s working-style rule: selective reporting is the failure mode this paper exists to criticise, and a sensitivity analysis is worth nothing if only run in the hope of confirmation.

#### D-057 — The pre-registration claim rested on rewritable git dates

**Date:** 2026-08-05 (Bitcoin anchoring completed 2026-08-06) · **Stage:** 1.6b · **Source:** own defect · **Status:** RESOLVED for OpenTimestamps; Software Heritage remains unavailable

The paper’s methodological selling point is that three analysis plans were frozen before the work they govern. That rested entirely on **git commit dates, which are self-asserted and rewritable** — commit --amend, rebase and filter-branch all rewrite them, and a tag can be moved. A commit hash proves **content integrity**; it does not prove **time**.

**Done.** OpenTimestamps proofs for all four freeze commits (0b10885, 6500647, 03916e7, 366bdd4), in freeze_proofs/, registered with four independent calendar servers. Only a hash leaves the machine; no repository content is published.

**UPDATE 2026-08-06 — the anchoring is complete.** ots upgrade was run and all four proofs now carry **Bitcoin block header attestations**: blocks **961233** (mined 2026-08-06T01:48:11Z) and **961237** (2026-08-06T02:29:29Z). Both block heights and their mined times were confirmed independently against a public block explorer rather than taken from the client’s own report. They may now be described as anchored in Bitcoin.

**Two qualifications that remain, and both belong in the manuscript:**

1. The anchor is an **upper bound**. Stamping happened on 2026-08-05, a day or two after the freeze commits, so it establishes that the manifest content — and therefore the commit, tree and config-blob hashes it names — existed by then. It cannot confirm the minute-level committer dates git asserts. That is still a neutral, non-overwritable timestamp from a source with no trusted authority, which is precisely what git cannot supply.
2. Local ots verify needs a Bitcoin node to look up the block hash and reports a connection error without one. That is a property of the verifying machine, not of the proof; ots info shows the attestations with no node, and anyone can verify at <https://opentimestamps.org>. Do not describe the CLI error as a failed verification.

**A trap worth recording.** The .txt manifests must not be edited. Each proof commits to its file’s SHA-256, and a single byte orphans it — which is exactly what happened when 16b was re-run inside run_all.R and rewrote a stamped_on: line. The manifests are byte-stable now and 16b refuses to touch one that already has a proof.

**Not done: Software Heritage.** Save Code Now ingests from a public URL. This repository has no remote and CLAUDE.md rule 16 forbids pushing, so **no SWHID exists and none has been invented**. The exact command is emitted for the author to run once the repository is public.

**For the manuscript.** No standards body — the Turing Way, COS/OSF, BITSS, rOpenSci — endorses a git commit *as* a formal pre-registration; they treat version control as provenance infrastructure that complements a registry, whose core value is a neutral, non-overwritable timestamp. External anchoring is what earns registry-equivalent credibility. Precedent: TOPO (Casas & Fidler, arXiv:2411.00072, *Open Journal of Astrophysics*). Report all three identifiers — commit hash for content, SWHID for history and a neutral date, Zenodo DOI for a citable snapshot — and say what each does not prove.

#### D-058 — A connection leak, not a GEO block, at exactly 125 records

**Date:** 2026-08-05 · **Stage:** 1.5i2 · **Source:** own defect · **Status:** RESOLVED

Route B died after exactly 125 records with all 128 connections are in use.

**Cause.** geo_record_html() read with

tryCatch(readLines(curl::curl(url), warn = FALSE), error = function(e) NULL)

The inner tryCatch handles errors, but the caller geo_request() also catches **warnings** and treats them as failures. A warning raised during the read unwinds the stack one frame up, past the inner handler, leaving the curl connection created by curl::curl() open and unregistered by anything that would close it. One leaked connection per such fetch; R’s connection table holds 128.

**Fix.** The connection is created explicitly and closed on every exit path via finally = try(close(con), silent = TRUE). Verified: three live fetches now leak zero connections. Route B then completed all 237 records in 2.6 minutes with zero blocks.

**This may partly explain D-029.** That entry diagnosed an IP-level GEO block because “the first census run completed exactly 125 series, then every one of the next 193 failed” — consecutive successes to #125, consecutive failures after. A 128-connection ceiling produces exactly that signature, and 125 is the same boundary observed here. The two runs are not identical (D-029 counted 125 *series* at ~3 requests each; this was 125 *requests*), so the earlier diagnosis is not simply overturned, and the conservative pacing it introduced has done no harm. But **the evidence that GEO imposes a cumulative per-window quota is weaker than D-029 states**, and a reader should be told that the block boundary and the connection-table limit coincide. Recorded rather than quietly corrected.

#### D-059 — Grouping variables were read as donor identifiers

**Date:** 2026-08-05 · **Stage:** 1.5m / 1.5n · **Source:** own defect · **Status:** RESOLVED

Found while putting the inflation statistics on one frame (D-049), by looking at which series sat at the top of the distribution.

count_biological_units() selected donor fields with grepl(PATIENT_FIELD_RX, keys, ignore.case = TRUE) — a **substring** match, guarded only by “more than one distinct value”. It therefore accepted patient group, subject diagnosis and donor group, which are **grouping variables**, and read a two-level case/control field as “two donors”.

| series | samples | field | distinct values | implied inflation |
| --- | --- | --- | --- | --- |
| GSE145669 | 9,148 | individual | 2 | **4,574×** |
| GSE304327 | 666 | donor | 4 | 166× |
| GSE183635 | 2,351 | patient group | 28 | 84× |

The damage runs in **both** directions. Upward, one parsing artefact set the maximum used by the upper Manski bound. Downward, collapsing hundreds of samples into a handful of “donors” makes a series fail L1c for a reason that has nothing to do with biological replication.

**Fix.** Two guards, neither of which invents a new outcome — a field that fails them yields ambiguous, which already means “repeated sampling possible but unestablished, manual review, never a pass”:

1. **Name.** A key matching an identifier word *and* a group-label word is a group label. No threshold involved.
2. **Plausibility.** A genuine subject identifier does not average more than donor_max_samples_per_unit samples per subject. Declared **post-hoc** in config/screening.yaml at 20, deliberately permissive, and it only ever moves a series out of both the numerator and the denominator of a donor-conditional statistic.

The guards are applied to the ARCHS4-derived resolutions directly and to the archived census checkpoints post-hoc, since those were written before the guards existed and record both the field name and its values.

**Numerically:**

|  | before guard | after guard |
| --- | --- | --- |
| series usable for donor counting | 293 | **247** |
| accessions per donor, that stratum | 4.26 | **2.65** |
| max per-series inflation | 4,574× | **20×** (the cap) |
| L1c weighted | 3.535% | **4.053%** |
| L1c stratum size | 229 | **199** |

**Consequence to report.** The upper Manski bound is now partly a function of a declared post-hoc parameter rather than of the data alone, because it is driven by the largest observed per-series value and that value is capped. inflation_census.csv says so in its basis column, and src/15n_inflation_census.R prints it.

#### D-060 — run_all.R’s “offline” mode was a guess, and it started a GEO crawl

**Date:** 2026-08-05 · **Stage:** 1.6 · **Source:** own defect · **Status:** RESOLVED

Found by running the remediation’s own acceptance test.

run_all.R documents its default mode as “offline … No external resource is contacted.” That was enforced at the **stage** layer, by a heuristic: if a stage’s checkpoint cache exists and holds at least the expected number of files, assume running it makes no network calls.

**The heuristic is unknowable in general, and false here.** src/15c_census.R loops all 3,631 candidates and fetches any without a checkpoint. Only **582** are banked, because the study deliberately censused a stratum rather than the population (FREEZE-S). The cache was “complete” by the registry’s count and radically incomplete by the script’s needs.

**What happened.** An end-to-end Rscript run_all.R began downloading GEO SOFT records for series that had never been fetched. Caught after about an hour, at **2,049 of 3,631** census checkpoints (from 582) and 2,359 cached SOFT files (from 1,130).

**Two harms, both repaired.**

1. **A frozen design was at risk.** Had the crawl completed and the cascade then been re-run, L2 would have silently expanded from 1,084 series to ~3,000 and the S1/S2 weighting would have been computed over a population the sampling plan never described. The 1,467 crawled checkpoints were quarantined to data/cache/census_unplanned_crawl/, restoring the cache to the 582 the frozen design assumes. Nothing was deleted.
2. **A committed result table was overwritten.** Re-running src/15b2_screen_revised.R rewrote results/tables/audit_l0_rev2_by_disease.csv with **1,263,376 rows in place of 1,124** — an assemble-step defect in that script, distinct from the crawl, and one the original run also hit (STATUS.md 1.5b2 logs “1263376 descriptors”). Restored from git. Every other committed table was checked; none had changed row count.

**Fix, at the layer where it is a fact rather than a guess.** IPF_OFFLINE=1, set by run_all.R in its default mode, makes every outbound call **stop loudly**: ncbi_call(), geo_request(), geo_record_html(), metasra_study() and recount3_project_runs(). It does not return NULL — callers treat NULL as a transient failure and would write a partial result, which is precisely the failure mode this project exists to criticise. require_entrez_key() becomes a no-op under the same flag, since offline no Entrez call can occur and the gate would otherwise block a stage that only reads its cache.

A partial stage category now marks the five stages whose caches are **intentionally** incomplete (15b2, 15c, 15c2, 15j, 15r). Offline they are skipped **by name with a stated reason** rather than attempted. 15r is there for a different reason worth recording: 31 of its 3,270 MetaSRA lookups failed, and a failed lookup is deliberately not cached, so its cache can never cover the population.

**Acceptance, after the fix:** Rscript run_all.R completes in **2.3 minutes** — 25 stages ok, 5 skipped by name, **0 failed**, no external resource contacted. Verified both directions: the guard blocks a live call when set and permits it when unset.

**Numerically.** No reported quantity changed as a result of the crawl or its repair. l1_layers.csv, the cascade tables and the census are unaffected. The re-run did propagate the D-059 donor guard into s1_census_l1.csv and cascade_l1.csv, changing unit_status for 16 series and n_units for 18 — the intended effect of that fix reaching a table written before it existed, not a consequence of this one.
