## Supplementary files 1-7 for "Most published human disease RNA-seq cannot be uniformly reanalyzed: a population-scale audit of reprocessability in the Gene Expression Omnibus": Additional_file_4_regression_test_suite.txt

ADDITIONAL FILE 4 - REGRESSION TEST SUITE
Reprocessability audit. R test files, reproduced in full.
==============================================================================

CONTENTS
   1. test_coverage_bounds.R
   2. test_determinism.R
   3. test_deviations.R
   4. test_eligibility.R
   5. test_freeze_integrity.R
   6. test_l1_layers.R
   7. test_mesh.R
   8. test_population.R
   9. test_retry.R
  10. test_three_valued.R

==============================================================================


==============================================================================
FILE: tests/test_coverage_bounds.R
==============================================================================

## tests/test_coverage_bounds.R -- a proportion must be a proportion.
##
## D-044. `recount3_cov` divided the matched recount3 PROJECT's sample count by
## the GEO SERIES' sample count. 79 series exceeded 1, to a maximum of 61.44.
## This single assertion would have caught it the day it was written.
##
## Run: Rscript tests/test_coverage_bounds.R

suppressPackageStartupMessages({ library(testthat) })
source("src/utils/runlevel.R")

CENSUS <- "results/tables/l3_l5_census.csv"
RL     <- "results/tables/recount3_runlevel.csv"
EXC    <- "results/tables/excluded_population_series.csv"

test_that("every coverage column in the census is NA or within [0, 1]", {
  skip_if_not(file.exists(CENSUS), "census absent")
  d <- read.csv(CENSUS, stringsAsFactors = FALSE)
  cols <- grep("_cov$", names(d), value = TRUE)
  ## The archived project-level ratio is retained deliberately, as evidence of
  ## the defect, and is the one column exempt from the bound.
  cols <- setdiff(cols, "recount3_cov_projectlevel")
  expect_gt(length(cols), 0L)
  for (k in cols) {
    v <- d[[k]]
    expect_true(all(is.na(v) | (v >= 0 & v <= 1)),
                info = sprintf("%s: %d value(s) outside [0,1], max %.3f",
                               k, sum(!is.na(v) & (v < 0 | v > 1)), max(v, na.rm = TRUE)))
  }
})

test_that("the retained project-level column still shows the defect it documents", {
  skip_if_not(file.exists(CENSUS), "census absent")
  d <- read.csv(CENSUS, stringsAsFactors = FALSE)
  skip_if_not("recount3_cov_projectlevel" %in% names(d), "pre-fix census")
  ## If this ever stops exceeding 1, the evidence for D-044 has been lost.
  expect_gt(max(d$recount3_cov_projectlevel, na.rm = TRUE), 1)
})

test_that("run-level coverage is bounded and consistent with its own counts", {
  skip_if_not(file.exists(RL), "run-level table absent")
  r <- read.csv(RL, stringsAsFactors = FALSE)
  for (k in c("recount3_cov_runlevel", "dee2_cov_runlevel")) {
    v <- r[[k]]
    expect_true(all(is.na(v) | (v >= 0 & v <= 1)), info = k)
  }
  ## An intersection cannot exceed either side.
  ok <- !is.na(r$n_overlap_recount3) & r$series_run_status == "resolved"
  expect_true(all(r$n_overlap_recount3[ok] <= r$n_series_runs[ok]))
  expect_true(all(r$n_overlap_recount3[ok] <= r$n_recount3_runs[ok]))
})

test_that("coverage is NA exactly when it is unassessable, and vice versa", {
  skip_if_not(file.exists(CENSUS), "census absent")
  d <- read.csv(CENSUS, stringsAsFactors = FALSE)
  skip_if_not("recount3_status" %in% names(d), "pre-fix census")
  for (nm in c("recount3", "dee2", "archs4")) {
    cov <- d[[paste0(nm, "_cov")]]; st <- d[[paste0(nm, "_status")]]
    expect_identical(is.na(cov), st == "unassessable", info = nm)
  }
})

test_that("status_of() is three-valued and never collapses NA to a negative", {
  expect_identical(status_of(NA_real_, 0.9), "unassessable")
  expect_identical(status_of(0.95, 0.9), "covered")
  expect_identical(status_of(0.90, 0.9), "covered")     # boundary is inclusive
  expect_identical(status_of(0.5,  0.9), "not_covered")
  expect_identical(status_of(0,    0.9), "not_covered")
})

test_that("runlevel_cov() refuses to guess and is bounded by construction", {
  a <- c("SRR1", "SRR2", "SRR3", "SRR4")
  expect_equal(runlevel_cov(a, c("SRR1", "SRR2"), "resolved", "resolved"), 0.5)
  expect_equal(runlevel_cov(a, a, "resolved", "resolved"), 1)
  ## extra runs on the resource side cannot push it above 1 -- the defect
  expect_equal(runlevel_cov(a, c(a, paste0("SRR", 5:500)), "resolved", "resolved"), 1)
  expect_true(is.na(runlevel_cov(a, a, "unresolved", "resolved")))
  expect_true(is.na(runlevel_cov(a, a, "resolved", "unresolved")))
  expect_true(is.na(runlevel_cov(character(0), a, "resolved", "resolved")))
})

test_that("the excluded-population sample obeys the same bound", {
  skip_if_not(file.exists(EXC), "excluded-population table absent")
  d <- read.csv(EXC, stringsAsFactors = FALSE)
  for (k in grep("_cov$", names(d), value = TRUE)) {
    v <- d[[k]]
    expect_true(all(is.na(v) | (v >= 0 & v <= 1)), info = k)
  }
})

cat("\ntests/test_coverage_bounds.R: all assertions passed\n")


==============================================================================
FILE: tests/test_determinism.R
==============================================================================

## tests/test_determinism.R -- one statistic, one value.
##
## D-054. STATUS.md logs two runs of 01h_inflation_bounds.R fifty-four seconds
## apart reporting IPW 5.09x and then 4.60x. The conditional estimate and both
## bounds were identical in each, so only the weighted estimator moved.
##
## This test does not reconstruct that state -- it no longer exists in the
## archive. It pins the property that was missing: run the estimator twice in
## one session and require the same answer, and require the seed to be recorded
## alongside the result so a future reader can tell whether a difference is a
## real change or a re-draw.
##
## Run: Rscript tests/test_determinism.R

suppressPackageStartupMessages({ library(testthat) })

INF <- "results/tables/inflation_census.csv"
DON <- "results/tables/donor_resolution_by_series.csv"

test_that("the census inflation table records its seed", {
  skip_if_not(file.exists(INF), "inflation table absent")
  d <- read.csv(INF, stringsAsFactors = FALSE)
  expect_true("seed" %in% names(d))
  expect_true(all(!is.na(d$seed)))
  expect_equal(length(unique(d$seed)), 1L)
})

test_that("the bootstrap for the Imbens-Manski CI is reproducible", {
  skip_if_not(file.exists(INF) && file.exists(DON), "inputs absent")
  d <- read.csv(INF, stringsAsFactors = FALSE)
  seed <- d$seed[1]
  don <- read.csv(DON, stringsAsFactors = FALSE)
  don$resolvable <- don$unit_status %in% c("resolved", "resolved_inferred") &
                    don$n_distinct_donors > 1L

  ## The same bound estimator 15n uses, run twice from the recorded seed.
  bound_pair <- function(x) {
    r <- x$resolvable
    ts <- sum(x$n_samples[r]); td <- sum(x$n_distinct_donors[r])
    us <- sum(x$n_samples[!r]); all_s <- ts + us
    mi <- max(x$n_samples[r] / x$n_distinct_donors[r])
    c(all_s / (td + us), all_s / (td + us / mi))
  }
  draw <- function() {
    set.seed(seed)
    replicate(50, bound_pair(don[sample.int(nrow(don), nrow(don), replace = TRUE), ]))
  }
  expect_identical(draw(), draw())
})

test_that("the point estimates are pure functions of the tables, not of run order", {
  skip_if_not(file.exists(DON), "donor summary absent")
  don <- read.csv(DON, stringsAsFactors = FALSE)
  don$resolvable <- don$unit_status %in% c("resolved", "resolved_inferred") &
                    don$n_distinct_donors > 1L
  cond <- function() {
    r <- don$resolvable
    sum(don$n_samples[r]) / sum(don$n_distinct_donors[r])
  }
  expect_identical(cond(), cond())
  skip_if_not(file.exists(INF), "inflation table absent")
  d <- read.csv(INF, stringsAsFactors = FALSE)
  expect_equal(d$value[d$quantity == "conditional"], cond(), tolerance = 1e-9)
})

test_that("the archived 51-series estimator is deterministic in-session", {
  f <- "results/tables/rescreen_corrected.csv"
  g <- "results/tables/design_features.csv"
  skip_if_not(file.exists(f) && file.exists(g), "roster inputs absent")
  x <- merge(read.csv(f, stringsAsFactors = FALSE),
             read.csv(g, stringsAsFactors = FALSE)[, -2], by = "accession")
  x$log_gsm <- log(x$n_gsm)
  res <- x$unit_status == "resolved"
  ipw <- function() {
    fit <- suppressWarnings(glm(res ~ log_gsm + n_char_keys + repeat_design_marker +
                                  title_repeat_frac, data = x, family = binomial()))
    p <- pmin(pmax(fitted(fit), 0.02), 0.98); w <- 1 / p[res]
    sum(w * x$n_gsm[res]) / sum(w * x$n_biological_units[res])
  }
  a <- ipw(); b <- ipw()
  expect_identical(a, b)
  ## The value that reproduces today. If this ever moves, an INPUT changed --
  ## which is the diagnosis D-054 could not make retrospectively.
  expect_equal(a, 4.599, tolerance = 0.01)
})

cat("\ntests/test_determinism.R: all assertions passed\n")


==============================================================================
FILE: tests/test_deviations.R
==============================================================================

## tests/test_deviations.R -- the deviation log counts itself.
##
## D-042 family. MANIFEST.md said 40 entries, docs/CP0b_REPORT.md §8 said 41,
## the manuscript said 41, and the truth is 40 with D-026 never issued. Three
## documents disagreed because all three quoted a remembered number. This test
## parses DEVIATIONS.md and writes the counts to a table, so every downstream
## document cites a generated figure.
##
## Run: Rscript tests/test_deviations.R

suppressPackageStartupMessages({ library(testthat) })

lines <- readLines("DEVIATIONS.md", warn = FALSE)

hdr <- grep("^## D-", lines, value = TRUE)
ids <- sub("^## (D-[0-9]+).*$", "\\1", hdr)
nums <- as.integer(sub("^D-", "", ids))

## "Source: ... own defect" appears on the metadata line of each entry.
own <- grep("\\*\\*Source:\\*\\*[^|]*own defect|Source:.*own defect", lines, value = TRUE)

n_entries <- length(ids)
n_own     <- length(own)
max_id    <- max(nums)
gaps      <- setdiff(seq_len(max_id), nums)

test_that("every entry heading has a unique, well-formed identifier", {
  expect_true(all(grepl("^D-[0-9]{3}$", ids)))
  expect_equal(anyDuplicated(ids), 0L)
})

test_that("identifiers are strictly increasing", {
  expect_true(all(diff(nums) > 0))
})

## D-026 was never issued and is a permanent hole. The 2026-08-05 remediation
## reserves D-042..D-058 by subject up front (REMEDIATION_PLAN Appendix B), so
## that the identifier cited for a defect is stable from the moment the defect
## is described, not assigned by whatever order the fixes happened to land in.
## Those identifiers are therefore allowed to be absent WHILE the remediation is
## in flight, and are required to be present once it closes -- which is what
## `remediation_complete` records and what Phase 6 asserts.
NEVER_ISSUED <- 26L
RESERVED     <- 42:60

test_that("no identifier is skipped except D-026 and the reserved remediation range", {
  stray <- setdiff(gaps, c(NEVER_ISSUED, RESERVED))
  expect_identical(stray, integer(0),
                   info = sprintf("unexplained gap(s): %s",
                                  paste0("D-", sprintf("%03d", stray), collapse = ", ")))
})

test_that("D-026 is absent and is documented as never issued", {
  expect_true(NEVER_ISSUED %in% gaps)
})

remediation_written  <- setdiff(RESERVED, gaps)
remediation_pending  <- intersect(RESERVED, gaps)
remediation_complete <- length(remediation_pending) == 0L

test_that("entry count is the number of headings, not the highest identifier", {
  expect_lt(n_entries, max_id)
  expect_equal(n_entries, max_id - length(gaps))
})

## MANIFEST.md and CP0b_REPORT.md describe the ORIGINAL run, so they quote the
## original-run count (40), not the running total, which grows as remediation
## entries land. Distinguishing the two is the whole point: the documents were
## wrong because all three quoted a remembered number.
n_original     <- sum(nums < min(RESERVED))
n_own_original <- length(grep("Source:.*own defect", lines[seq_len(
  if (any(nums >= min(RESERVED)))
    grep(sprintf("^## D-%03d", min(nums[nums >= min(RESERVED)])), lines)[1] - 1L
  else length(lines))], value = TRUE))

test_that("the numbering note is present and states the original-run counts", {
  head_txt <- paste(head(lines, 40), collapse = "\n")
  expect_match(head_txt, "D-026` was never issued")
  expect_match(head_txt, sprintf("\\*\\*%d entries\\*\\*", n_original))
  expect_match(head_txt, "D-042")
})

## Documents that quote these counts must agree with them.
check_doc <- function(path, pattern, label) {
  test_that(sprintf("%s states %s", basename(path), label), {
    skip_if_not(file.exists(path), sprintf("%s absent", path))
    txt <- paste(readLines(path, warn = FALSE), collapse = "\n")
    expect_match(txt, pattern)
  })
}
check_doc("MANIFEST.md", sprintf("%d entries", n_original),
          sprintf("%d entries", n_original))
check_doc("MANIFEST.md", sprintf("five defects"), "five own-code defects")
check_doc("docs/CP0b_REPORT.md", sprintf("%d deviations", n_original),
          sprintf("%d deviations", n_original))
check_doc("docs/CP0b_REPORT.md", "five defects", "five own-code defects")

fmt <- function(x) if (length(x)) paste0("D-", sprintf("%03d", x), collapse = ";") else "none"

out <- data.frame(
  quantity = c("n_entries", "n_own_defect", "max_identifier",
               "skipped_identifiers", "n_skipped",
               "n_entries_original_run", "n_entries_remediation",
               "remediation_pending", "remediation_complete"),
  value = c(as.character(n_entries), as.character(n_own), as.character(max_id),
            fmt(gaps), as.character(length(gaps)),
            as.character(n_original),
            as.character(length(remediation_written)),
            fmt(remediation_pending), as.character(remediation_complete)),
  basis = "parsed from DEVIATIONS.md by tests/test_deviations.R",
  stringsAsFactors = FALSE)

dir.create("results/tables", recursive = TRUE, showWarnings = FALSE)
write.csv(out, "results/tables/deviation_counts.csv", row.names = FALSE)

cat("\n-- deviation counts --\n")
print(out, row.names = FALSE)
cat("\nWrote results/tables/deviation_counts.csv\n")


==============================================================================
FILE: tests/test_eligibility.R
==============================================================================

## tests/test_eligibility.R -- known-answer tests for the CP-0b screening fix.
##
## The B1 filter exists because an exact-term list failed once already
## (GSE188464, "cultered"). These cases pin the behaviour so it cannot silently
## regress. Run: Rscript tests/test_eligibility.R

source("src/utils/eligibility.R")
suppressPackageStartupMessages(library(testthat))

# ---- B1: must EXCLUDE ------------------------------------------------------

test_that("GSE188464's verbatim record is excluded despite GEO's typo", {
  ## The literal string from the live record -- "cultered", not "cultured".
  fields <- paste(
    "Disease-specific transcriptomic changes induced by NFkB inhibition in",
    "idiopathic pulmonary fibrosis.",
    "Primary cultered human fibroblasts from 16 individuals, 8 with idiopathic",
    "pulmonary fibrosis (IPF) and 8 age-matched controls.",
    "Primary lung fibroblasts derived from eight patients with IPF and eight",
    "age-matched non-diseased controls (NDC) were treated with 0, 3 or 10 µM",
    "ACT001 in the presence/absence of pro-fibrotic cytokine TGF-b1 for 24 hours.")
  r <- screen_eligibility("GSE188464",
        sample_fields = "Primary Lung Fibroblasts, Apex drug treatment: Control drug concentration: 0uM",
        study_fields = fields,
        char_keys = c("participant id","age","gender","disease state",
                      "drug treatment","drug concentration"))
  expect_false(r$eligible)
  expect_match(r$ineligible_reason, "in vitro")
  expect_match(r$ineligible_reason, "perturbation")
})

# ---- B1: real records that MUST be admitted (regression guards) ------------
## These three were wrongly excluded by the first version of the filter, which
## searched the whole summary. Sample-level fields are taken verbatim from the
## cached GEO records.

test_that("GSE92592 is admitted -- 'in vitro' appears only in its summary", {
  r <- screen_eligibility("GSE92592",
        sample_fields = "lung tissue tissue: lung disease state: Idiopathic Pulmonary Fibrosis",
        study_fields  = "... we confirmed these findings in vitro ...",
        char_keys = c("tissue","disease state"))
  expect_true(r$eligible)
  expect_true(r$flag_for_review)     # recorded for audit, but not excluded
})

test_that("GSE134692 is admitted -- ALI here is ACUTE LUNG INJURY", {
  r <- screen_eligibility("GSE134692",
        sample_fields = paste("human subject tissue: Lung batch: 2 diseasestatus: ALI",
                              "samplename: ALI_R2_B2 age: 45 tobacco: Never smoked"),
        study_fields  = "Transplant stage IPF lung",
        char_keys = c("tissue","batch","diseasestatus","age","race","gender","tobacco"))
  expect_true(r$eligible)
})

test_that("GSE334185 is still excluded -- on sample-level evidence", {
  r <- screen_eligibility("GSE334185",
        sample_fields = paste("lung tissue: lung cell line: primary cell type: fibroblast",
                              "infection: IAV treatment: QUI disease: healthy"),
        study_fields  = "siRNA-mediated depletion of IRF3 or STAT1",
        char_keys = c("tissue","cell line","cell type","age","Sex","infection",
                      "treatment","disease","batch"))
  expect_false(r$eligible)
  expect_match(r$ineligible_reason, "in vitro")
})

test_that("GSE213001 is admitted -- rnaconcentration is a QC metric, not a dose", {
  r <- screen_eligibility("GSE213001",
        sample_fields = paste("Lung,IPF.Apex,Whole lung tissue: Whole lung donorid: ALF008",
                              "rnaconcentration ngul: 201 lungweight mg: 22 rin: 7.7"),
        study_fields  = "Multi-region lung tissue RNA-seq in IPF",
        char_keys = c("tissue","donorid","diseasegroup","rnaconcentration ngul",
                      "lungweight mg","rin","volum ul","processingdate"))
  expect_true(r$eligible)
})

test_that("a genuine drug-dose design is still caught", {
  expect_false(screen_eligibility("m", sample_fields = "lung",
    char_keys = c("tissue","drug concentration","drug treatment"))$eligible)
})

test_that("a literal 'cultured' term would NOT have caught it -- stem is required", {
  fields <- "Primary cultered human fibroblasts from 16 individuals"
  expect_false(grepl("cultured", fields, fixed = TRUE))   # the original miss
  expect_false(screen_eligibility("x", sample_fields = fields)$eligible)  # the stem catches it
})

test_that("cell-culture and perturbation designs are excluded", {
  expect_false(screen_eligibility("a", sample_fields = "primary lung fibroblast cell line")$eligible)
  expect_false(screen_eligibility("b", sample_fields = "airway epithelial cells at air-liquid interface")$eligible)
  expect_false(screen_eligibility("b2", sample_fields = "lung", char_keys = c("cell type","dose"))$eligible)
  expect_false(screen_eligibility("c", sample_fields = "fibroblasts treated with TGFb1 vs control")$eligible)
  expect_false(screen_eligibility("d", sample_fields = "siRNA knockdown of SMAD3 in lung fibroblasts")$eligible)
  expect_false(screen_eligibility("e", sample_fields = "organoid cultures derived from IPF lung")$eligible)
  expect_false(screen_eligibility("f", sample_fields = "cells exposed to 10 nM dexamethasone")$eligible)
})

test_that("model-organism series are excluded", {
  expect_false(screen_eligibility("g", sample_fields = "C57BL/6 mice, bleomycin-induced fibrosis")$eligible)
  expect_false(screen_eligibility("h", sample_fields = "genotype: SftpcI73T/I73T tamoxifen")$eligible)
})

# ---- B1: must ADMIT (guard against over-exclusion) -------------------------
## Over-exclusion is as damaging as under-exclusion: it would shrink Paper 1's
## L1 denominator and make the cascade look worse than it is.

test_that("genuine bulk lung tissue case-control series are admitted", {
  expect_true(screen_eligibility("i",
    "Whole lung homogenate from 20 IPF patients and 19 controls, Lung Tissue Research Consortium")$eligible)
  expect_true(screen_eligibility("j",
    "Bulk RNA-seq of lung tissue from patients with idiopathic pulmonary fibrosis and non-fibrotic donor lung")$eligible)
  expect_true(screen_eligibility("k",
    "paired central and peripheral lung explant biopsies from 13 IPF and 8 non-IPF donors")$eligible)
})

test_that("the word 'control' alone does not trigger the perturbation filter", {
  expect_true(screen_eligibility("l", sample_fields = "IPF cases versus healthy control lung tissue")$eligible)
})

# ---- B2: biological units, not GSM files -----------------------------------

test_that("distinct donors are counted, not sample files", {
  ## 6 files, 2 donors, 3 dose arms each -- the GSE188464 shape.
  ch <- lapply(1:6, function(i)
    c(`participant id` = paste0("P", ((i - 1) %/% 3) + 1),
      `disease state`  = if (i <= 3) "IPF" else "Normal"))
  r <- count_biological_units(ch)
  expect_equal(r$status, "resolved")
  expect_equal(r$n_units, 2)          # NOT 6
})

test_that("paired-region sampling collapses to patients", {
  ch <- lapply(1:4, function(i) c(subject = paste0("P", ((i - 1) %/% 2) + 1)))
  r <- count_biological_units(ch)
  expect_equal(r$n_units, 2)          # NOT 4
})

test_that("a constant identifier column is not treated as an identifier", {
  ch <- lapply(1:4, function(i) c(donor = "pooled", tissue = "lung"))
  expect_true(count_biological_units(ch)$status != "resolved")
})

test_that("no donor field with repeated title stems is ambiguous, never a pass", {
  r <- count_biological_units(lapply(1:4, function(i) c(tissue = "lung")),
                              titles = c("Sma1C", "Sma1P", "Sma2C", "Sma2P"))
  expect_equal(r$status, "ambiguous")
  expect_true(r$status != "resolved")
})

test_that("one sample per patient with no donor field is unresolved, not ambiguous", {
  r <- count_biological_units(lapply(1:3, function(i) c(tissue = "lung")),
                              titles = c("IPF_alpha", "IPF_beta", "CTRL_gamma"))
  expect_equal(r$status, "unresolved")
  expect_equal(r$n_units, 3)
})

cat("\nAll eligibility tests passed.\n")


==============================================================================
FILE: tests/test_freeze_integrity.R
==============================================================================

## tests/test_freeze_integrity.R -- prove the frozen configs did not move.
##
## D-053. The manuscript said the pre-registered configs were "literally
## unmodifiable" because a Claude Code deny rule forbade editing them. That
## mechanism is weaker than described: a permission rule only binds a session
## whose project root is this repository, it is a property of the working copy
## rather than of the archive, and a reader of the deposit cannot check it at
## all. Git can be checked by anyone with the deposit, so that is what this
## test uses.
##
## Two assertions per frozen config:
##   1. the freeze commit exists and the recorded hash is real
##   2. the PARSED content at HEAD is identical to the parsed content at the
##      freeze commit -- no parameter moved
##
## Assertion 2 rather than "no commit touched the file" is deliberate, and the
## difference is a real finding: config.yaml WAS committed once after its
## freeze (4d3d31c), which wrote the freeze hash into its own header comment.
## No parameter changed. A test that forbade any post-freeze commit would fail
## on a purely bookkeeping edit and would tell a reader less, not more.
##
## Run: Rscript tests/test_freeze_integrity.R

suppressPackageStartupMessages({ library(testthat); library(yaml) })

FREEZES <- list(
  list(freeze = "FREEZE-A",    file = "config/config.yaml",   hash = "0b10885",
       note = "disease-selection rule, frozen before any coverage query"),
  list(freeze = "FREEZE-L0P",  file = "config/l0prime.yaml",  hash = "03916e7",
       note = "attribution-verification plan"),
  list(freeze = "FREEZE-S",    file = "config/sampling.yaml", hash = "366bdd4",
       note = "verification sampling plan, committed before the sample was drawn")
)

git <- function(...) {
  out <- suppressWarnings(system2("git", c(...), stdout = TRUE, stderr = FALSE))
  if (!is.null(attr(out, "status")) && attr(out, "status") != 0) character(0) else out
}

rows <- list()

for (fz in FREEZES) {
  test_that(sprintf("%s (%s) is a real commit that touched the file", fz$freeze, fz$hash), {
    full <- git("rev-parse", fz$hash)
    expect_length(full, 1L)
    expect_true(nchar(full[1]) == 40L)
    touched <- git("show", "--name-only", "--format=", fz$hash)
    expect_true(fz$file %in% touched,
                info = sprintf("%s did not touch %s", fz$hash, fz$file))
  })

  test_that(sprintf("%s content is unchanged since %s", fz$file, fz$freeze), {
    at_freeze <- git("show", sprintf("%s:%s", fz$hash, fz$file))
    expect_gt(length(at_freeze), 0L)
    now <- readLines(fz$file, warn = FALSE)

    y_freeze <- yaml::yaml.load(paste(at_freeze, collapse = "\n"))
    y_now    <- yaml::yaml.load(paste(now, collapse = "\n"))
    expect_identical(y_now, y_freeze,
                     info = sprintf("a PARAMETER changed in %s after %s", fz$file, fz$freeze))
  })

  after <- git("log", "--format=%h", sprintf("%s..HEAD", fz$hash), "--", fz$file)
  at_freeze <- git("show", sprintf("%s:%s", fz$hash, fz$file))
  now <- readLines(fz$file, warn = FALSE)
  rows[[length(rows) + 1L]] <- data.frame(
    freeze = fz$freeze, file = fz$file, freeze_commit = fz$hash,
    n_commits_after_freeze = length(after),
    commits_after_freeze = if (length(after)) paste(after, collapse = ";") else NA_character_,
    parsed_content_identical = identical(
      yaml::yaml.load(paste(now, collapse = "\n")),
      yaml::yaml.load(paste(at_freeze, collapse = "\n"))),
    bytes_identical = identical(now, at_freeze),
    note = fz$note, stringsAsFactors = FALSE)
}

## screening.yaml is deliberately NOT frozen. If it ever acquires a deny rule or
## is described as pre-registered, that is the error this guards against.
test_that("screening.yaml is declared post-hoc and is not frozen", {
  expect_true(file.exists("config/screening.yaml"))
  txt <- paste(readLines("config/screening.yaml", warn = FALSE), collapse = "\n")
  expect_match(txt, "POST-HOC DECLARATION, NOT A PRE-REGISTRATION")
  settings <- paste(readLines(".claude/settings.json", warn = FALSE), collapse = "")
  expect_false(grepl("screening.yaml", settings, fixed = TRUE))
})

tab <- do.call(rbind, rows)
dir.create("results/tables", recursive = TRUE, showWarnings = FALSE)
write.csv(tab, "results/tables/freeze_integrity.csv", row.names = FALSE)

cat("\n-- freeze integrity --\n")
print(tab[, c("freeze", "file", "freeze_commit", "n_commits_after_freeze",
              "parsed_content_identical", "bytes_identical")], row.names = FALSE)
cat("\nWrote results/tables/freeze_integrity.csv\n")


==============================================================================
FILE: tests/test_l1_layers.R
==============================================================================

## tests/test_l1_layers.R -- L1 must know what it counted.
##
## D-046: evaluate_l1() was called without unit_ids, so counted_on was "samples"
## for every pair, and `counted_on` was then discarded so nothing recorded it.
## D-047: the reported L1 rate was design eligibility alone.
##
## Run: Rscript tests/test_l1_layers.R

suppressPackageStartupMessages({ library(testthat) })
source("src/utils/casecontrol.R")

PAIR  <- "results/tables/audit_pair_level.csv"
LAY   <- "results/tables/l1_layers.csv"
DON   <- "results/tables/donor_ids_by_series.csv"
DONS  <- "results/tables/donor_resolution_by_series.csv"

# ---- the function itself ----------------------------------------------------

blobs_case <- rep("IPF lung tissue disease state: idiopathic pulmonary fibrosis", 20)
blobs_ctrl <- rep("control healthy donor lung tissue", 20)
B <- c(blobs_case, blobs_ctrl)
ALTS <- c("idiopathic pulmonary fibrosis", "IPF")

test_that("without unit_ids the outcome is downgraded, not silently passed", {
  ev <- evaluate_l1(B, ALTS)
  expect_identical(ev$counted_on, "samples")
  expect_identical(ev$status, "pass_sample_counted")
  expect_true(ev$l1_pass)
})

test_that("with resolved unit_ids the donor branch is taken", {
  ## 20 cases from 4 donors, 20 controls from 4 donors -> fails on donors
  ids <- c(rep(paste0("d", 1:4), each = 5), rep(paste0("c", 1:4), each = 5))
  ev <- evaluate_l1(B, ALTS, unit_ids = ids, unit_status = "resolved")
  expect_identical(ev$counted_on, "biological_units")
  expect_identical(ev$status, "fail_size")
  expect_equal(ev$n_case, 4L)
  expect_equal(ev$n_control, 4L)
  expect_false(ev$l1_pass)
})

test_that("an unresolved status keeps sample counting even if ids are supplied", {
  ids <- c(rep(paste0("d", 1:4), each = 5), rep(paste0("c", 1:4), each = 5))
  ev <- evaluate_l1(B, ALTS, unit_ids = ids, unit_status = "unresolved")
  expect_identical(ev$counted_on, "samples")
})

test_that("resolved_inferred is honoured, ambiguous is not", {
  ids <- c(rep(paste0("d", 1:4), each = 5), rep(paste0("c", 1:4), each = 5))
  expect_identical(evaluate_l1(B, ALTS, unit_ids = ids,
                               unit_status = "resolved_inferred")$counted_on,
                   "biological_units")
  expect_identical(evaluate_l1(B, ALTS, unit_ids = ids,
                               unit_status = "ambiguous")$counted_on, "samples")
})

test_that("donor counting cannot inflate a count above sample counting", {
  ids <- c(rep(paste0("d", 1:4), each = 5), rep(paste0("c", 1:4), each = 5))
  s <- evaluate_l1(B, ALTS)
  d <- evaluate_l1(B, ALTS, unit_ids = ids, unit_status = "resolved")
  expect_lte(d$n_case, s$n_case)
  expect_lte(d$n_control, s$n_control)
})

# ---- what the pipeline persisted --------------------------------------------

test_that("the pair table records HOW each pair was counted", {
  skip_if_not(file.exists(PAIR), "pair table absent")
  p <- read.csv(PAIR, stringsAsFactors = FALSE)
  for (nm in c("status", "counted_on", "counted_on_donor", "donor_counted",
               "l1_size_pass_donor", "unit_status", "case_rule"))
    expect_true(nm %in% names(p), info = nm)
  expect_true(all(p$counted_on_donor %in% c("samples", "biological_units")))
  ## The donor branch must actually have been exercised somewhere.
  expect_gt(sum(p$counted_on_donor == "biological_units"), 0L)
  ## and the archived behaviour is still recorded for comparison
  expect_true(all(p$counted_on == "samples"))
})

test_that("donor_counted agrees with counted_on_donor", {
  skip_if_not(file.exists(PAIR), "pair table absent")
  p <- read.csv(PAIR, stringsAsFactors = FALSE)
  expect_identical(as.logical(p$donor_counted), p$counted_on_donor == "biological_units")
})

test_that("evaluate_l1 received ids wherever the donor table says it could", {
  skip_if_not(file.exists(PAIR) && file.exists(DONS), "tables absent")
  p <- read.csv(PAIR, stringsAsFactors = FALSE)
  s <- read.csv(DONS, stringsAsFactors = FALSE)
  usable <- s$accession[s$usable_for_donor_counting]
  ## every pair of a usable series must have been donor-counted
  k <- p$accession %in% usable
  if (any(k)) expect_true(all(p$counted_on_donor[k] == "biological_units"))
  ## and no pair of a non-usable series was
  expect_true(all(p$counted_on_donor[!k] == "samples"))
})

test_that("l1_layers.csv carries L1a, L1b and L1c weighted and unweighted", {
  skip_if_not(file.exists(LAY), "layers table absent")
  l <- read.csv(LAY, stringsAsFactors = FALSE)
  for (lay in c("L1a", "L1b", "L1c", "L1b_on_L1c_stratum"))
    for (w in c("weighted", "unweighted"))
      expect_equal(sum(l$layer == lay & l$weighting == w & l$stratum == "pooled"), 1L,
                   info = paste(lay, w))
  expect_true(all(c("S1", "S2", "pooled") %in% l$stratum))
})

test_that("L1b is a subset of L1a, and rates are ordered accordingly", {
  skip_if_not(file.exists(LAY), "layers table absent")
  l <- l0 <- read.csv(LAY, stringsAsFactors = FALSE)
  g <- function(lay, w) l$rate[l$layer == lay & l$weighting == w & l$stratum == "pooled"]
  for (w in c("weighted", "unweighted")) expect_lt(g("L1b", w), g("L1a", w))
  ## L1c is on a different denominator, so it is NOT comparable to L1b directly;
  ## the like-for-like comparator is L1b_on_L1c_stratum, which must exceed it if
  ## accessions overstate replication at all.
  for (w in c("weighted", "unweighted"))
    expect_gt(g("L1b_on_L1c_stratum", w), g("L1c", w))
})

test_that("L1c is reported on the resolvable stratum, not the full frame", {
  skip_if_not(file.exists(LAY), "layers table absent")
  l <- read.csv(LAY, stringsAsFactors = FALSE)
  a <- l[l$layer == "L1a" & l$stratum == "pooled" & l$weighting == "weighted", ]
  c_ <- l[l$layer == "L1c" & l$stratum == "pooled" & l$weighting == "weighted", ]
  expect_lt(c_$n_series, a$n_series)
  expect_match(c_$basis, "RESOLVABLE STRATUM ONLY")
})

test_that("donor ids align to samples and carry a three-valued status", {
  skip_if_not(file.exists(DON), "donor id table absent")
  d <- read.csv(DON, stringsAsFactors = FALSE)
  expect_true(all(c("accession", "sample_index", "gsm", "donor_id",
                    "unit_status", "unit_field") %in% names(d)))
  expect_true(all(d$unit_status %in% c("resolved", "resolved_inferred",
                                       "ambiguous", "unresolved")))
  ## sample_index must be a dense 1..n within each series
  chk <- tapply(d$sample_index, d$accession, function(v) identical(sort(v), seq_along(v)))
  expect_true(all(unlist(chk)))
  ## an unresolved series carries no donor ids
  expect_true(all(is.na(d$donor_id[d$unit_status == "unresolved"])))
})

test_that("the D-059 guard is in force: no absurd samples-per-donor survives", {
  skip_if_not(file.exists(DONS), "donor summary absent")
  s <- read.csv(DONS, stringsAsFactors = FALSE)
  cap <- yaml::read_yaml("config/screening.yaml")$screening$donor_max_samples_per_unit
  u <- s[s$usable_for_donor_counting, ]
  expect_true(all(u$n_samples / u$n_distinct_donors <= cap))
})

cat("\ntests/test_l1_layers.R: all assertions passed\n")


==============================================================================
FILE: tests/test_mesh.R
==============================================================================

## tests/test_mesh.R -- FREEZE-A revision 2 mechanisms.
source("src/utils/mesh.R")
suppressPackageStartupMessages({ library(testthat); library(digest) })

test_that("tree adjudication never counts a sibling", {
  ## The defect that forced the revision: Diabetes Insipidus vs Diabetes Mellitus.
  expect_false(tn_is_self_or_descendant("C19.700.482", "C19.246"))
  expect_false(tn_is_self_or_descendant("C19.246", "C19.700.482"))
  ## Self and true descendants do count.
  expect_true(tn_is_self_or_descendant("C08.381.483", "C08.381.483"))
  expect_true(tn_is_self_or_descendant("C08.381.483.652.500", "C08.381.483"))
})

test_that("tree adjudication is not fooled by string prefixes", {
  expect_false(tn_is_self_or_descendant("C08.381.4830", "C08.381.483"))
  expect_false(tn_is_self_or_descendant("C08.381.483.652.500", "C08.381.484"))
})

test_that("ancestors are dot-truncations", {
  expect_equal(tn_ancestors("C08.381.483.652.500"),
               c("C08", "C08.381", "C08.381.483", "C08.381.483.652"))
  expect_equal(tn_ancestors("C08"), character(0))
})

test_that("comma-inverted MeSH labels are de-inverted", {
  expect_true("Interstitial Lung Diseases" %in% deinvert_label("Lung Diseases, Interstitial"))
  expect_true("Rheumatoid Arthritis" %in% deinvert_label("Arthritis, Rheumatoid"))
  expect_true("Systemic Lupus Erythematosus" %in% deinvert_label("Lupus Erythematosus, Systemic"))
  expect_equal(deinvert_label("Pulmonary Fibrosis"), character(0))   # nothing to invert
})

test_that("the revised query includes de-inverted forms and entry terms", {
  q <- build_query_rev2("Lung Diseases, Interstitial")
  expect_true(grepl('"Interstitial Lung Diseases"[Title]', q, fixed = TRUE))
  expect_true(grepl("[Description]", q, fixed = TRUE))
  q2 <- build_query_rev2("Idiopathic Pulmonary Fibrosis", "Usual Interstitial Pneumonia")
  expect_true(grepl("Usual Interstitial Pneumonia", q2, fixed = TRUE))
})

test_that("a label containing a quote cannot break the query", {
  expect_false(grepl('""', build_query_rev2('Weird "quoted" Disease'), fixed = TRUE))
})

test_that("slugs never collide on non-ASCII input", {
  s <- safe_slug(c("НЕРВНОЙ СИСТЕМЫ", "中枢神経系疾患", "Central Nervous System Diseases"))
  expect_equal(length(unique(s)), 3L)
})

cat("\nAll mesh tests passed.\n")


==============================================================================
FILE: tests/test_population.R
==============================================================================

## tests/test_population.R -- the screening rule reproduces the audited population.
##
## D-042/D-043. The population of 3,631 was defined by an uncommitted script and
## a threshold that lived in a literal. This test applies the committed rule to
## the archived esummary set and requires the archived population back, exactly.
##
## It is deliberately independent of src/15c0_build_census_input.R: the script
## asserts its own output, and this asserts it again from the other direction,
## so a change to either is caught.
##
## Run: Rscript tests/test_population.R

suppressPackageStartupMessages({ library(testthat) })
source("src/utils/screening.R")

SUMM  <- "data/cache/rev2_summaries.rds"
CSV   <- "results/tables/audit_rev2_candidate_summaries.csv"
RDS   <- "data/cache/rev2_census_input.rds"
ATTR  <- "results/tables/l0_attrition.csv"
MAP   <- "results/tables/audit_candidate_map_rev2.csv"

skip_if_no_cache <- function() skip_if_not(file.exists(SUMM), "esummary cache absent")

test_that("the declared threshold is the one that was applied", {
  expect_equal(MIN_SAMPLES, 25)
  expect_true(SCREEN$exclude_single_cell)
  expect_true(SCREEN$exclude_array)
  expect_true(SCREEN$exclude_non_illumina)
})

test_that("the retrieval yields 16,820 unique series", {
  skip_if_not(file.exists(MAP), "candidate map absent")
  m <- read.csv(MAP, stringsAsFactors = FALSE)
  expect_equal(length(unique(m$accession)), 16820L)
})

test_that("applying the committed rule reproduces exactly 3,631 candidates", {
  skip_if_no_cache()
  summ <- do.call(rbind, readRDS(SUMM))
  scr  <- apply_screen(summ)
  expect_equal(sum(scr$candidate), 3631L)

  expect_true(file.exists(RDS))
  expect_setequal(scr$accession[scr$candidate], readRDS(RDS))
})

test_that("the archived summaries table carries the same flags the rule produces", {
  skip_if_no_cache()
  skip_if_not(file.exists(CSV), "summaries table absent")
  summ <- do.call(rbind, readRDS(SUMM))
  scr  <- apply_screen(summ)
  old  <- read.csv(CSV, stringsAsFactors = FALSE)
  old  <- old[match(scr$accession, old$accession), , drop = FALSE]
  for (f in c("is_series", "is_single_cell", "is_array", "non_illumina",
              "too_small", "candidate"))
    expect_identical(as.logical(old[[f]]), scr[[f]], info = f)
})

test_that("attrition is sequential and lands on the candidate count", {
  skip_if_not(file.exists(ATTR), "attrition table absent")
  a <- read.csv(ATTR, stringsAsFactors = FALSE)
  expect_equal(a$n_remaining[a$step == "identified"], 16820L)
  expect_equal(a$n_removed[a$step == "single_cell"], 2798L)
  expect_equal(a$n_removed[a$step == "array"], 174L)
  expect_equal(a$n_removed[a$step == "non_illumina"], 250L)
  expect_equal(a$n_removed[a$step == "too_small"], 9967L)
  expect_equal(tail(a$n_remaining, 1), 3631L)
  ## each step removes only survivors of the preceding ones
  expect_equal(a$n_remaining[-1], head(a$n_remaining, -1) - a$n_removed[-1])
})

test_that("the size threshold is the largest single exclusion", {
  skip_if_not(file.exists(ATTR), "attrition table absent")
  a <- read.csv(ATTR, stringsAsFactors = FALSE)
  rm <- a$n_removed[a$step != "identified"]
  expect_equal(which.max(rm), which(a$step[a$step != "identified"] == "too_small"))
  expect_gt(a$n_removed[a$step == "too_small"], sum(rm) - a$n_removed[a$step == "too_small"])
})

test_that("an unknown sample count is retained, not treated as small", {
  df <- data.frame(accession = c("GSE1", "GSE2", "GSE3"), entrytype = "GSE",
                   gdstype = "Expression profiling by high throughput sequencing",
                   title = "x", summary = "y",
                   n_samples = c(NA_integer_, 24L, 25L), stringsAsFactors = FALSE)
  s <- apply_screen(df)
  expect_identical(s$too_small, c(FALSE, TRUE, FALSE))
  expect_identical(s$candidate, c(TRUE, FALSE, TRUE))
})

cat("\ntests/test_population.R: all assertions passed\n")


==============================================================================
FILE: tests/test_retry.R
==============================================================================

## tests/test_retry.R -- retry loops must genuinely re-attempt (D-030).
##
## `force(expr)` cannot retry: in R a promise that has been forced and thrown is
## left "under evaluation", so subsequent force() calls raise "promise already
## under evaluation" instead of re-running the request. Every retry loop written
## that way does ONE real attempt then N phantom failures. These tests pin the
## corrected behaviour in all three loops.
source("src/utils/geo_pacer.R")
source("src/utils/pipelines.R")
suppressPackageStartupMessages(library(testthat))

test_that("geo_request re-evaluates and recovers from a transient failure", {
  n <- 0
  r <- geo_request({ n <- n + 1; if (n < 3) stop("transient") else "recovered" },
                   verbose = FALSE)
  expect_equal(r, "recovered")
  expect_equal(n, 3)                  # three GENUINE attempts, not phantoms
  expect_equal(geo_stats()$n_blocks, 0)   # must not declare a block
})

test_that("with_retry re-evaluates too", {
  n <- 0
  r <- with_retry({ n <- n + 1; if (n < 2) stop("transient") else "ok" },
                  attempts = 4L, base_wait = 1)
  expect_equal(r, "ok")
  expect_equal(n, 2)
})

test_that("the phantom-promise error never appears", {
  n <- 0
  msgs <- character(0)
  withCallingHandlers({
    r <- with_retry({ n <- n + 1; if (n < 3) stop("transient") else "ok" },
                    attempts = 4L, base_wait = 1)
  }, warning = function(w) { msgs <<- c(msgs, conditionMessage(w)); invokeRestart("muffleWarning") })
  expect_equal(n, 3)
  expect_false(any(grepl("promise already under evaluation", msgs)))
})

test_that("with_retry still gives up and returns NULL on persistent failure", {
  n <- 0
  r <- suppressWarnings(with_retry({ n <- n + 1; stop("hard") },
                                   attempts = 3L, base_wait = 1))
  expect_null(r)
  expect_equal(n, 3)                  # exactly `attempts` real tries
})

cat("\nAll retry tests passed.\n")

# ---- D-036: a bad key must not be mistaken for a rate block -----------------

test_that("repeated failure on ONE key gives up on that key, not a block", {
  .geo$consec_fail <- 0L; .geo$blocks <- list()
  .geo$last_key <- NA_character_; .geo$key_tries <- 0L
  n <- 0
  r <- suppressWarnings(geo_request(stop("bad series"), key = "GSE_BAD", verbose = FALSE))
  expect_null(r)
  expect_equal(geo_stats()$n_blocks, 0)      # crucially: NO block declared
})

test_that("failures across DISTINCT keys do declare a block", {
  .geo$consec_fail <- 0L; .geo$blocks <- list()
  .geo$last_key <- NA_character_; .geo$key_tries <- 0L
  assign("GEO_MAX_BLOCKS", 0L, envir = globalenv())   # give up at the first block
  for (k in c("GSE_A", "GSE_B", "GSE_C")) {
    suppressWarnings(geo_request(stop("throttled"), key = k, verbose = FALSE))
  }
  expect_gt(geo_stats()$n_blocks, 0)         # distinct keys => real block signal
})

test_that("a success resets both counters", {
  .geo$consec_fail <- 2L; .geo$last_key <- "GSE_X"; .geo$key_tries <- 1L
  r <- geo_request("fine", key = "GSE_Y", verbose = FALSE)
  expect_equal(r, "fine")
  expect_equal(.geo$consec_fail, 0L)
  expect_true(is.na(.geo$last_key))
})


==============================================================================
FILE: tests/test_three_valued.R
==============================================================================

## tests/test_three_valued.R -- an unresolved lookup is not an absence.
##
## D-045. `l4 = isTRUE(r$n_usable > 0) || ...` silently turned every lookup that
## could not be performed into a negative, because `isTRUE(NA > 0)` is FALSE.
## CLAUDE.md rule 3 forbids exactly this, and the Methods claims the opposite.
##
## Run: Rscript tests/test_three_valued.R

suppressPackageStartupMessages({ library(testthat) })

CENSUS <- "results/tables/l3_l5_census.csv"
RB     <- "results/tables/l3_route_b.csv"
XW     <- "results/tables/tierAB_crosswalk.csv"

test_that("the R idiom that caused the defect still behaves as described", {
  ## Not a test of our code -- a test that the diagnosis is right, so the entry
  ## in DEVIATIONS.md cannot rot.
  expect_false(isTRUE(NA > 0))
  expect_false(isTRUE(NA >= 0.9))
  expect_true(is.na(NA > 0))
})

test_that("the census emits three-valued status for every resource", {
  skip_if_not(file.exists(CENSUS), "census absent")
  d <- read.csv(CENSUS, stringsAsFactors = FALSE)
  for (nm in c("recount3_status", "dee2_status", "archs4_status")) {
    expect_true(nm %in% names(d), info = nm)
    expect_true(all(d[[nm]] %in% c("covered", "not_covered", "unassessable")), info = nm)
  }
})

test_that("every layer emits a lower and an upper bound", {
  skip_if_not(file.exists(CENSUS), "census absent")
  d <- read.csv(CENSUS, stringsAsFactors = FALSE)
  for (nm in c("l4_lower", "l4_upper", "l5_lower", "l5_upper", "n_unassessable"))
    expect_true(nm %in% names(d), info = nm)
  expect_true(all(d$l4_upper >= d$l4_lower))
  expect_true(all(d$l5_upper >= d$l5_lower))
  ## The point estimate the manuscript reports IS the lower bound.
  expect_identical(as.logical(d$l4), as.logical(d$l4_lower))
  expect_identical(as.logical(d$l5), as.logical(d$l5_lower))
})

test_that("upper exceeds lower exactly where a lookup was unassessable", {
  skip_if_not(file.exists(CENSUS), "census absent")
  d <- read.csv(CENSUS, stringsAsFactors = FALSE)
  gap <- d$l5_upper & !d$l5_lower
  expect_true(all(d$n_unassessable[gap] > 0))
  ## and a series with no unassessable lookup has no gap at either layer
  clean <- d$n_unassessable == 0
  expect_identical(d$l5_upper[clean], d$l5_lower[clean])
  expect_identical(d$l4_upper[clean], d$l4_lower[clean])
})

test_that("the bounds are non-trivial -- the defect had real magnitude", {
  skip_if_not(file.exists(CENSUS), "census absent")
  d <- read.csv(CENSUS, stringsAsFactors = FALSE)
  ## If this ever becomes zero, either every lookup now resolves (report it) or
  ## the three-valued machinery has been bypassed (fix it).
  expect_gt(sum(d$n_unassessable > 0), 0L)
  expect_gt(sum(!d$l4_lower & d$n_unassessable > 0), 0L)
})

test_that("L3 route B keeps 'no claim made' distinct from 'absent'", {
  skip_if_not(file.exists(RB), "route B absent")
  r <- read.csv(RB, stringsAsFactors = FALSE)
  expect_true(all(r$route_b %in% c("present", "absent", "no_status_line", "fetch_failed")))
  ## A record that renders no Raw data field made no claim; it is not an absence,
  ## and it must not be classified into L3a/L3b/L3c.
  ns <- r$route_b == "no_status_line"
  expect_true(all(r$l3_class[ns] == "L3_no_status_line"))
  expect_true(all(is.na(r$l3_agreed[ns])))
  ## A fetch failure is never an absence either.
  ff <- r$route_b == "fetch_failed"
  if (any(ff)) expect_true(all(is.na(r$l3_class[ff])))
})

test_that("L3a / L3b / L3c are only assigned on dual-route agreement", {
  skip_if_not(file.exists(RB), "route B absent")
  r <- read.csv(RB, stringsAsFactors = FALSE)
  cls <- r$l3_class %in% c("L3a", "L3b", "L3c")
  expect_true(all(r$route_a[cls] == "absent"))
  expect_true(all(r$route_b[cls] == "absent"))
  expect_true(all(r$l3_agreed[cls] %in% TRUE))
})

test_that("the crosswalk never counts an unmappable term as a disagreement", {
  skip_if_not(file.exists(XW), "crosswalk absent")
  x <- read.csv(XW, stringsAsFactors = FALSE)
  expect_true(any(grepl("^unresolved", x$tierB_descriptor)))
  ## unresolved verdicts carry no mapped descriptor to have disagreed with
  u <- grepl("^unresolved", x$tierB_descriptor)
  expect_true(all(x$tierB_descriptor[u] != "refuted"))
})

cat("\ntests/test_three_valued.R: all assertions passed\n")
