## Supplementary files 1-7 for "Most published human disease RNA-seq cannot be uniformly reanalyzed: a population-scale audit of reprocessability in the Gene Expression Omnibus": INDEX.txt

ADDITIONAL FILES — Reprocessability audit manuscript
Abdullah Bin Murad, Northeastern University
Source deposit: https://doi.org/10.5281/zenodo.21817531

AF1  Additional_file_1_per_disease_cascade.xlsx
     Per-disease cascade for all 1,124 enumerated MeSH descriptors, including the
     1,035 that yielded no qualifying series. 26 columns, one row per descriptor.

AF2  Additional_file_2_series_level_results.csv
     Series-level results for all 3,631 audited series, carrying the verbatim status
     string for every accession-level claim, three-valued per-resource coverage
     statuses, run-level coverage fractions, and both route-A and dual-route L3.

AF3  Additional_file_3_deviation_log.docx
     Full deviation log: 40 entries for the original run (D-026 never issued),
     including five defects found in the analysis code, plus 19 recorded during
     remediation. 59 entries total.

AF4  Additional_file_4_regression_test_suite.txt
     All 10 R regression test files reproduced in full, covering the screening rule,
     coverage bounds, three-valued handling, replication-counting basis, freeze
     integrity, deviation counts, eligibility filters, MeSH tree logic, determinism
     and retry behaviour.

AF5  Additional_file_5_case_definition_sensitivity.xlsx
     Sheet "summary": qualifying pairs, series and retained diseases under literal
     versus by-exclusion case assignment. Sheet "pair_level": all 1,835 pairs.

AF6  Additional_file_6_PRISMA_S_and_flow.xlsx
     Sheet "PRISMA-S 16 items": the search-reporting checklist mapped to the GEO /
     Entrez dataset search. Sheet "PRISMA 2020 flow": 17 boxes reconciling 16,820
     retrieved records to the 3,631 audited, with N/A items justified. Sheet
     "Search strategies": the query string for each of the 1,124 descriptors under
     both the original and revised retrieval templates.

AF7  Additional_file_7_ontology_crosswalk.xlsx
     Sheet "Concordance by policy": descriptor-level Tier A/B agreement under
     exact-only, exact-plus-broad, and exact-plus-broad-plus-cross-reference mapping
     policies. Sheet "Mapping coverage": per-term mapping outcome, predicate and route.
